# Quantitative Colour Pattern Analysis (QCPA): A Comprehensive Framework for the Analysis of Colour Patterns in Nature

**DOI:** 10.1101/592261

**Authors:** Cedric P. van den Berg, Jolyon Troscianko, John A. Endler, N. Justin Marshall, Karen L. Cheney

**Affiliations:** The School of Biological Sciences, The University of Queensland, Australia; Centre for Ecology & Conservation, Exeter University, UK; School of Life & Environmental Sciences, Deakin University; Queensland Brain Institute, The University of Queensland, Australia

**Author notes:** Joint first authors (Cedric P. van den Berg, Jolyon Troscianko).

**Keywords:** colour pattern analysis, colour perception, image analysis, visual modelling, receptor noise limited model, animal colouration, colour space

## Abstract

1. To understand the function of colour signals in nature, we require robust quantitative analytical frameworks to enable us to estimate how animal and plant colour patterns appear against their natural background as viewed by ecologically relevant species. Due to the quantitative limitations of existing methods, colour and pattern are rarely analysed in conjunction with one another, despite a large body of literature and decades of research on the importance of spatiochromatic colour pattern analyses. Furthermore, key physiological limitations of animal visual systems such as spatial acuity, spectral sensitivities, photoreceptor abundances and receptor noise levels are rarely considered together in colour pattern analyses.
2. Here, we present a novel analytical framework, called the ‘Quantitative Colour Pattern Analysis’ (QCPA). We have overcome many quantitative and qualitative limitations of existing colour pattern analyses by combining calibrated digital photography and visual modelling. We have integrated and updated existing spatiochromatic colour pattern analyses, including adjacency, visual contrast and boundary strength analysis, to be implemented using calibrated digital photography through the ‘Multispectral Image Analysis and Calibration’ (MICA) Toolbox.
3. This combination of calibrated photography and spatiochromatic colour pattern analyses is enabled by the inclusion of psychophysical colour and luminance discrimination thresholds for image segmentation, which we call ‘Receptor Noise Limited Clustering’, used here for the first time. Furthermore, QCPA provides a novel psycho-physiological approach to the modelling of spatial acuity using convolution in the spatial or frequency domains, followed by ‘Receptor Noise Limited Ranked Filtering’ to eliminate intermediate edge artefacts and recover sharp boundaries following smoothing. We also present a new type of colour pattern analysis, the ‘Local Edge Intensity Analysis’ (LEIA) as well as a range of novel psycho-physiological approaches to the visualisation of spatiochromatic data.
4. QCPA combines novel and existing pattern analysis frameworks into what we hope is a unified, user-friendly, free and open source toolbox and introduce a range of novel analytical and data-visualisation approaches. These analyses and tools have been seamlessly integrated into the MICA toolbox providing a dynamic and user-friendly workflow.
5. QCPA is a framework for the empirical investigation of key theories underlying the design, function and evolution of colour patterns in nature. We believe that it is compatible with, but more thorough than, other existing colour pattern analyses.

## Introduction

Animal colour patterns are complex traits which serve a multitude of purposes, including defence against predators (such as camouflage and aposematism), social signalling and thermoregulation (Cott, 1940). How colour patterns are perceived by animals is unique to a given visual system in a specific context. It depends on the visual background against which they are viewed, the visual capabilities of the signal receiver, the distance from which the pattern is viewed and the ambient light environment (Endler, 1978, 1990; Lythgoe, 1979; Merilaita *et al.*, 2001; Cuthill *et al.*, 2017). Animal visual systems are diverse, and vary in eye shape and size, visual pigment number and absorbance maximum, photoreceptor type and number, and retinal and post-retinal processing (Lythgoe, 1979; Cronin *et al.*, 2014). When determining the perception of colour patterns in other animals, it is therefore essential to consider spatial acuity (and viewing distance) as well as colour and luminance discrimination abilities (Endler, 1978). Humans have greater spatial acuity and contrast sensitivity than most vertebrates, except birds (da Silva Souza *et al.*, 2011; Caves *et al.*, 2016). We also have a different number of receptor classes, and different spectral sensitivity ranges compared to many animals (Cronin *et al.*, 2014). For example, most other mammals are dichromats (i.e. they have only 2 compared to our 3 cone types), while most birds, reptiles and some amphibians, spiders and fish possess an ultraviolet cone sensitivity and may be tetrachromats (Osorio & Vorobyev, 2005, 2008; Cronin & Bok, 2016). Among invertebrates the number of receptor classes may exceed 10 (Cronin *et al.*, 2014).

To examine the perception of colour signals by animals, studies generally measure colour, luminance and pattern characteristics (e.g. Marshall *et al.*, 2006; Cortesi & Cheney, 2010; Zylinski *et al.*, 2011; Allen & Higham, 2013; Xiao & Cuthill, 2016). For example, colour (chromatic) and luminance (achromatic) contrast is measured between colour patches within an animal, or between an animal and its background, and is calculated in terms of perceptual distances in colour space often using the Receptor Noise Limited Model (RNL) (Vorobyev & Osorio, 1998). This model assumes that the noise inside a given class of photoreceptors, in combination with their relative abundance and opponent colour processing mechanisms, are the fundamental limits of colour and luminance contrast perception. The relative stimulation of photoreceptors can then be used to map the perceptual distances between colour patches in colour space (reviewed by Renoult *et al.*, 2017). These Euclidian or geometric distances are expressed in terms of ΔS values (Vorobyev *et al.*, 2001; Siddiqi *et al.*, 2004). The model predicts that a ‘Just Noticeable Difference’ (JND) should be equivalent to ΔS = 1 if model conditions and assumptions are met (Vorobyev & Osorio, 1998). For quantifying the spatial properties of patterns, Fast Fourier Transform (FFT) analyses of pixel intensity in digital images (Switkes *et al.*, 1978), pixel or location dependent transition matrices (Endler, 2012) or landmark based pattern metrics are often used (Lowe, 1999; Troscianko *et al.*, 2017; Belleghem *et al.*, 2018).

These types of analyses aim to computationally reproduce the retinal processing of visual information, but often investigate colour, luminance or pattern contrast in isolation. For example, Cheney *et al.* (2014) quantified the conspicuousness of nudibranch molluscs (marine gastropods) by measuring pattern contrast against their natural backgrounds using FFT on digital images. They then measured chromatic contrast (ΔS) between animal and background using point measurements obtained by a spectrophotometer. While useful for many studies of animal colouration, these individual analyses ignore the interaction of visual information at various perceptual stages (for discussion see Stevens & Merilaita, 2011; Rowe, 2013; Endler & Mappes, 2017; Ng *et al.*, 2018; Ruxton *et al.*, 2018).

Spatiochromatic colour pattern analyses overcome these limitations as they are designed to consider perceptual interactions between spatial, chromatic and achromatic information (Endler, 1978, 1990; Vorobyev *et al.*, 2001; Endler & Mielke, 2005; Marshall *et al.*, 2006; Stevens & Merilaita, 2011; Endler & Mappes, 2017; Olsson *et al.*, 2017; Endler *et al.*, 2018; Ruxton *et al.*, 2018). Such approaches parameterise the properties of colour patterns, including colour adjacency, pattern regularity, visual contrast and colour pattern similarity (e.g. Endler & Mielke, 2005; Endler, 2012; Endler *et al.*, 2018). For example, not only is the efficiency of visual signals dependent on the presence or absence of colours, but also how those colours are arranged in patterns (Endler, 1984, 2012; Endler *et al.*, 2018; Green *et al.*, 2018; Troscianko *et al.*, 2018). The importance of this approach for understanding the design, function and evolution of visual signals has been highlighted repeatedly (e.g. Endler, 1978, 1984, 2012; Osorio *et al.*, 2004; Stevens & Merilaita, 2011; Endler *et al.*, 2018; Ruxton *et al.*, 2018).

Current methods for spatiochromatic colour pattern analysis (Endler, 2012; Endler *et al.*, 2018), which have recently been implemented by PAVO 2.0 (Maia *et al.*, 2018), are only suitable for processing colour patterns and visual scenes which have very clear colour differences (i.e. sharp boundaries with high chromatic and achromatic contrast), so that spectral data can be collected easily from each colour patch. Alternatively, these methods would require a prohibitively large number of spectral measurements to be made from a scene containing typical levels of natural variation; even the lowest acuity receivers would require many thousands of points to be measured. Digital imaging is therefore ideally suited to this type of analysis, because each image can rapidly and non-invasively capture millions of point samples which can provide the necessary chromatic and spatial information. However, currently available image segmentation and processing techniques do not incorporate physiological and cognitive limitations of ecologically relevant viewers. Indeed, many approaches rely on manually drawing the outlines of colour pattern elements by a human observer or clustering algorithms using uninterpreted RGB information inside a digital image (e.g. Endler & Houde, 1995; Isaac & Gregory, 2013; Allen & Higham, 2015; Winters *et al.*, 2017). Such approaches are prone to introduce anthropocentric (qualitative) as well as quantitative bias in interpreting the design, function and evolution of animal colouration, unless the colours fall in clear classes and they have been checked and calibrated with a spectrometer or calibrated digital photography.

In this paper, we introduce a method to overcome these problems and present a user-friendly, open-source framework, which we call ‘Quantitative Colour Pattern Analysis’ (QCPA). QCPA is a comprehensive approach to the study of the design and function of colour patterns in nature. It combines calibrated digital photography (Stevens *et al.*, 2007), visual modelling and colour pattern analysis into an analytical framework that is seamlessly integrated into the ‘Multispectral Image Analysis and Calibration Toolbox’ (MICA) (Troscianko & Stevens, 2015). QCPA enables the use of existing, revised and newly developed colour pattern analyses on an unprecedented quantitative and qualitative scale. This is enabled by image segmentation using combined colour and luminance discrimination thresholds (RNL Clustering) as well as improved modelling of visual acuity (RNL Ranked Filtering). Pattern analyses included in QCPA are colour adjacency analysis, visual contrast analysis and boundary strength analysis (Endler & Mielke, 2005; Endler, 2012; Endler *et al.*, 2018), which we have expanded, adapted and revised. For example, we include local edge intensity analysis, a novel extension to boundary strength analysis (Endler *et al.*, 2018), which allows for colour pattern edge intensity analysis approximating the scale of receptive fields of a visual system while not requiring a segmented image. QCPA provides the user with a freely adjustable network of image processing tools which can convert visual information into a highly descriptive array of numbers and representative figures which may then be used to examine a variety of evolutionary, behavioural and ecological questions (Fig. 1). Potential applications of QCPA include (but are not limited to): background matching, disruptive colouration, polymorphism, mimicry, aposematism, sexual signalling, territorial signalling, thermoregulation and landscape analysis.

**Figure 1:**
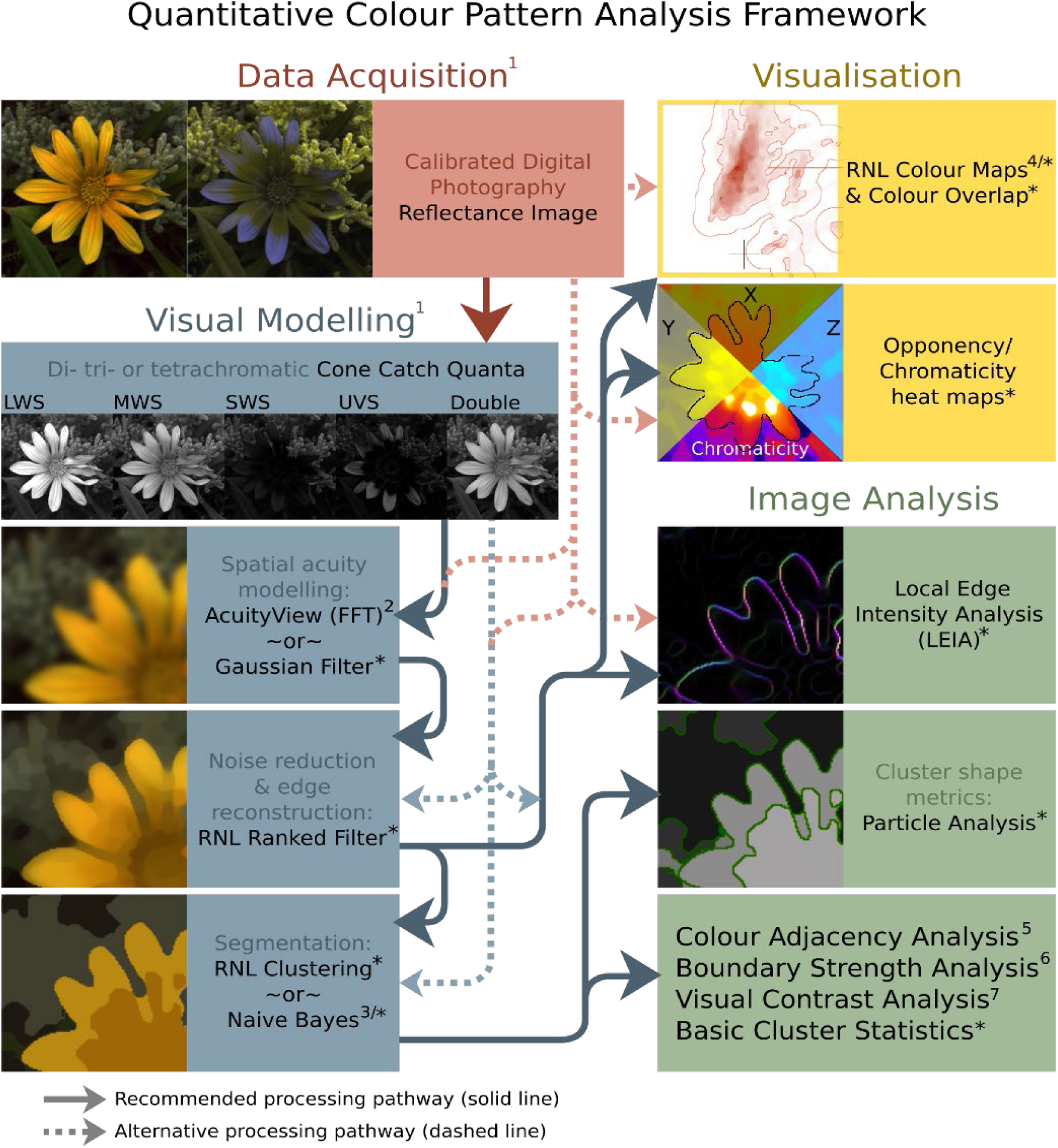
Schematic of the ‘Quantitative Colour Pattern Analysis’ QCPA framework. Asterisks (*) show steps in the framework which are novel, or have been heavily adapted for use in this framework, while numbers refer to existing techniques. Cone-catch images are the input into the framework, which can be generated with the MICA toolbox (^1^ Troscianko & Stevens 2015). Spatial acuity modelling is then used to remove visual information which would not be visible given the acuity and viewing distance (using either AcuityView 2.0, Caves & Johnsen 2017, or a Gaussian convolution-based approach*). Acuity correction generates blurred images with intermediate colours that are not likely to be perceived by the receiver. The RNL ranked filter* is therefore used to recreate sharp boundaries. These images are ideal input for the local edge intensity analysis (LEIA)*, and for generating colour maps in RNL chromaticity space (*/4, Hempel De Ibarra *et al*., 2002; Kelber *et al.*, 2003; Renoult *et al.,* 2017). RNL clustering* or Naive Bayes clustering (*/3, Koleček *et al.*, 2018) are then used to segment the image prior to colour adjacency analysis (5, Endler 2012), boundary strength analysis (6, Endler *et al.* 2018), visual contrast analysis (7, Endler 1991; Endler & Mielke 2005), and particle shape analysis*.

## Materials and Methods

We first provide a brief description of the acquisition of calibrated digital images and theoretical visual modelling of the viewer and then describe individual tools of the QCPA in more detail, including:

- **Modelling of spatial acuity**: using an adaptation of Fast Fourier transform or Gaussian filters;
- **Image smoothing and edge reconstruction**: using the receptor noise limited ranked filter;
- **Image segmentation**: using receptor noise limited clustering and naïve Bayes clustering;
- **Pattern analysis:** using adjacency, boundary strength, visual contrast analysis, local edge intensity analysis and particle analysis;
- **Data visualisation:** using ΔS edge intensity images, XYZ opponency images, RNL saturation images and colour maps

Finally, we describe how the rich numerical output of QCPA can be used to investigate complex interactions between output parameters, and how these can be used to investigate the design, function and evolution of colour patterns in nature. We also provide additional technical details and worked examples in the Supplementary Information.

### Step 1: Acquisition of calibrated digital images

Acquiring data suitable for analysing the spatiochromatic properties of a scene is the first requirement for implementing the QCPA. The open-source and user-friendly MICA toolbox can be used to generate calibrated multispectral images and cone-catch images from almost any digital camera (Troscianko & Stevens, 2015). Cone-catch images model the photoreceptor stimulation of an animal for every single pixel within an image, with additional support for ultraviolet (UV)-sensitive cameras when modelling the vision of species with UV sensitivity (Fig. 1 & 2) (Troscianko & Stevens, 2015). While hyperspectral cameras are, theoretically, also well-suited to this task (e.g. Long & Sweet, 2006; Russell & Dierssen, 2015), there are a number of limitations in their use including cost and image resolution. However, the QCPA framework can also be used for the analysis of hyperspectral images. Precise instructions on how to obtain high quality calibrated image data are outlined in Troscianko & Stevens (2015).

**Figure 2:**
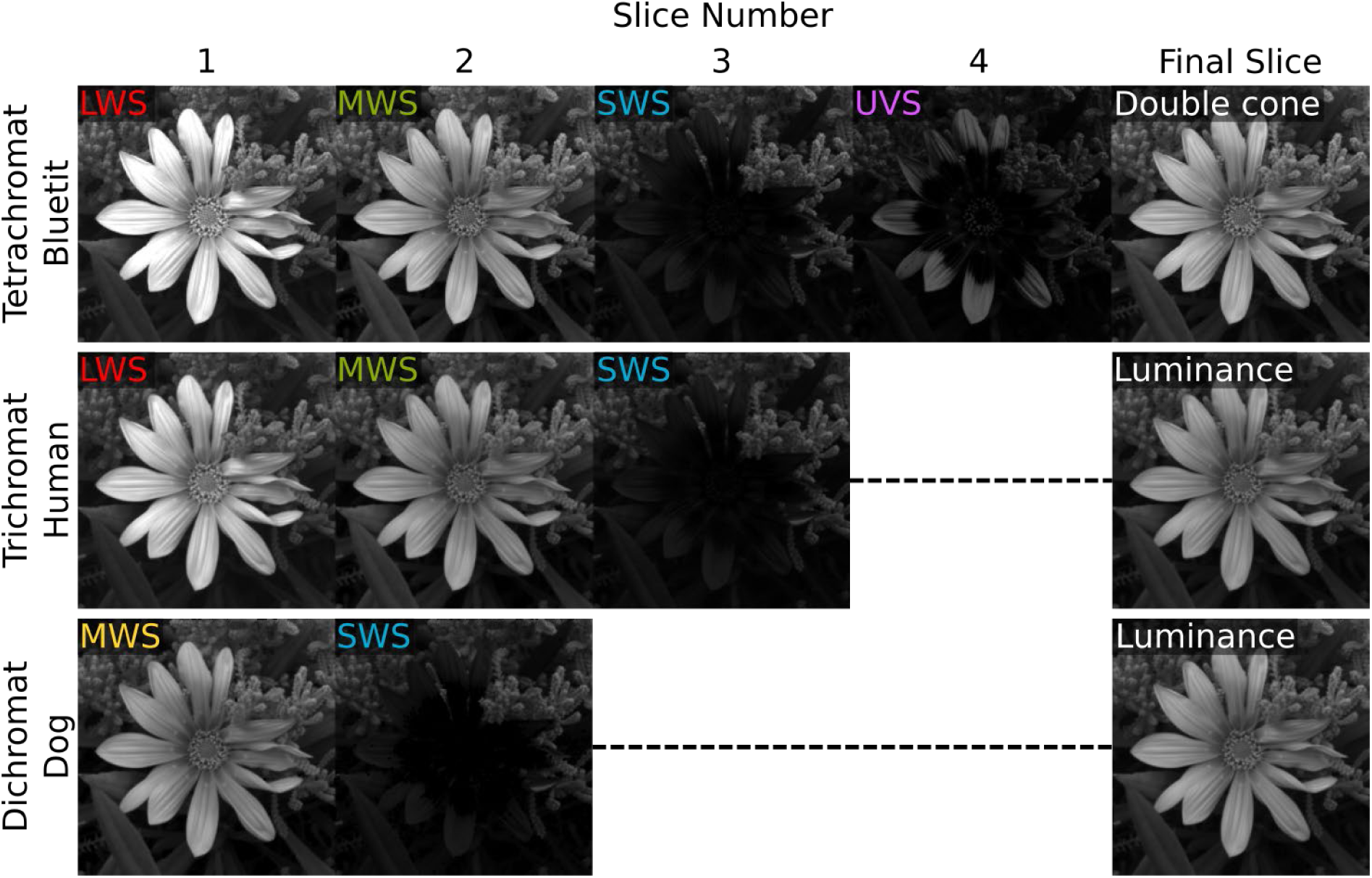
Example of multispectral image stacks as an output of MICA. Note that each image stack has a designated luminance channel layer. This is needed for QCPA to allow inferences based on luminance discrimination thresholds.

The MICA toolbox provides its own growing set of image analysis tools (e.g. Troscianko *et al.*, 2017) to which the QCPA contributes. Importantly, MICA allows the user to model cone captures in response to any possible light environment. This is very useful, as it allows to observe a visual scene in one light environment (e.g. a flower in a field at noon on a cloudy day) and translate the scene to another light environment (e.g. the same flower but under a long-wavelength enriched clear-sky sunrise light spectrum). MICA also lets the user switch between spectral sensitivities and cone channels of different species if that information is available (e.g. the same flower looked at by a bee in comparison to a bird). This function of MICA is increasingly being used by a range of researchers to introduce animal colour vision to their colour pattern studies (e.g. Chan *et al.*, 2019). Species-specific information on spectral sensitivities is often hard to obtain. However, in many cases it is possible to overcome this by estimating spectral sensitivities using information from closely-related species (Kemp *et al.*, 2015; Olsson *et al.*, 2017).

### Step 2: Defining discrimination thresholds

The chromatic (ΔS_C_) and achromatic contrast (ΔS_L_) within an image can be calculated in terms of perceptual distance between any two pixels in 1 to *n*-dimensional colour space (Clark *et al.*, 2017) using the RNL model (Vorobyev & Osorio, 1998) (as per equations in Renoult *et al.*, 2017 and Gawryszewski 2018) for chromatic contrast, and Siddiqi *et al.*, 2004 for achromatic contrast). In its current state, QCPA uses the RNL equations for bright light (photopic) conditions. However, to calculate photoreceptor stimulation in dim light conditions, we advise the use of the dim light corrected receptor noise equations or, if necessary, rod based estimates (Vorobyev & Osorio, 1998; Veilleux & Cummings, 2012). These contrast values can then be used to remove pixel noise (fluctuations in pixel intensity due to noise in the camera sensor) from a digital image, as well as for its segmentation into colour patterns. Species specific data on visual systems (particularly receptor noise) can be difficult to obtain. This often results in model parameters being estimated. In combination with deviations from assumptions of the RNL model (Vorobyev & Osorio, 1998) this emphasizes the need to validate discrimination thresholds using behavioural experiments or choosing conservative thresholds (for discussion see Olsson *et al.*, 2017).

### Step 3: Modelling of spatial acuity

The ability of an animal to resolve patterns depends on the spatial acuity of its vision, which may be determined through anatomical, behavioural or physiological measurements (e.g. Champ *et al.*, 2014), in addition to the distance at which the objects is viewed. To understand why animals display particular colour patterns, it is important to investigate whether or not a colour pattern element is visible to an animal from a certain distance (Endler, 1978; Marshall, 2000). For example, a worker bee does not perceive the intricate UV patterns of a flower that guide the bee to its nectar storage until it is close due to the limitations of its visual acuity (Fig. 3). QCPA adapts and expands upon existing tools for modelling spatial acuity by using an adaptation of AcuityView (Caves & Johnsen, 2017) and Gaussian filter mediated blurring.

### Step 4: Eliminating problems in acuity-related processing using the RNL Ranked Filter

As noted by Caves & Johnsen (2017), the blurring of images to model visual acuity (Step 3) is not intended to manipulate images to represent how the scene would be perceived by the receiver; instead, it eliminates details which the specific visual system cannot resolve (Caves & Johnsen, 2017). It is likely, just like a bee looking at a field of flowers from 1m distance, that many animals perceive clearly delineated spatial information as the available visual information is integrated in retinal or post-retinal processing. Furthermore, blurred edges are problematic for clustering techniques or boundary comparison techniques and may create further artefacts of processing that are likely irrelevant to the animal. Pixel noise fluctuation in the sensor of a digital camera can also interfere with the clustering process, creating false edges, artificial colour pattern elements or influencing edge structure of colour pattern elements.

To overcome these issues and recreate sharp edges and mitigate pixel noise, we have developed an RNL mediated filter that can be applied to an image prior to clustering, which we call the ‘RNL Ranked Filter’. Subjectively, the filter resembles the ‘Smart Blur’ used in photo editing software packages (such as the ‘Adobe Creative Cloud’) and other rank selection filters, which rank the pixels in a kernel and modify the pixels based on that ranking. However, our custom written algorithm uses an estimate of an animal’s psychophysical ability (Using the RNL model) to discriminate between colours and luminance to recreate sharp edges and reduce pixel noise in a cone catch image (Fig. 4c).

**Figure 3:**
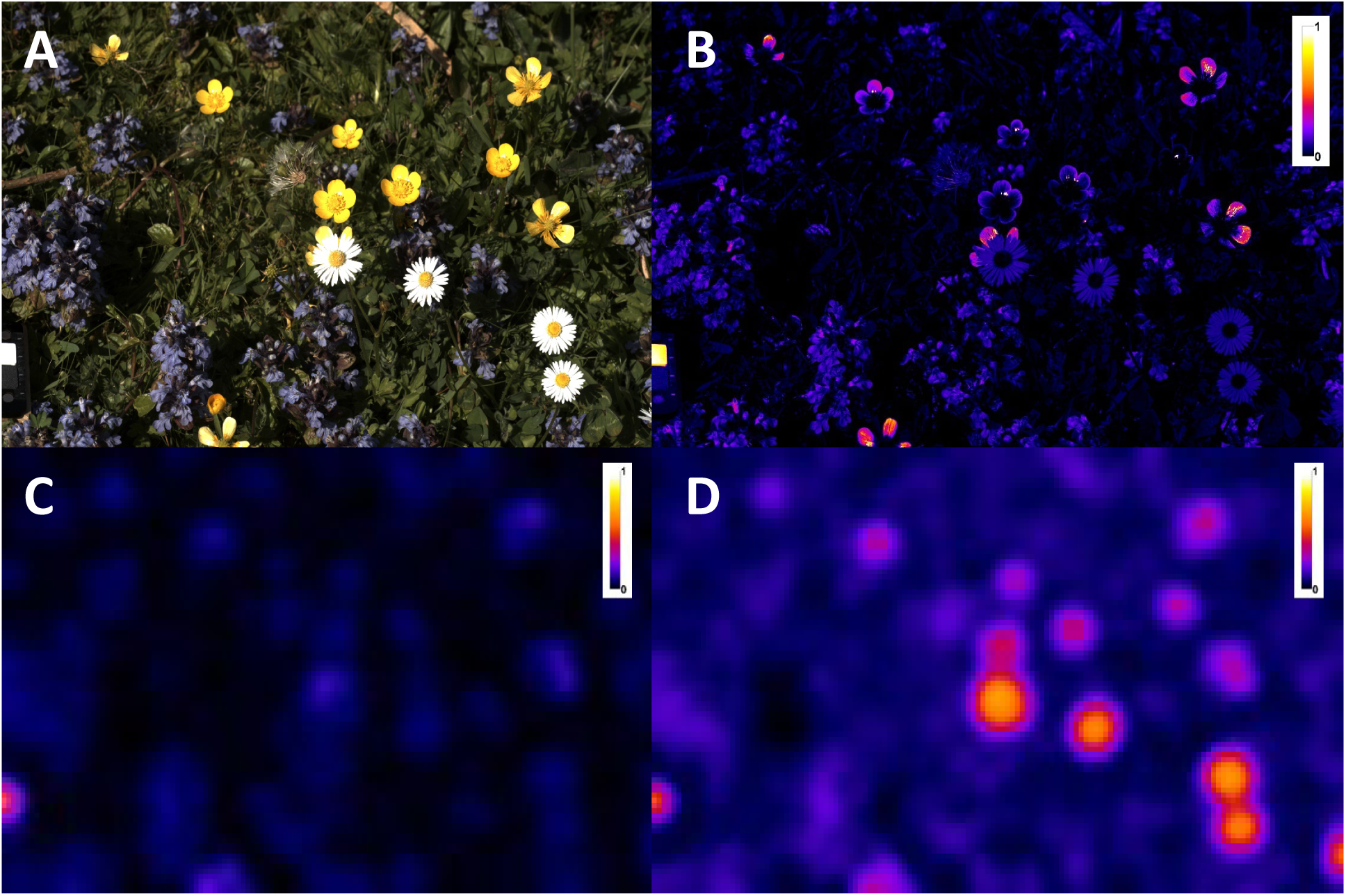
A) A flower meadow as seen by a human observer. B: UV intensity as detected by a worker bee with superior spatial acuity, which may lead to the false assumption of the UV information being available to the bee from a distance. C) UV intensity as detected by a worker bee with a spatial acuity of 0.5 cycles/degree at 1m distance. D) Medium-wavelength sensitive photoreceptor stimulation (used for luminance detection) of a worker bee at 1m distance. Note: the white standard (bottom left) remains detectable in all pictures. The scale in the top right corner of each image shows the relative stimulation of the given receptor channel.

**Figure 4:**
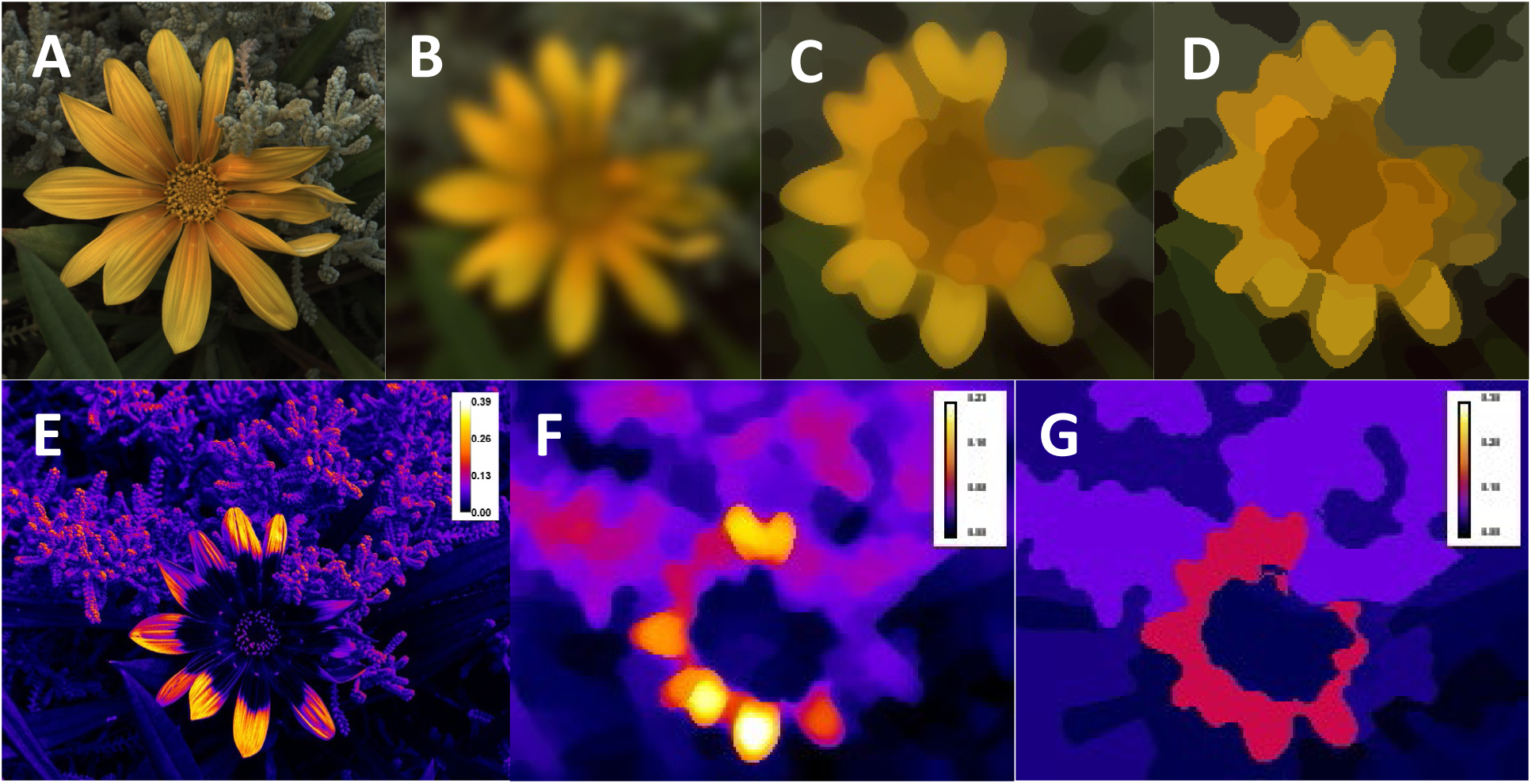
A) A reconstructed RGB image of a daisy using cone stimulation of the short, medium and long-wavelength sensitive photoreceptor channels of a blue tit (*Cyanistes caeruleus*). The UV photoreceptor is not shown for simplicity. B) The image after FFT filtering using a spatial acuity of 4.8 cycles/degree and a viewing distance of 2m. C) Recreation of sharp edges using the RNL ranked filter. D) Clustering the image into colour pattern elements using the RNL clustering. C & D assume a conservative receptor noise of 0.05 and a cone ratio of 1:2:3:3 (Hart et al., 2000). D uses a colour discrimination threshold of 3 ΔS and a luminance discrimination threshold of 4 ΔS. See Step 1 for details on multispectral imaging. E: UV information without acuity modelling as perceived by a worker bee (*Apis melifera*) F: Acuity and RNL ranked filtered as per 15cm viewing distance G: RNL clustered UV layer using a chromatic threshold of 2 ΔS and an achromatic threshold of 4 ΔS. The scale on the top right of the images indicates the stimulation of the uv receptor channel.

### Step 5: Psychophysical image segmentation using RNL Clustering

While a range of pattern analyses, including granularity analysis (Stoddard & Stevens, 2010) or NaturePatternMatch (Stoddard *et al.*, 2014) can be applied to an unsegmented picture (Steps 1-4), other pattern analyses, such as Patternize (Belleghem *et al.*, 2018) or those in PAVO 2.0 (Maia *et al.*, 2018) require an image segmented into colour pattern elements. However, image segmentation is often created subjectively using human perception; for example, a researcher estimating how many colour elements there are within a pattern. This may be sufficient for simple patterns but is likely to introduce significant anthropocentric bias when analysing complex patterns and when the visual system of the animal differs dramatically from a human visual system. Here, we present an agglomerative hierarchical clustering approach which uses colour and luminance discrimination thresholds of an animal, either in combination with each other or separately. By comparing each pixel to its neighbouring pixels, we can use the log-transformed RNL model to determine whether any two pixels could be discriminated based on colour and/or luminance contrast perceived by an animal. Once completed across an entire sample, this process results in an image that is segmented according to an animal’s psychophysiological discrimination thresholds (Fig. 4d).

### Step 6: Colour pattern analysis

At this point of the QCPA workflow (Fig. 1), the user has an image which has been filtered and modified according to the physiological and psychophysical limitations of an animal visual system, in the context of the physical environment. We can now quantify this information to investigate questions on the design and function of a colour pattern.

In this section, we present a range of secondary image statistics that can be derived from un-clustered, filtered as well as clustered images. We have, for this purpose, adapted and interpreted analytical frameworks such as adjacency analysis (Endler, 2012), visual contrast analysis (Endler, 1991; Endler & Mielke, 2005) as well as boundary strength analysis (Endler *et al.*, 2018). We also present new parameters and alternative outputs of these frameworks, new types of pattern analyses as well as various ways of visualising and plotting image and pattern properties (Table 1, Fig. 1).

**Table 1.**
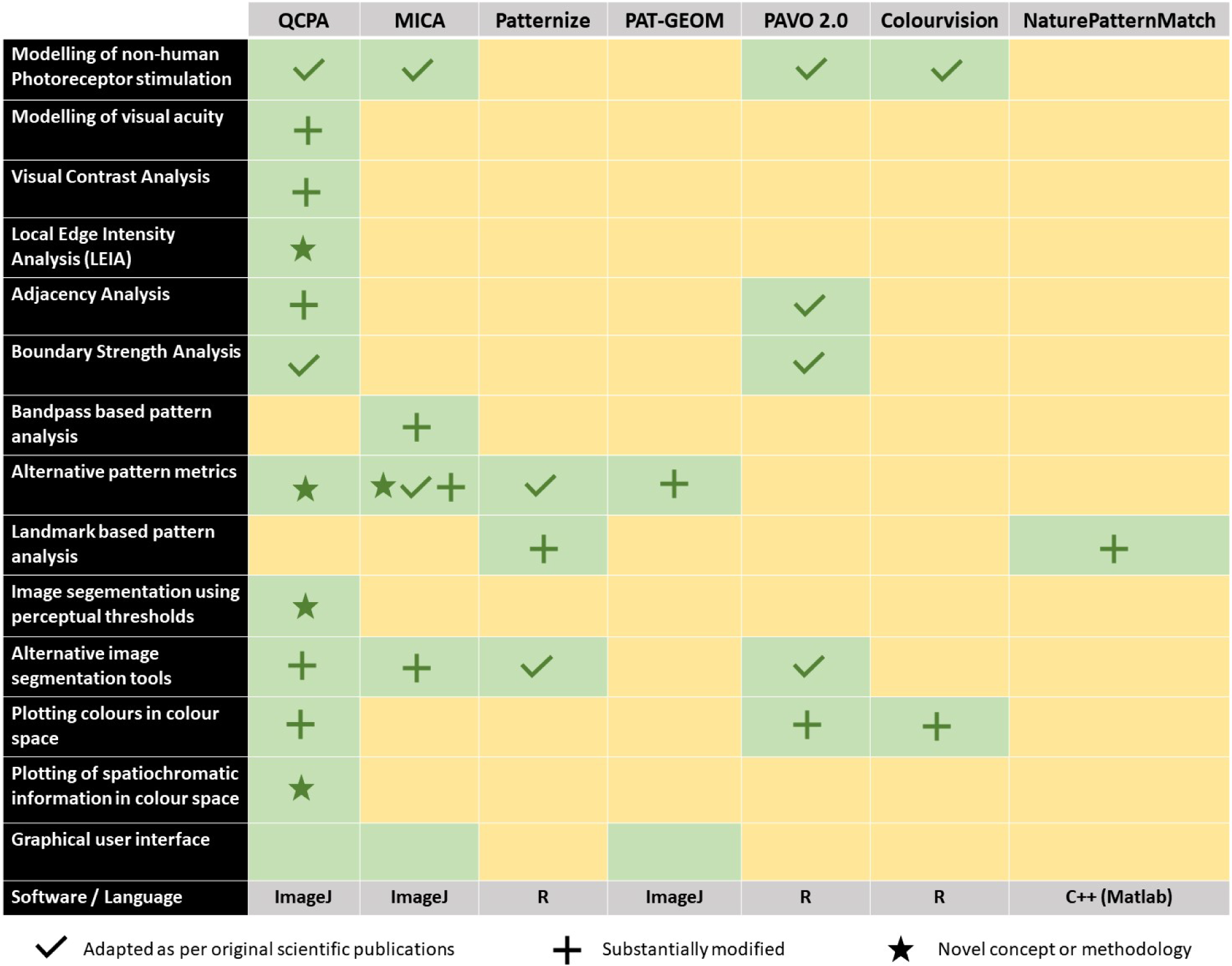
A comparison of the QCPA framework to other existing pattern analyses and frameworks. For patternize see (Belleghem *et al.*, 2018). For PAT-GEOM see Chan *et al.* (2018). For PAVO see Maia *et al.* (2019). For NaturePatternMatch see Stoddard *et al.* (2014). For Colourvision see Gawryszewski (2018). We would also like to point out an approach by Pike (2018) which operates on a similar principle to NaturePatternMatch.

#### 6.1: Adjacency Analysis

Adjacency analysis provides an approach for measuring the geometric properties of colour patterns and entire visual scenes (Endler, 2012). The concept is based on measuring the frequencies of transitions along transects across an image parallel and perpendicular to an animal’s body axis, or at random as in ecological studies of habitats. The information is captured in a transition matrix which can then be used to derive relevant pattern parameters relative to pattern geometry and potential function. In addition to providing frequently used metrics to describe patterns (e.g. aspect ratios, regularity, entropy, patch size), adjacency analysis allows quantification of pattern elements and their neighbours (Fig. 5). Colour adjacency analysis can be used for the quantification of mimicry and colour pattern polymorphism as well as colour pattern complexity. For example, in many cases of mimicry, the mimic only replicates the presence or absence of specific model colours in their patterning, without matching the model’s spatial arrangement. To a human observer this imperfect mimicry might be immediately apparent, while the intended receiver is unable to distinguish between the model and mimic (Fig 6) (Mallet & Joron, 1999; Endler, 2012; Dalziell & Welbergen, 2016). In a hypothetical case, colour adjacency analysis could be used to investigate imperfect sexual mimicry of orchids (Fig. 6) where the plant tries to mimic both the visual and chemical appearance of a potential mate (e.g. Gaskett & Herberstein, 2010). For further discussion of the biological relevance and application of this approach see Endler (2012), Rojas *et al.* (2014) and Winters *et al.* (2018).

**Figure 5:**
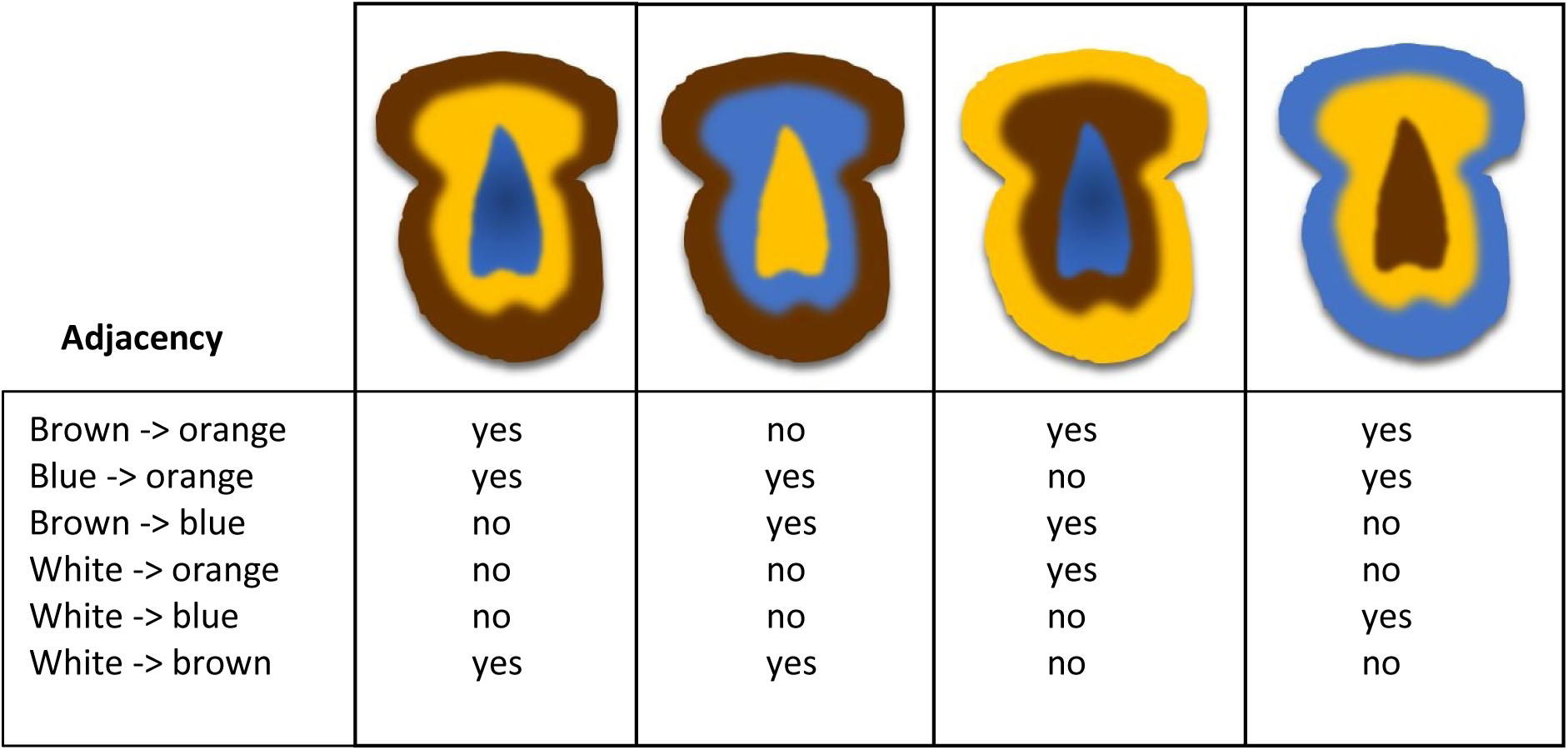
Four hypothetical colour pattern variations in the central part of an orchid flower aiming to mimic the female of an insect pollinator. Adjacency analysis can be used to discriminate between all of them. Other measures of pattern, such as complexity, would remain identical between the variants.

**Figure 6:**
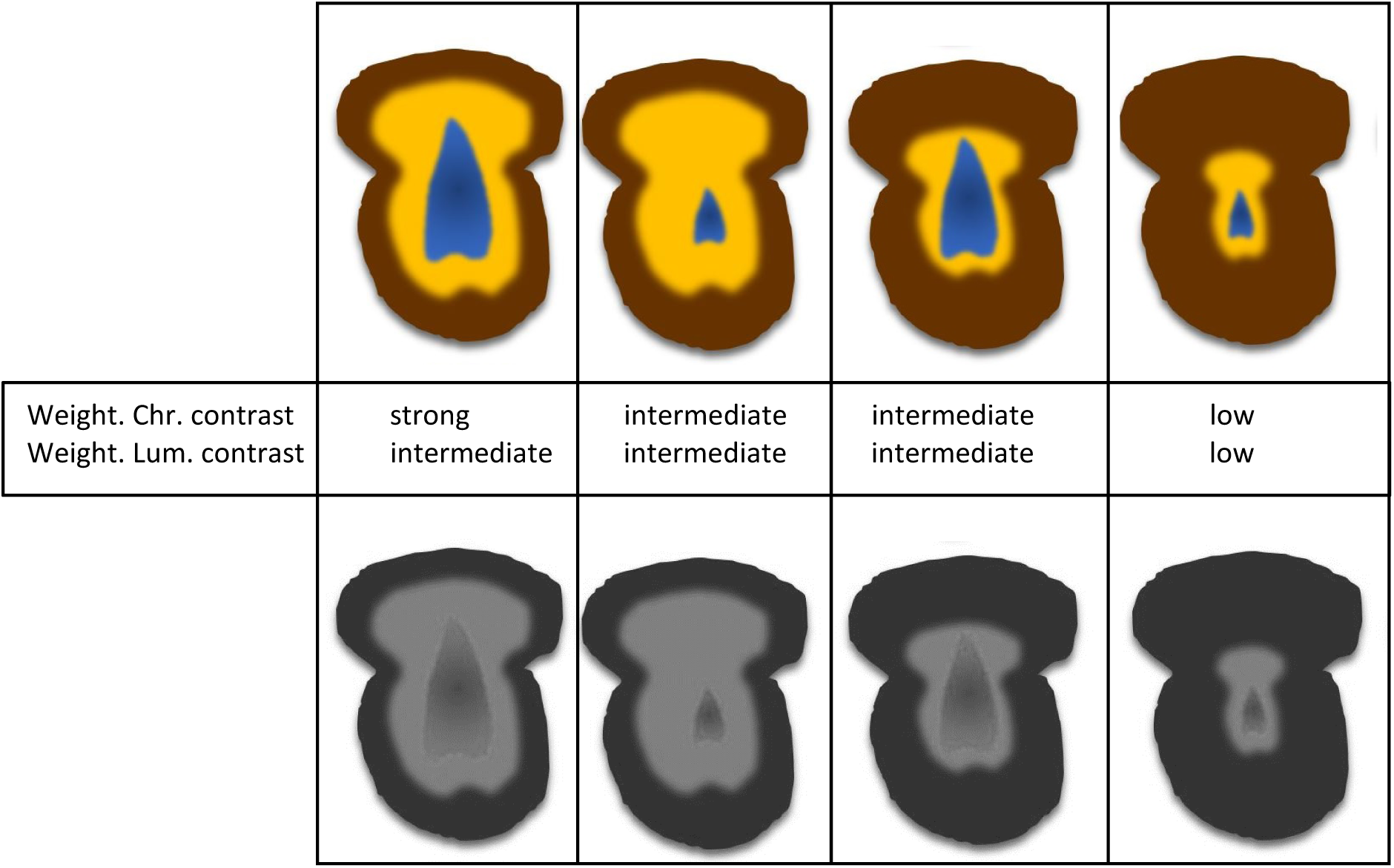
A hypothetical example of four pattern variations found in an orchid species. Visual contrast analysis can be used to quantify the combined effect of the colour pattern element relative abundances in combination with the reflective properties of the pattern elements. In this case, morph A has a strong abundance weighted chromatic and luminance contrast as opposed to morph D.

#### 6.2: Visual Contrast Analysis

Colour adjacency analysis focuses on pattern geometry, whereas visual contrast analysis is designed to investigate colour, pattern and luminance simultaneously (Endler & Mielke, 2005). This is important because spatial, chromatic and achromatic properties of colour patterns and visual scenes can interact to promote or suppress functional components of pattern conspicuousness such as its saliency, vividness, memorability and detectability (e.g. Endler & Houde, 1995; Green *et al.*, 2018). The perception of visual contrast is combination of the spatial (relative size and position of colour pattern elements), chromatic (hue and saturation) and achromatic (luminance) properties of a colour pattern due to low and higher level neuronal processing of visual information (e.g. Pearson & Kingdom, 2002; Simmons & Kingdom, 2002; Willis & Anderson, 2002; White *et al.*, 2017). We have adapted some of these metrics to use known or assumed colour opponency mechanisms to measure chromaticity (Hempel De Ibarra *et al.*, 2002). The interaction between the absolute and relative size of colour pattern elements and their chromatic and achromatic properties includes simultaneous colour contrast and colour constancy mechanisms that are understood in very few visual systems (e.g. Simpson *et al.*, 2016). The visual contrast analysis provides a set of metrics that are designed to be capable of capturing some of these effects. Using the previous orchid example, this analysis could be used to investigate how polymorphism in our hypothetical population interacts with pollinator learning and flower detectability (Fig. 6).

#### 6.3: Boundary Strength Analysis

Boundary strength analysis (BSA, Endler *et al.*, 2018) is an extension of the adjacency analysis (Endler, 2012). The transition matrices generated in the process of adjacency analysis can be used to measure the properties of boundaries between colour pattern elements. The underlying argument for this type of analysis is that the relative size, abundance, colour, brightness and adjacency of the patches within a colour pattern, and the intensity (ΔS) of the boundaries between adjacent patches, influence its signalling properties (e.g. Green *et al.*, 2018). These parameters also define the properties of the edges between and within parts of visual scenes and textures. On top of investigating detectability, saliency and polymorphism, BSA and the adjacency and visual contrast analysis are also capable of quantifying possible effects of viewer perspective and movement (Endler *et al.*, 2018). For example, in the two previous orchid examples, the geometry of the flower colour pattern changes (Fig. 5 & 6) but neither the adjacency, nor the visual contrast analysis quantify how these changes might correlate with the perception of the edges by a pollinator or what that could mean for the signal function.

#### 6.4: Local Edge Intensity Analysis (LEIA) and ΔS Edge Maps

Boundary strength analysis (BSA) depends on a segmented image with clearly delineated (clustered) colour pattern elements (Endler *et al.*, 2018). However, the segmentation process removes a large degree of subthreshold information, particularly smooth gradients of brightness and colour which the viewer may perceive. For this purpose, we provide ‘Local Edge Intensity Analysis’ (LEIA), as a way of quantifying edge properties in an image or ROI (Region of interest) that does not rely on a segmented input. By comparing each pixel to its horizontal, vertical and diagonal neighbours LEIA quantifies edge intensities in terms of colour and luminance contrast in log-linear RNL opponent space (Renoult *et al.*, 2017). The result can be visualised as ‘ΔS Edge Images’ (Fig. 7). While the BSA weights the strength of boundary classes according to their global (across an entire image or ROI) relative abundance, LEIA provides a local measurement of edge intensity on roughly the scale of an edge detecting receptive field. While LEIA is suited to the investigation of similar aspects of colour pattern design and function as BSA, it can do this without the need for clustering an image, while using a more neurophysiological approach than BSA. However, both have their own advantages and limitations. We recommend that LEIA should be used on images which have first been controlled for acuity (to remove imperceptible edge/gradient information) and images which have also been through the RNL ranked filter, so that local chromatic and luminance edges have been reconstructed to their maximal values. LEIA also provides numerical output describing the distribution of edge intensities across an image. These parameters are specifically designed to be robust in the case of non-normally distributed edge intensities in an image (e.g. a small conspicuous object on a homogeneous background). Local edge contrast can be visualised as ΔS edge intensity images (Fig. 7 b & c).

**Figure 7:**
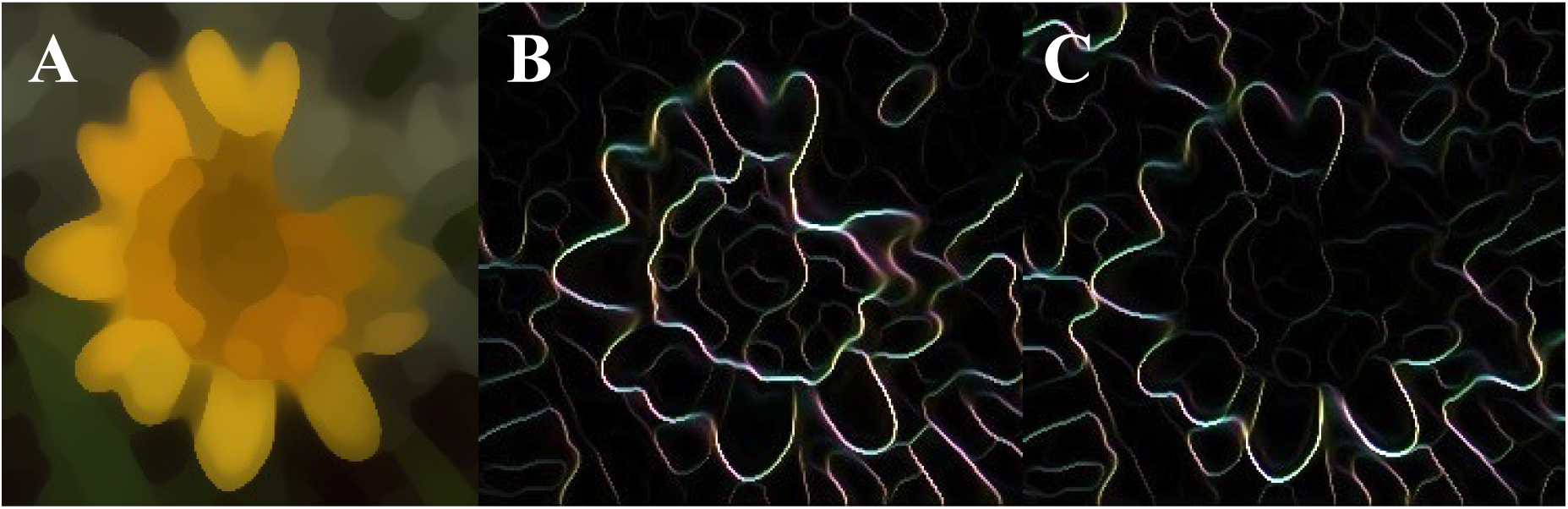
A) The RNL filtered flower from fig. 5c. B) Edge intensities of chromatic ΔS contrast. Different colours indicate different angles of edge detectors, the intensity reflects the contrast. C) Edge intensities of achromatic (luminance) ΔS contrast. Colours show edge angle whereas intensity shows edge strength.

### Step 7: Data visualisation

We provide a range of novel approaches for data visualisation. Calibrated digital photography and the coupled transformation of image data into psychophysical colour spaces provides a challenge but also an opportunity for visualisation. We have already introduced the ΔS edge intensity images and extend that selection with colour maps, XYZ opponency images and saturation images.

#### 7.1 ‘Colour Maps’ and ‘XYZ Opponency & Saturation Images’

The representation of chromatic information in colour spaces is a useful tool for data visualisation in visual ecology (Maia *et al.*, 2013; Renoult *et al.*, 2017; Gawryszewski, 2018). To date, most studies present their data as a scattering of points, which are either discrete measurements taken with spectrometers, or the mean centroids of image ROI cone-catch values. Techniques such as area or volume overlap between point clouds, or permutation analysis are then used to determine how dissimilar two colour patches are (e.g. Endler & Mielke, 2005; Kemp *et al.*, 2015; Maia & White, 2018).

Colour space data visualisations generally do not incorporate any spatial (colour pattern) information. The use of calibrated digital imaging provides thousands, or even millions of colour measurements within each ROI, capturing the entire range of chromatic gradients present in any natural pattern. Using the log transformed opponent colour space (Hempel De Ibarra *et al.*, 2002; Kelber *et al.*, 2003; Renoult *et al.*, 2017) we provide representations of spatiochromatic information in a perceptually calibrated colour space. ‘Colour Maps’ allow for the representation and delineation of entire visual scenes in a psychometric colour space, incorporating information on the chromaticity and saturation of each pixel in addition to the abundance of colours across part of the image (Fig. 8). Among other purposes, colour maps may be used for visualisations and investigations of chromatic background matching. The degree to which segments of an image overlap in colour space can be expressed as a percentage of overlap weighted by abundance. QCPA integrates tools which enable colour maps to be flexibly combined and compared between image sections, or measurements taken from a large dataset of images.

**Figure 8:**
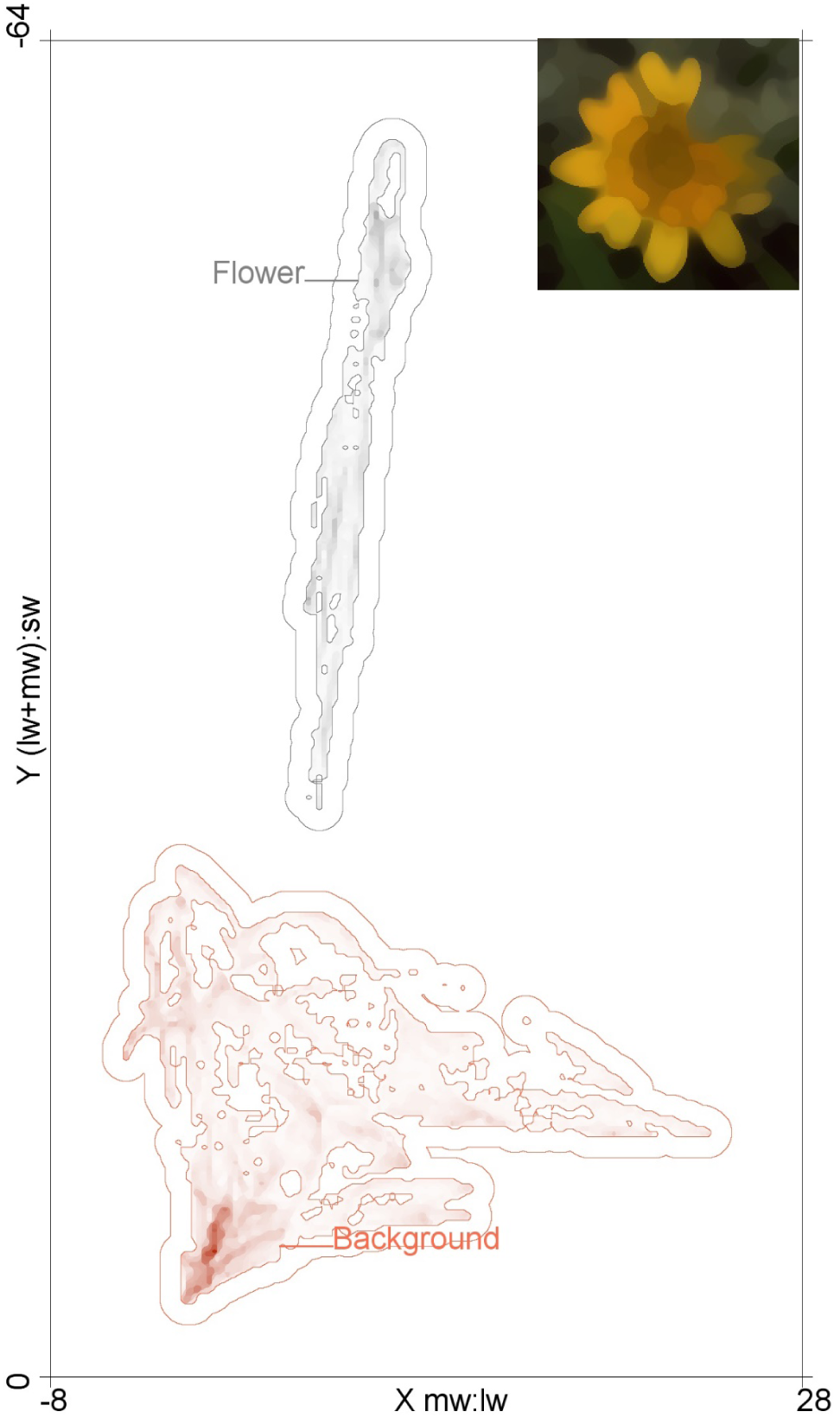
A Colour Map of the non-UV information in figure 5c. The boundary around each ROI pixel cloud reflects 1 ΔS. Darker parts of the cloud indicate more pixels in that ROI are located at that coordinate in the log-transformed RNL colour space. In this case, the flower and its background do not overlap (0%). For tetra-chromatic colour maps the Z-axis is represented as a stack of X&Y maps.

We also introduce the ability to convert cone-catch images to RNL XYZ chromaticity images, which allow visualisation and measurement of the independent axes of colour in a di-tri-or tetra-chromatic image, in addition to generating a saturation image (showing the Euclidian distance of each pixel’s RNL XYZ chromaticity values to the achromatic point) (Fig. 9).

**Figure 9:**
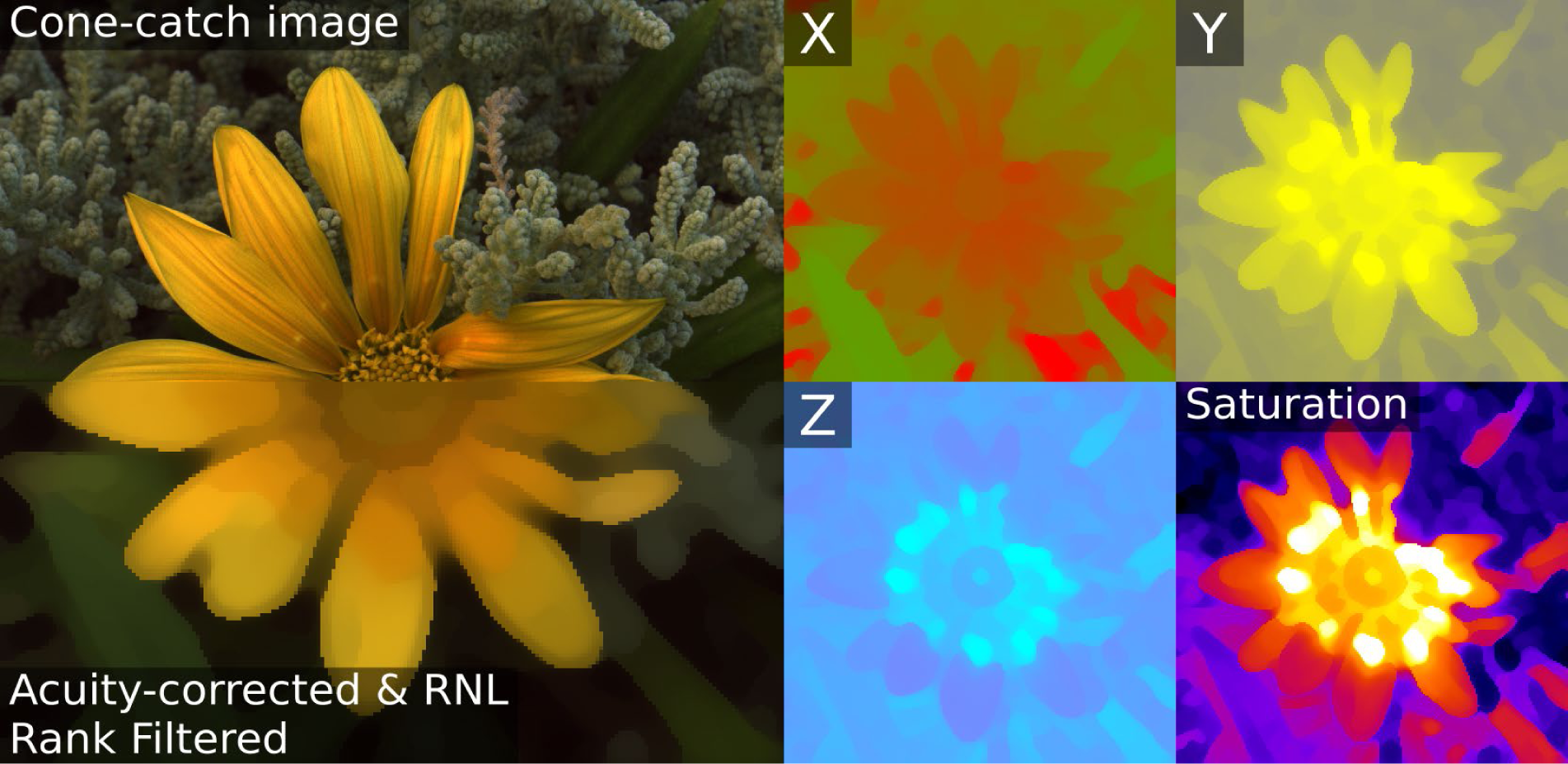
An example of the red-green (lw:mw) opponent channel (X), blue-yellow ((lw+mw)/sw) channel (Y) and the UV channel (Z) where the colour indicates the position of a pixel along that axis. The chromaticity map shows the distance of each pixel to the achromatic point.

### Step 8: Finding meaning in a pattern space

The QCPA provides a huge range of metrics from each image (currently 181 parameters), breaking each image down into a “data fingerprint“. Some of these parameters are likely to correlate well with aspects of animal evolution, behaviour and neurophysiology, while others are likely to show no signal. Likewise, some parameters will operate synergistically with each-other, while others are independent or antagonistic. Moreover, these relationships could be fundamentally different between taxa, meaning caution should be used when comparing results between highly divergent taxa (such as vertebrate versus invertebrate systems). When there is no *a priori* reason to choose the use of specific parameters for testing a specific hypothesis, we recommend the use of multidimensional data analyses, such as principal component (PCA) analysis or metric-and non-metric multidimensional scaling (MMDS/NMDS) or similar multivariate approaches to identify correlations between parameters, animal behaviour and other observational data (e.g. Winters *et al.*, 2018). Doing so can be thought of as a creating and operating in a multidimensional pattern space (for discussion see Stoddard & Osorio, 2019). Such a pattern space can include categorical data (e.g. presence/absence), data from other pattern analyses (table 1) as well as environmental data, which could include factors such as temperature, time of the year and wind direction.

## Discussion

Quantitative Colour Pattern Analysis (QCPA) is a framework for the analysis of colour patterns in nature at an unprecedented quantitative and qualitative level. At its core, QCPA uses the advantages offered by calibrated digital photography to enable the use of existing spatiochromatic colour pattern analyses (Fig. 1). It also improves existing methodologies used in visual ecology by introducing a user-friendly and open-source framework which incorporates the ability to contextualise visual scenes according to photoreceptor spectral sensitivities, receptor noise levels and abundances, natural light environments, complex natural backgrounds, spatial acuity, viewing distance and object pattern elements.

The individual modelling components of the framework rely on approximations and assumptions, which are based on our best current understanding of the underlying biological processes. As such, it is important to be aware of the limitations and underlying assumptions of the individual components of QCPA, some of which we discuss. QCPA makes extensive use of the receptor noise limited model (RNL) which has been behaviourally validated in a range of species including: humans, honeybees, birds, lizards, reef fish and freshwater fish (e.g. Vorobyev & Osorio, 1998; Vorobyev *et al.*, 2001; Champ *et al.*, 2016; Escobar-Camacho *et al.*, 2017, Sibeaux *et al.*, in review). However, the photopic version of the RNL (which we use here) was developed to model colour discrimination near the achromatic point under photopic (“bright” daytime lighting) conditions (Vorobyev & Osorio, 1998; Vorobyev *et al.*, 2001). The model is likely unable to represent the entire perceptual complexity of natural visual scenes for all species across all light regimes. For example, behavioural experiments have shown decreased sensitivity to differences in colour in specific quadrants of colour space relevant to the behavioural ecology of a species (Caves *et al.*, 2018, Sibeaux *et al.,* in review, Green *et al.* in prep). Additionally, when visual systems operate in crepuscular or scotopic conditions, the retinal stimulation to visual information becomes the result of both cone and rod stimulation or rod stimulation only (Vorobyev & Osorio, 1998). This has been discussed and implemented in past studies investigating animal vision under scotopic conditions (e.g. Kelber *et al.*, 2002; Cummings, 2004; Veilleux & Cummings, 2012) and will likely be implemented into QCPA in the near future. Further experiments are required to assess whether a “Just Noticeable Difference” (JND) corresponds to a behaviourally tested discrimination threshold of ΔS=1 throughout different parts of colour space.

QCPA allows the user to apply known sensory limitations to filter the information that is subsequently processed by low-level vision. The ability to measure image parameters is very different from being able to prove that there indeed is a link between a parameter and animal behaviour and thus ecology and evolution. While a range of the parameters provided by the QCPA have been empirically proven to be of importance in some species, many remain to be applied and investigated in a broad range of behavioural contexts and visual systems. However, herein lies one of the strengths of QCPA as it does not claim that one parameter or a way of obtaining it is superior than another. Instead, QCPA provides multiple parameters and a diverse range of image analyses, which might be of relevance in a specific situation. To what extent the observed parameterisation of visual information bears ecological or behavioural significance subsequently must be inferred and calibrated using behavioural experimentation (Olsson *et al.*, 2017).

While QCPA provides numerous parameters based on concepts shown to be relevant to a range of natural contexts (Endler, 1991, 2012; Endler & Houde, 1995; Rojas & Endler, 2013; Rojas *et al.*, 2014; Endler *et al.*, 2018; Winters *et al.*, 2018) it also provides various parameters which are yet to be validated, particularly on a quantitative scale. This provides great potential for future research as well as parameter calibration using behavioural experiments and highlights the importance and feasibility of a reductionist approach to the quantification of colour patterns and their function (*sensu* Stoddard & Osorio, 2019). Given the ability to link QCPA parameters and animal behaviour we encourage the use of the analysis to design carefully calibrated behavioural experiments in the context of complex colour patterns and visual backgrounds.

There is considerable potential to further improve QCPA by continuing to refine, test and develop its components. For example, while we currently have not included any modelling regarding the loss of spatial and chromatic information due to light scattering, particularly in aquatic or dusty environments, this would be a meaningful implementation (e.g. Nilsson, Warrant, & Johnsen, 2014). Furthermore, many animal eyes do not have uniform retinas which, in combination with diversity in eye movements and eye shapes, leads to a little investigated diversity of visual perception in addition to the already discussed perceptual diversity in animal visual systems (Land, 1999; Willis & Anderson, 2002; Land & Nilsson, 2012; Daly *et al.*, 2018; Hughes, 2018; Sibeaux *et al.*, 2019). The QCPA could also be adapted in the future to investigate moving patterns (e.g. Endler, 2012; Endler *et al.*, 2018), particularly given recent advances in the methods available for the study of colour pattern functionality in the context of motion (e.g. Fleishman, 1986; Hughes, Troscianko, & Stevens, 2014; Ramos & Peters, 2017; Murali, 2018; Nityananda et al., 2018). As outlined before, we still lack detailed knowledge about the actual physiology of colour and brightness perception in most animals, and how to represent this knowledge in terms of visual modelling in the context of perception and cognition. For example, assuming a uniform colour discrimination threshold across an entire visual scene and colour space is likely a very simplistic assumption. Furthermore, the complexity of simultaneous colour contrast and colour constancy need to be included. There are types of visual information we have barely begun understanding, such as polarisation vision, the use of fluorescence as well as their interaction with an animal’s perception of colour and brightness (Foster *et al.*, 2017; Marshall & Johnsen, 2017; Marshall *et al.*, 2018).

Recent years have seen growing diversity of colour pattern analyses (Table 1). While some use conceptually similar pattern statistics to QCPA, others provide alternative approaches such as scale invariant feature (SIFT) analysis based pattern metrics (Lowe, 1999) and combinations with models to describe cognitive aspects of attention (Rosenholtz *et al.*, 2010). The concept of QCPA based pattern analysis is entirely compatible with any of these methods. Furthermore, QCPA provides an unprecedented level of accessibility and user-friendliness by being free, open-source, graphical user interface mediated and accompanied by a vast body of support material. QCPA presents a comprehensive, dynamic and coherent work process starting with the acquisition of calibrated digital images and ending with the extraction of behaviourally and neurophysiologically contextualised pattern space. QCPA provides a promising platform and conceptual framework for future implementations of computational approaches to higher level neuronal processing of visual information. ImageJ has been the software platform of choice for image analysis for decades. Its architecture minimises the risk of non-compatibilities due to future patches of co-dependant packages (Often seen in R or Matlab) which makes QCPA (and MICA) well equipped for the medium and long-term future. ImageJ and MICA provide their own, rich, sets of image and pattern analysis and manipulation tools that QCPA profits from and can interact with. For example, GabRat (Troscianko *et al.*, 2017) can be used in combination with QCPA to investigate chromatic aspects of disruptive colouration in the context of spatial acuity. Furthermore, it is possible to use QCPA and MICA with a simple smartphone or cheap digital camera and a colour chart for calibration. While it is advantageous to have access to spectrophotometry for comparison of modelling output, this is no longer a requirement and reduces the cost for equipment drastically. There are many theories and predictions regarding the design, function and evolution of colour patterns in nature which, if at all, have only been investigated in comparably simplistic or qualitative ways. QCPA provides a powerful framework to investigate these theories in a novel quantitative and qualitative context.

## Acknowledgments

We would like to thank Simon Blomberg and Samuel Bear Powell for valuable discussions, and Miriam Heinze and Wen-Sung Chung for technical assistance.

## Authors’ contributions

C.v.d.B and J.T. conceived and tested the QCPA framework based on an original concept by C.v.d.B. J.T. wrote the software code and conceived the original (chromatic) RNL clustering and RNL ranked filter algorithms, with further testing, debugging and conceptual input and modification provided by C.v.d.B. C.v.d.B and K.L.C. conceived the combination of chromatic and achromatic discrimination thresholds for image segmentation. C.v.d.B. wrote the MATLAB based precursor of the QCPA user interface and pattern analysis which contains original code by J.A.E.

J.A.E. contributed many of the original concepts, J.A.E., K.L.C and N.J.M. have contributed to conceptual discussions and manuscript review.

