## Supplementary material for "Quantitative Colour Pattern Analysis (QCPA): A Comprehensive Framework for the Analysis of Colour Patterns in Nature": Suppl. Material

### QCPA
Quantitative Colour Pattern Analysis

Cedric P. van den Berg, Jolyon Troscianko, John A. Endler, N. Justin Marshall, Karen L. Cheney

#### Supplementary Material

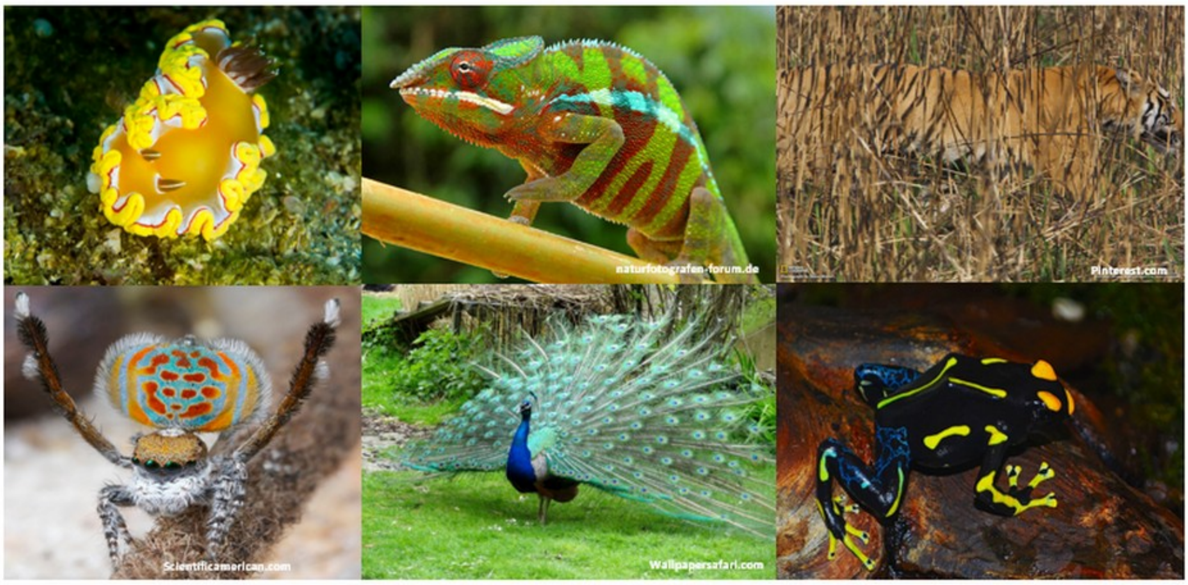

micaToolbox

empirical imaging

empiricalimaging.com

#### Introduction

This document provides detailed information on the mechanics of the individual tools. Detailed equations to all output parameters can also be found here. It also provides further discussion on important aspects of how to use QCPA and additional considerations. It also provides a range of applied examples to visualise the effect of different tools while also showing worked examples where QCPA (or parts of it) has been applied to a specific example. We do encourage the user to use the website as their primary source of information on how to use QCPA as well as further worked examples and tutorial videos.

#### Code and Download Instructions:

QCPA and its open source code (JAVA script) are available for download as part of the MICA toolbox at:

www.empiricalimaging.com

A fully functional MATLAB based precursor of QCPA can be accessed at:

https://github.com/cedricvandenberg/QCPA-Matlab

#### Manuals, Tutorials, FAQs, Forums and Updates

The website provides detailed manuals, tutorials, FAQs, a dedicated forum and updates regarding QCPA and the MICA toolbox. If you intend to use QCPA and/or the micaToolbox, please use this website to familiarise yourself with the latest updates.

#### Glossary

ΔS: The Euclidian distance between two points in the receptor noise limited opponent colour space (Vorobyev & Osorio, 1998; Hempel De Ibarra *et al.*, 2002). ΔS is simply a distance and does not make any inference on discriminability. Therefore, describing distances in colour space in terms of ΔS is correct. In fact, ΔS is the Mahalanobis distance between two stimuli, a classical measure of multivariate distance (Clark *et al.*, 2017; Endler *et al.*, 2018). Describing distances in "Just Noticeable Differences" (JNDs) is not correct because the relationship between ΔS and perception is nonlinear. JND only applies near threshold (Vorobyev & Osorio, 1998).

Weber Fraction: The Weber fraction describes the relation between the absolute intensity of a stimulus (e.g. cone stimulation) and the noise that underlies the perception (receptor noise). Weber fractions are a key concept of psychophysiology and apply to all our senses (weight discrimination, hearing, etc..). Weber fractions are a constant for a given sensory channel. It is crucial to point out that there is a moderate level of confusion about what constitutes a Weber fraction in visual ecology. We therefore recommend to strictly adhere to the original equations of Vorobyev & Osorio (1998). A detailed discussion can be found here.

JND: A "Just Noticeable Difference" (JND) describes the psychometric discrimination threshold under specific conditions. This is an estimate of the point at which the contrast between two stimuli is detectable by a sensory system. While a JND conceptually should correspond to a distance of 1 ΔS, this often is not the case due to real-time higher-level processing of visual information and possible interactions between vision and prior experience. These thresholds need to be determined with behavioural experiments and are likely to be highly context dependent.

Receptor Noise: Photoreceptors have an inherent “dark noise”. Meaning, they constantly produce a weak signal, similar to the audio noise of a radio without a signal. A visual system can only detect a signal once the level of stimulation exceeds that level of receptor noise. These noise levels, in combination with the relative abundance of each photoreceptor type and the corresponding opponent channels fundamentally determine the ability of a visual system to perceive colour and luminance contrast. It is important to distinguish between the noise level in a single neuron and the channel specific noise level determined by receptor abundance.

Hue: Hue, in an anthropocentric meaning, describes the ‘kind’ of colour we perceive. E.g. ‘red’ or ‘blue’. Physically speaking, it refers to the location of the peaks and steps in the reflectance spectra of a surface and the illuminant spectra in relation to the spectral sensitives and opponent channels of a given visual system (relative photoreceptor stimulation). In simplistic terms: hue is the photoreceptor stimulation relative to each other. This can be graphically represented as the angle of a colour relative to the achromatic spot in a colour space (see discussions in Endler, 1990 and Endler & Mielke, 2005).

Saturation: Saturation describes how ‘pure’ a colour is. The more grey it contains (the less pronounced the peaks and steps of the reflectance spectra), the less saturated it will appear. The perception of saturation is possible due to the difference between high and low intensity parts of a spectrum being captured by at least two different cone classes. This can be graphically represented as the distance to the achromatic point in a colour space.

Colour Space: The stimulation (or relative stimulation in case of opponent processing) of photoreceptors can be displayed on one or more axes. Thus, the photoreceptor stimulation a colour produces can be displayed in a n-dimensional colour space where n is defined by the number of receptors or opponent channels. For a detailed review of colour spaces see (Renoult *et al.*, 2017).

Chromaticity: Chromaticity refers to “colourfulness”. As colour is defined by both hue and saturation, chromaticity is a term that refers to both simultaneously. Opponent channels can be calculated which eliminate the achromatic signal and describe chromaticity in a single dimension of colour (such as the red-green and blue-yellow opponent systems described in humans).

Luminance: In the context of visual ecology “luminance” encompasses both the perceived “lightness” and “brightness” of a given surface and is dependent on the spectral sensitivity of the receiver receptor classes, the intensity of the signal in addition to a context-dependent cues and cognitive processes.

Brightness: Brightness refers to the perceived amount of light a given surface seems to emit or reflect. As such it is confounded by viewer perception and highly context dependent due to cognitive processes. It is frequently used incorrectly and/or loosely in the literature to mean a combination of luminance and saturation and often is used instead of saturation. It is best avoided altogether if not used in its proper definition.

Lightness: Lightness refers to the perceived reflectance or intensity of a surface. Like brightness, it is a perceptual property of surfaces and highly confounded by the perceptual context.

Spatiochromatic: Spatiochromatic is a term that implies spatial (what is where) and chromatic (colour and luminance) properties of objects or scenes being considered within each other's context.

Spectralon: A patented material consisting of sintered PTFE (Polytetrafluorethylen) powder. It is known for its near-Lambertian and spectrally flat properties because it has nearly the same reflectance, no matter what angle you look at it from. This makes it a grey standard of choice. It can be bought with various degrees of carbon in it which alters its grey value. While the Lambertian properties of Spectralon are undisputed in air, this is not the case for its use underwater. Without enough pressure from the surrounding water (e.g. in shallow water) the material's hydrophobic properties create an air-water barrier on the material's surface, which creates reflections and renders the spectralon non-Lambertian. However, how this correlates with depth is poorly researched. In shallow water sand-blasted or acid-etched marine-grade stainless steel has been used as an alternative.

RAW: RAW refers to the unprocessed information a camera’s sensory array has captured. Each manufacturer has its own RAW file format, e.g. .orf for Olympus and .nef for Nikon. It is the format of choice for calibrated photography. For detailed information see Troscianko & Stevens (2015).

JPEG: JPEG is the most commonly encountered image format. It refers to an image that has been compressed to save space, using a specific compression algorithm. However, as JPEG compression is a lossy format (e.g. it produces fringes), it is not recommended for calibrated digital photography.

Camera calibration: This refers to obtaining the spectral sensitivities of a camera’s sensory array.

Calibrated image: An image where the pixel values are linear in respect of radiance measured at the sensor, and which has been normalised so that pixel values are represented as being relative to the reflectance of a reflectance standard (thereby controlling for variations in lighting conditions and camera exposure). This is the type of image required for mapping to cone-catch images.

Cone-catch image: An image where pixel values are expressed as the predicted cone-catch quanta for a given visual system’s receptor classes.

Aposematism: A type of defensive colouration that is defined as the use of conspicuous colouration in combination with unprofitability.

Crypsis: Any type of mechanism that prevents or minimises detection.

Camouflage: All forms of concealment, including those preventing recognition as well as detection.

Mimicry: An organism that has developed similarity to another organism as a result of fitness benefits of close resemblance.

Transition matrix: Cumulative count of transitions along transects across a segmented image. The basis for most QCPA pattern statistics.

Minimum Resolvable Angle (MRA): The angular width of the narrowest black/white line pair that can be discerned by a visual system. This is similar to cycles/degree which describes how many sine waves (black to white being one wave) can be resolved within a degree of the visual field.

#### Additional Considerations for Underwater Calibrated Photography

The QCPA is intended to be applied in both terrestrial and aquatic environments. While the acquisition of calibrated digital images in terrestrial environments is well documented, no studies have yet used calibrated digital photography underwater. The aquatic environment comes with its separate set of constraints, particularly regarding the light environment. Ambient light underwater ranges from almost unaltered daylight in clear shallow water to a rapid loss of both short- and long-wavelength light in combination with decreasing light levels with depth, as well as the effects of particles an pigments in the water (Jerlov, 1976; Lythgoe, 1979). As a result, underwater photography in most cases relies on some form of artificial illumination such as strobes or video lights unless taken in shallow water. However, artificial illumination introduces three key issues. One, using a strong light source is likely to introduce light gradients within an image. This can result in artificially cast shadows and general heterogeneity of light levels within an image that can significantly degrade further image analysis. Two, unless absolutely dominating over the ambient light, the resulting illuminant will likely be an unknown mixture of natural and artificial light which makes it hard to choose a suitable illuminant in subsequent visual modelling. Three, illuminating the image with artificial light removes the natural light conditions under which the scene would usually be viewed. Furthermore, an aquatic environment can pose problems regarding the use of suitable colour and grey standards for image calibration (Voss & Zhang, 2006). These issues need to be addressed and solved prior to engaging in data acquisition and may require substantial equipment costs and testing. If the user is generally unfamiliar with underwater photography, we strongly recommend getting advice from professionals. We would like to emphasise that the quality of the images on which the subsequent analyses are conducted fundamentally constrains the meaning and realistic interpretation of QCPA output.

#### Additional Considerations for Colour & Grey Standard Choice

Ideally, the grey scales should differ in luminance by the behaviourally validated discrimination or detection thresholds of the modelled viewer. This allows for adjusting and approximating the optimum luminance discrimination threshold for each image separately while making sure that the resulting clustering agrees with behaviourally tested discrimination thresholds. Additional chromatic tiles in the colour standard allow for better adjustment and quality control of the clustering process and can also be chosen to correspond to key perceptual limits of the colour vision of a selected animal. However, the use of customised colour and grey standards can be regarded as an easy-to-implement failsafe but is not a necessary component of the RNL clustering as a lack of behavioural data is often the case when studying non-model organisms. CMOS and CCD camera sensors behave linearly with the radiance of light measured at each photosite (Maître, 2017), and linear image data can typically be extracted from RAW images produced by most consumer cameras using the appropriate software (DCRAW), or using the new linearisation modelling functions of the micaToolbox (Troscianko, unpublished). Therefore as long as the grey standard is well exposed (i.e. not saturated or so close to zero that it's value is affected by sensor noise), there is no light "bleeding" onto the sensor (e.g. due to lens flare, which is very common in ultraviolet photography), and the scene is not being photographed through a transparent layer or through mist/haze/turbidity, then only a single standard is required for normalisation where none of the image channels are over-exposed (i.e. reaching saturation point of the sensor). Ideally this standard should have a reflectance value similar to the scene average to ensure appropriate exposure. e.g. a white standard risks being over-exposed in a scene which is otherwise very dark. Otherwise two or more standards should be used, in which case the standards should ideally have reflectance values at the higher and lower end of the scene's reflectance. A standard colour chart (e.g. X-Rite color checker) has a range of grey standards, so is ideal for covering all eventualities and exposures.

#### Additional Considerations for Image Illumination when using QCPA

Image analysis based on luminance discrimination thresholds is very sensitive to gradients of illumination; therefore, it is necessary to make sure that illumination is uniform inside an image (or represents natural variation in illumination). Using diffusors, taking images in a diffuse light environment or using diffuse artificial illumination can achieve this. There are post-processing techniques, such as local mean removal, which are able to mitigate the effect of illumination gradients inside an image (Buades *et al.*, 2005). However, such post-processing may have profound effects on the results and must only be attempted with much caution and extensive calibration. Variances in both natural and artificial illumination can also lead to chromatic differences, which can have significant impacts on the perception of a visual scene (Endler, 1993). Therefore, careful consideration of illumination conditions is paramount to the use of calibrated digital photography.

#### How to Report Methodology using QCPA

QCPA provides a lot of functionality which goes along with many choices the user must make. To create reproducible research, the following information needs to be provided either in the main manuscript or in the supplement:

- Camera model & lens & Underwater housing (if applicable)
- Camera settings (Aperture, shutter speed and white balance)
- Image file type / compression (RAW, JPG, etc.)
- Camera calibration (How did you calibrate your camera)
- Illuminant (Give the spectrum AND its units)
- Colour & grey standard (What is it made of, what does it look like)
- Photoreceptor ratios (And how you normalise them)
- Receptor noise (Of the single photoreceptor and the channel specific noise)
- Spectral sensitivities (Give the original citation)
- Spatial acuity
- Viewing distance
- Colour & luminance discrimination thresholds
- ANY settings you choose in QCPA and MICA (e.g. negative value replacement, etc.)

QCPA provides an automatic file renaming system that helps in keeping track of chosen settings. However, many choices the user makes are not recorded by the file renaming and must be kept track of. For additional guidance on good methods reporting in visual ecology see White *et al.* (2015).

#### Fundamental Mechanics of CAA, VCA & BSA

All three pattern analyses (Colour Adjacency Analysis/CAA, Visual Contrast Analysis/VCA and Boundary Strength Analysis/BSA) share a common baseline mechanism: They derive their information of pattern geometry from a transition matrix. This transition matrix is created by running horizontal and vertical sampling transects across every line and column in a pixel matrix and summing up the synonymous and non-synonymous transitions along each transect and storing the information (Fig. 1S). It is this transition matrix that is then used to calculate the relative and absolute abundance of colour pattern elements and boundaries. The strength of this approach is that it conserves the information of 'What is touching what and how much' which gives rise to unique parameters which can account for adjacency. This conserved information about the identity of each patch or border type is also the basis for the combination of the spatial and chromatic properties into spatiochromatic pattern parameters. As the horizontal and vertical transects can be used to create a separate horizontal and vertical transition matrix, this allows CAA, VCA and BSA to analyse horizontal pattern properties separately from vertical ones (The user can make this choice in the interface). This has some intriguing potential application i.e. in the context of motion and viewer perspective as is discussed in both Endler (2012) and Endler *et al.* (2018).

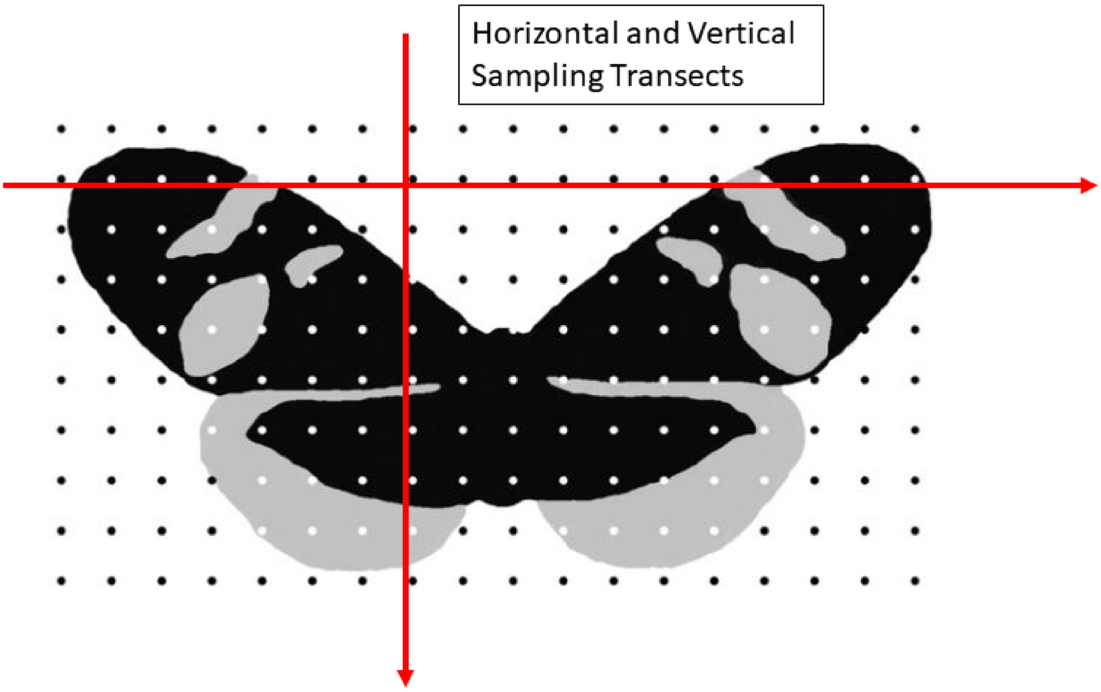

**Figure 1S:** A schematic example of horizontal and vertical transects across a clustered image of a butterfly. There are 3 cluster classes present in this image: Black, grey and white. Modified from Endler (2012).

#### Variable Summary

**<u>Note:</u>** Blue text denotes parameters which are either entirely new or have been changed from their original publication or have never been written down as a specific equation despite having been mentioned in these publications. It is not advised to blindly assume identical numerical outcomes matching the original publications for these parameters.

**General variables**

| | |
|---|---|
| $k$ | Number of colour pattern elements inside a colour pattern |
| $E$ | Number of all theoretically possible types of transitions inside a colour pattern. ($E$=$k$*($k$-1) /2). |
| $n$ | Number of types of non-zero transitions (types of non-zero off-diagonal entries) |
| $n_{off,j}$ | Number of non-synonymous transitions (e.g. red -> blue) of type *i* |
| $n_{dia,i}$ | Number of synonymous transitions (e.g. red -> red) of type *i,* the diagonals of a transition matrix. |
| $f_i$ / $f_j$ | Relative abundance of a colour pattern element *i* or *j* (from the diagonal of a transition matrix) |
| $t_{ij}$ | Relative abundance of a transition between patch i and patch j, i.e. the sum of instances in the image where a pixel of pattern element *i* is directly adjacent to pixel of pattern element *j*, divided by the total number of pattern element transitions (termed "off-diagonal" transitions) across the entire image. |
| $D_{max,i}$ | Maximum possible opponency contrast of colour pattern element *i* |
| $S_i$ or $S_j$ | Cone capture quanta or cone stimulation of colour pattern element *i* or *j* in a given class of photoreceptors |

$L_i$ or $L_j$ Cone capture quanta or cone stimulation of colour pattern element *i* or *j* in the photoreceptor channel responsible for luminance contrast perception

$\Delta S_{Sat,i}$ Euclidian distance to the achromatic point in the log-transformed Receptor Noise Limited colour space of a colour pattern element *i*. This corresponds to the saturation of a colour pattern element.

$\Delta S_{i,j}$ Euclidian distance in the log-transformed Receptor Noise Limited colour space between two colour pattern elements *i* and *j*

$\Delta S_{L,i,j}$ Euclidian distance in the log-transformed Receptor Noise Limited colour space between two colour pattern elements *i* and *j* specific to the photoreceptor channel ($S_i$) responsible for luminance contrast perception

**Output parameters of the Adjacency Analysis**

$S_c$ Simpson colour diversity

$J_c$ Relative Simpson colour diversity

$S_t$ Simpson transition diversity

$J_t$ Relative Simpson transition diversity

$H_c$ Shannon colour diversity

$Q_c$ Relative Shannon colour diversity

$H_t$ Shannon transition diversity

$Q_t$ Relative Shannon transition diversity

$S_{cpl}$ Simpson pattern complexity

$Q_{cpl}$ Shannon pattern complexity

$C$ Pattern Complexity

$PT$ Average Patch size in pixels

$A$ Aspect ratio

**Output parameters Visual Contrast Analysis**

$M_L$ Weighted mean pattern $L$ contrast

$M_{Dmax}$ Weighted mean pattern $D_{max}$ contrast

$M_{\Delta SSat}$ Mean weighted pattern $\Delta S_{Sat}$ contrast

$M_{\Delta SL}$ Mean weighted pattern $\Delta S_L$ contrast

$M_{\Delta S}$ Mean weighted pattern $\Delta S$ contrast

$s_{Dmax}$ Weighted standard deviation pattern $D_{max}$ contrast

$s_L$ Weighted standard deviation pattern $L$ contrast

$s_{Dmax}$ Weighted mean pattern $D_{max}$ contrast

$s_{\Delta SSat}$ Weighted standard deviation pattern $\Delta S_{Sat}$ contrast

$s_{\Delta SL}$ Weighted standard deviation pattern $\Delta S_{Lum}$ contrast

$s_{\Delta S}$ Weighted standard deviation pattern $\Delta S$ contrast

$CV_L$ Weighted coefficient of variation pattern $L$ contrast

$CV_{Dmax}$ Weighted coefficient of variation pattern $D_{max}$ contrast

$CV_{\Delta SSat}$ Weighted coefficient of variation pattern $\Delta S_{Sat}$ contrast

$CV_{\Delta SL}$ Weighted coefficient of variation pattern $\Delta S_{Lum}$ contrast

$CV_{\Delta S}$ Weighted coefficient of variation pattern $\Delta S$ contrast

**Output parameters of the Boundary Strength Analysis**

$BM_{\Delta S}$ Weighted mean of pattern boundary $\Delta S$ contrast

$BM_{\Delta SL}$ Weighted mean of pattern boundary $\Delta SL$ contrast

$BM_{L}$ Weighted mean of pattern boundary $L$ contrast

$BM_{Dmax}$ Weighted mean of pattern boundary $Dmax$ contrast

$BM_{\Delta Ssat}$ Weighted mean of pattern boundary $\Delta S_{Sat}$ contrast

$Bs_{\Delta S}$ Weighted standard deviation pattern boundary $\Delta S$ contrast

$Bs_{\Delta SL}$ Weighted standard deviation pattern boundary $\Delta S_{L}$ contrast

$Bs_{L}$ Weighted standard deviation of pattern boundary $L$ contrast

$Bs_{Dmax}$ Weighted standard deviation of pattern boundary $Dmax$ contrast

$Bs_{\Delta Ssat}$ Weighted standard deviation of pattern boundary $\Delta S_{Sat}$ contrast

$BCV_{\Delta S}$ Weighted coefficient of variation of pattern $\Delta S$ contrast

$BCV_{\Delta SL}$ Weighted coefficient of variation of pattern $\Delta S_{L}$ contrast

$BCV_{L}$ Weighted coefficient of variation of pattern boundary $L$ contrast

$BCV_{Dmax}$ Weighted coefficient of variation of pattern boundary $Dmax$ contrast

$BCV_{\Delta Ssat}$ Weighted coefficient of variation of pattern boundary $\Delta S_{Sat}$ contrast

#### Output parameters of the Colour Adjacency Analysis (CAA)

##### Simpson Colour Diversity ($S_c$)

A measure called the colour diversity ($S_c$) can be obtained by calculating the inverse Simpson diversity index of the diagonal of the transition matrix ($f_i$ representing the relative abundance of each colour/luminance class $i$). $S_c$ ranges from 0 to $k$ (k being the number of different colour pattern elements inside the pattern). It describes how evenly the colour/luminance classes are represented inside a pattern. $S_c=k$ when all classes are equally abundant. Therefore, the higher the number of different colour classes inside a pattern, the higher the potential maximum value of $S_c$.

$$S_c = \frac{1}{\sum_{i=1}^{k} f_i^2} \qquad (1)$$

##### Relative Simpson Colour Diversity ($J_c$)

The range of $S_c$ depends on $k$ (the number of colour pattern elements inside a pattern) and as such it can be useful to express colour diversity in relative terms, therefore making patterns more comparable if they possess different $k$. This eliminates the information on how high $k$ inside the animal is and simply shows how evenly the available colour classes are distributed

$$J_c = \frac{S_c}{k} \qquad (2)$$

Simpson Transition Diversity ($S_t$):

The regularity of a colour pattern can be described by analysing the relative transition frequencies ($t_{ij}$) which show what is next to what and how much. As stated in Endler (2012), these transition frequencies must be transformed to add up to 1, e.g. relative transition frequencies ($t_{ij}$) and not the actual values from the off-diagonal in the transition matrix, therefore $t_{ij}$ is divided by the sum of transitions in the matrix (n). As with $S_c$, the inverse Simpson diversity index of the transition frequencies is calculated resulting in $S_t$ which is ranged between 0 and $k$. $S_t=k$ when all possible transitions are equally frequent.

$$S_t = \frac{1}{\sum_{i=1}^{k}\left[\sum_{j=i+1}^{k} t_{i,j}^{2}\right]} \tag{3}$$

Relative Simpson Transition Diversity ($J_t$):

To eliminate the differences in transition diversity between patterns due to different numbers of pattern elements (*k)*, the transition diversity can be divided by the absolute number of possible transitions inside the pattern (*n*). This gives us the relative transition diversity which describes how evenly the available transitions inside the pattern are distributed independent of *k.*

$$J_t = \frac{S_t}{E} \tag{4}$$

Shannon Colour Diversity ($H_c$):

The diversity of colour pattern elements can alternatively be represented by calculating the Shannon diversity index ($H_c$) of the colour pattern elements. $H_c(max)$ = ln($k$) when all colour pattern elements ($k$) are equally frequent. This is also referred to as entropy.

$$H_c = -\sum_{i=1}^{k} f_i ln(f_i) \tag{5}$$

Relative Shannon Colour Diversity ($Qc$):

$H_c$ is confounded by the number of colour pattern elements ($k$) in the pattern. We can normalise it by dividing it by its maximum possible value, so it ranges from 0 to 1.

$$Q_c = \frac{H_c}{ln(k)} \tag{6}$$

Shannon transition diversity ($H_t$)

The diversity of transitions between colour pattern elements can be described using the Shannon index ($H_t$) of the relative transition frequencies ($t_{ij}$). $H_t(max)$ = ln($n$) when all types of non-zero transitions ($n$) are equally frequent.

$$H_t = -\sum_{i=1}^{n}\left[\sum_{j=i+1}^{n} t_{i,j} \ln\left(t_{i,j}\right)\right] \tag{7}$$

Relative Shannon transition diversity ($Q_t$)

$H_t$ is confounded by the number of non-zero transition types (*n*) in the pattern. We can normalise it by dividing it by its maximum possible value (ln(*n*)), so it ranges from 0 to 1 where 1 corresponds to a maximum diversity.

$$Q_t = \frac{H_t}{ln(n)} \quad (8)$$

Simpson colour pattern complexity ($S_{cpl}$)

As per Endler (2012), a possible way of defining colour pattern complexity is to combine $J_t$ (Relative Simpson Transition Diversity) and $J_c$ (Relative Simpson Colour Diversity). This is based on the argument that the colour patterns possessing $J_t$ and $J_c$ closest to 1 would have the highest possible complexity. It is easy to imagine this by thinking of a perfectly regular chessboard like a pattern that makes use of all available colours in equal frequency. Consequently, a low $S_{cpl}$ corresponds to a simple pattern whereas a value close to 1 would refer to a complex or even pattern.

$$S_{cpl} = (J_t + J_c)/2 \quad (9)$$

Shannon colour pattern complexity ($Q_{cpl}$)

The Shannon equivalent to $S_{cpl}$ is $Q_{cpl}$. A low $Q_{cpl}$ corresponds to a simple pattern whereas a value close to 1 would refer to a complex or even pattern.

$$Q_{cpl} = (Q_t + Q_c)/2 \quad (10)$$

Pattern Complexity (*C*):

A simple way of describing the geometric complexity of a pattern is to count the ratio between the sum off the actual diagonal values in the transition matrix ($n_{dia}$) and the sum of all values in the off-diagonal ($n_{dia}$). *C*=1 if every single pixel is adjacent to a pixel belonging to a colour class other than itself. Therefore, a pattern with more complex structures will exhibit a higher *C.*

$$C = \frac{n_{off,i}}{n_{dia,i} + n_{off,i}} \tag{11}$$

Average Patch size (*PT*)

The average patch size (*PT*) in pixels in a pattern is calculated by averaging the mean number of sequential transitions across horizontal and vertical transects where no change of colour pattern element occurs; these are the diagonal entries in the transition matrix, which need to be divided by the sum of the diagonals (the trace). The average horizontal and average vertical patch size are then multiplied with each other to calculate the average patch size in pixels. To translate the patch size into an area metric *PT* can be divided by the pixel/distance ratio derived from a scale bar. For a precise measure of patch size, we recommend the particle analysis tool in imageJ and the dedicated particle analysis in QCPA.

Aspect Ratio (*A*)

The aspect ratio describes the relation between the horizontal (h) and vertical (v) average patch size (v/h+v). A value close to 0 corresponds to a horizontally elongated pattern, whereas a value close to 1 refers to a vertically elongated pattern. A value close to 0.5 indicates circular or quadratic patterning.

#### Output parameters of the Visual Contrast Analysis (VCA)

**Colour pattern element (Patch) specific parameters:**

**Patch** maximum possible chromaticity ($D_{max}$)

The maximum possible chromatic contrast ($D_{max}$) a colour patch can elicit in a hypothetical opponent process (Endler & Mielke, 2005). Given that most opponent processing pathways are unknown in animals, this parameter calculates the cone catch contrast for each possible combination of photoreceptors in a visual system and lists the maximum. With photoreceptor stimulation $S_{i,n}$ and $S_{i,m}$, corresponding to the patch specific photoreceptor stimulation of colour patch *i* in photoreceptor class *n* or *m*. The comparison of which photoreceptors $D_{max}$ corresponds to is listed in the output summary under '$D_{max}$ *Channel*'.

$$D_{max} = max_i \left( \frac{S_{i,n} - S_{i,m}}{S_{i,n} + S_{i,m}} \right) \tag{12}$$

**Patch** Luminance ($L_i$)

Like the chromaticity of a patch, the perceived luminance of a patch ($L_i$) can be represented by the cone catch of the photoreceptor class responsible for luminance contrast detection ($S_l$). In most animals this is either a single class of photoreceptor such as the longwave (LWS) cone or the sum/mean of the cone captures of both members of the double cones.

Patch Receptor Noise Limited (RNL) Saturation ($\Delta S_{Sat}$)

The saturation of each colour pattern element can be expressed as the Euclidian distance from the achromatic point in the n-dimensional log-transformed RNL colour space ($\Delta S_{Sat}$) (Vorobyev & Osorio, 1998; Hempel De Ibarra *et al.*, 2002; Endler & Mielke, 2005; Renoult *et al.*, 2017).

**Pattern visual contrast parameters:**

Weighted mean of pattern Luminance contrast ($M_L$)

The luminance contrast of a pattern can be expressed as the mean luminance of the pattern elements *(k)* weighted by their relative abundance ($f_i$).

$$M_L = \sum_{i=1}^{k} f_i \, L_i \tag{13}$$

Weighted standard deviation pattern *L* contrast ($s_L$)

Similarly, we can calculate the standard deviation of the luminance contrast inside a colour pattern, weighted by the relative abundance of each colour pattern element ($f_i$). Where $k$ is the number of colour pattern elements in a pattern, $f_i$ is the relative abundance of a given colour pattern element, $L_i$ is the $L$ value of a given ($i$) colour pattern element and $M_L$ is the mean weighted luminance contrast of a colour pattern. Different to Endler & Mielke 2005 we make use of the standard deviation instead of the variance as proposed in Endler et al 2018.

$$s_L = \sqrt{\frac{k \sum_{i=1}^{k} f_i (L_i - M_L)^2}{(k-1) \sum_{i=1}^{k} f_i}} \qquad (14)$$

Weighted coefficient of variation pattern *L* contrast ($CV_L$)

To express the variation of the luminance contrast inside a pattern we can calculate the weighted coefficient of variation relative to the weighted mean (Endler *et al.*, 2018).

$$CV_{Dmax} = \frac{s_L}{M_L} \qquad (15)$$

Weighted mean of pattern $D_{max}$ Contrast ($M_{Dmax}$)

Similarly, we can calculate the chromatic contrast in a colour pattern by calculating the mean maximum hypothetical chromaticity of the pattern elements ($k$) weighted by their relative abundance ($f_i$).

$$M_{Dmax} = \sum_{i=1}^{k} f_i \, D_{max,i} \qquad (16)$$

Weighted standard deviation pattern $D_{max}$ contrast ($s_{Dmax}$)

We can also calculate the standard deviation of the $D_{max}$ chromaticity contrast inside a colour pattern, weighted by the relative abundance of its colour pattern elements. Where $k$ is the number of colour pattern elements in a pattern, $f_i$ is the relative abundance of a given colour pattern element, $D_{max,i}$ is the $D_{max}$ value of a given (*i*) colour pattern element and $M_{Dmax}$ is the mean weighted $D_{max}$ chromaticity contrast of a colour pattern. Different to Endler & Mielke 2005 we make use of the standard deviation instead of the variance as proposed in Endler et al 2018.

$$s_{Dmax} = \sqrt{\frac{k \sum_{i=1}^{k} f_i \left(D_{max,i} - M_{D_{max}}\right)^2}{(k-1) \sum_{i=1}^{k} f_i}} \tag{17}$$

Weighted coefficient of variation pattern *Dmax* contrast ($CV_{Dmax}$)

To express the variation of the *Dmax* contrast inside a pattern we can calculate the weighted coefficient of variation relative to the weighted mean (Endler *et al.*, 2018).

$$CV_{Dmax} = \frac{s_{Dmax}}{M_{Dmax}} \tag{18}$$

Weighted mean of pattern $\Delta S_{Sat}$ contrast $\left(M_{\Delta S_{Sat}}\right)$

And the same goes for the RNL Saturation. Note that this parameter measures saturation as the distance of each colour pattern element from the achromatic point in the log transformed RNL colour space and not a pairwise comparison inside a colour pattern as suggested by Endler & Mielke 2005.

$$M_{\Delta S_{Sat}} = \sum_{i=1}^{k} f_i \, \Delta S_{Sat,i} \tag{19}$$

Weighted standard deviation pattern $\Delta S_{Sat}$ contrast ($s_{\Delta S_{Sat}}$)

Similarly, we can calculate the standard deviation of the RNL saturation contrast inside a colour pattern, weighted by the relative abundance of each colour pattern element ($f_i$). Where $k$ is the number of colour pattern elements in a pattern, $f_i$ is the relative abundance of a given colour pattern element, $\Delta S_{Sat,i}$ is the $\Delta S_{Sat}$ value of a given ($i$) colour pattern element and $M_{\Delta S_{Sat}}$ is the weighted mean RNL saturation contrast of a colour pattern. Different to Endler & Mielke 2005 we make use of the standard deviation instead of the variance as proposed in Endler *et al.* 2018.

$$s_{\Delta S_{Sat}} = \sqrt{\frac{k \sum_{i=1}^{k} f_i \left(\Delta S_{Sat,i} - M_{\Delta S_{Sat}}\right)^2}{(k-1) \sum_{i=1}^{k} f_i}} \qquad (20)$$

Weighted coefficient of variation pattern $\Delta S_{Sat}$ contrast ($CV_{\Delta S_{Sat}}$)

To express the variation of the RNL saturation contrast inside a pattern we can calculate the weighted coefficient of variation relative to the weighted mean (Endler *et al.*, 2018).

$$CV_{\Delta S_{Sat}} = \frac{s_{\Delta S_{Sat}}}{M_{\Delta S_{Sat}}} \qquad (21)$$

Weighted mean of pattern$\Delta S_L$ contrast ($M_{\Delta S_L}$)

Similarly, the luminance contrast can be thought of as being limited by the receptor noise and photoreceptor abundance in the luminance channel (Siddiqi *et al.*, 2004). Thus, the luminance contrast in a pattern can be expressed as the RNL luminance contrast between pattern elements $i$ and $j$ ($\Delta S_{L,i,j}$)

in response to their relative abundance of the cone class responsible for luminance contrast detection and the photoreceptor specific noise weighted by the mean relative abundance ($f_i$ and $f_j$) of each colour pattern element combination ($(f_i + f_j)/2$) (Endler & Mielke, 2005). Note, this is different to the Boundary Strength Analysis (BSA) which looks at the edge contrast between adjacent colour pattern elements with respect to chromatic and luminance contrasts separately (Endler *et al.*, 2018).

$$M_{\Delta S_L} = \frac{\sum_{i=1}^{k}\left[\sum_{j=i+1}^{k}\frac{f_i + f_j}{2}\Delta S_{L,i,j}\right]}{\sum_{i=1}^{k}\left[\sum_{j=i+1}^{k}\frac{f_i + f_j}{2}\right]} \tag{22}$$

Weighted standard deviation pattern $\Delta S_L$ contrast $s_{\Delta S_L}$

Similarly, we can calculate the standard deviation of the RNL luminance contrast inside a colour pattern, weighted by the relative abundance of each colour pattern element ($f_i$ and $f_j$). Where $k$ is the number of colour pattern elements in a pattern, $f_i$ is the relative abundance of a given colour pattern element, $\Delta S_{L,i,j}$ is the $\Delta S_{Lum}$ value between two given ($i, j$) colour pattern elements and $M_{\Delta SLl}$ is the mean weighted luminance contrast of a colour pattern. Different to Endler & Mielke 2005 we make use of the standard deviation instead of the variance as proposed in Endler *et al.* 2018.

$$s_{\Delta S_L} = \sqrt{\frac{k\sum_{i=1}^{k}\left[\sum_{j=i+1}^{k}\frac{f_i + f_j}{2}\left(\Delta S_{L,i,j} - M_{\Delta S_L}\right)^2\right]}{(k-1)\sum_{i=1}^{k}\left[\sum_{j=i+1}^{k}\frac{f_i + f_j}{2}\right]}} \tag{23}$$

Weighted coefficient of variation of pattern $\Delta S_L$ contrast ($CV_{\Delta SL}$)

To express the variation of the RNL luminance contrast inside a pattern we can calculate the weighted coefficient of variation relative to the weighted mean (Endler *et al.*, 2018).

$$CV_{\Delta S_L} = \frac{s_{\Delta S_L}}{M_{\Delta S_L}} \tag{24}$$

Weighted mean of $\Delta S$ pattern contrast ($M_{\Delta S}$)

Colour contrast inside a colour pattern can result from the pattern elements being comparably contrasting to each other as opposed to being very chromatic *per se*. Thus, the chromatic contrast in a pattern can be expressed as the mean RNL chromaticity contrast between pattern elements *i* and *j* $(\Delta S_{,i,j})$ weighted by the mean relative abundance of each colour pattern element combination (($f_l$ + $f_j$)/2) (Endler & Mielke, 2005). **Note, this is different to the Boundary Strength Analysis (BSA) which looks at the edge contrast between adjacent colour pattern elements.**

$$M_{\Delta S} = \frac{\sum_{i=1}^{k}\left[\sum_{j=i+1}^{k}\frac{f_i + f_j}{2}\Delta S_{i,j}\right]}{\sum_{i=1}^{k}\left[\sum_{j=i+1}^{k}\frac{f_i + f_j}{2}\right]} \tag{25}$$

Weighted standard deviation pattern $\Delta S$ contrast ($s_{\Delta S}$)

Similarly, we can calculate the standard deviation of the chromaticity contrast inside a colour pattern, weighted by the relative abundance of each colour pattern element ($f_l$ and $f_j$). Where $k$ is the number of colour pattern elements in a pattern, $f_i$ is the relative abundance of a given colour pattern element,

$\Delta S_{i,j}$ is the Euclidian distance in log-transformed RNL colour space between two given (*i, j*) colour pattern elements and $M_{\Delta S}$ is the mean weighted luminance contrast of a colour pattern. Different to Endler & Mielke 2005 we make use of the standard deviation instead of the variance as proposed in (Endler *et al.*, 2018).

$$s_{\Delta S} = \sqrt{\frac{k \sum_{i=1}^{k} \left[ \sum_{j=i+1}^{k} \frac{f_i + f_j}{2} \left( \Delta S_{i,j} - M_{\Delta S} \right)^2 \right]}{(k-1) \sum_{i=1}^{k} \left[ \sum_{j=i+1}^{k} \frac{f_i + f_j}{2} \right]}} \qquad (26)$$

<u>Weighted coefficient of variation of pattern $\Delta S$ contrast ($CV_{\Delta S}$)</u>

To express the variation of the RNL luminance contrast inside a pattern we can calculate the weighted coefficient of variation relative to the weighted mean (Endler *et al.*, 2018).

$$CV_{\Delta S} = \frac{s_{\Delta S}}{M_{\Delta S}} \qquad (27)$$

#### Output parameters of the Boundary Strength Analysis (BSA)

<u>Weighted mean of luminance (*L*) boundary strength ($BM_L$)</u>

Similar to Endler *et al.* 2018 we can calculate the mean boundary strength in a colour pattern in terms of luminance (*L*) contrast using the following formula. The number of colour pattern elements present in a colour pattern corresponds to *k* (The length of the diagonal of the transition matrix). $t_{i,j}$ corresponds to the relative proportion of the non-zero transition frequencies (the number of transitions of a boundary type divided by the sum of all transitions). The term $\left|\frac{L_i - L_j}{L_i + L_j}\right|$ corresponds to the absolute Michelson luminance contrast of that type of boundary ($t_{i,j}$).

$$BM_L = \frac{\sum_{i=1}^{k}\left[\sum_{j=i+1}^{k} t_{i,j}\left|\frac{L_i - L_j}{L_i + L_j}\right|\right]}{\sum_{i=1}^{k}\left[\sum_{j=i+1}^{k} t_{i,j}\right]} \quad (28)$$

<u>Weighted standard deviation of luminance (*L*) boundary strength</u> $(Bs_L)$

Similar to Endler *et al.* (2018) we can calculate the standard deviation of the boundary strength in a colour pattern in terms of luminance contrast using the following formula. The number of colour pattern elements present in a colour pattern corresponds to *k* (The length of the diagonal of the transition matrix). $t_{i,j}$ corresponds to the relative proportion of the non-zero transition frequency between pattern element *i* and *j*. The term $\left|\frac{L_i - L_j}{L_i + L_j}\right|$ corresponds to the absolute Michelson luminance (*L*) contrast of that type of boundary ($t_{i,j}$). The number of different types of non-zero entries in the off diagonal of the transition matrix (types of present types of boundaries) corresponds to *n*.

$$Bs_L = \sqrt{\frac{n\sum_{i=1}^{k}\left[\sum_{j=i+1}^{k} t_{i,j}\left(\left|\frac{L_i - L_j}{L_i + L_j}\right| - BM_L\right)^2\right]}{(n-1)\sum_{i=1}^{k}\left[\sum_{j=i+1}^{k} t_{i,j}\right]}} \quad (29)$$

Weighted coefficient of variation of luminance (*L*) boundary strength ($BCV_L$)

As in Endler *et al.* (2018) we can express the variation of the boundary intensities in a colour pattern relative to the mean by calculating the corresponding coefficient of variance.

$$BCV_L = \frac{Bs_L}{BM_L} \tag{30}$$

Weighted mean of *Dmax* boundary strength ($BM_{Dmax}$)

Similar to Endler *et al.* 2018 we can calculate the mean boundary strength in a colour pattern in terms of *Dmax* contrast using the following formula. The number of colour pattern elements present in a colour pattern corresponds to *k* (The length of the diagonal of the transition matrix). $t_{i,j}$ corresponds to the relative proportion of the non-zero transition frequencies (the number of transitions of a boundary type divided by the sum of all transitions). The term $\left|\frac{Dmax_i - Dmax_j}{Dmax_i + Dmax_j}\right|$ corresponds to the absolute Michelson *Dmax* contrast of that type of boundary ($t_{i,j}$).

$$BM_{Dmax} = \frac{\sum_{i=1}^{k}\left[\sum_{j=i+1}^{k} t_{i,j}\left|\frac{Dmax_i - Dmax_j}{Dmax_i + Dmax_j}\right|\right]}{\sum_{i=1}^{k}\left[\sum_{j=i+1}^{k} t_{i,j}\right]} \tag{31}$$

Weighted standard deviation of *Dmax* boundary strength ($Bs_{Dmax}$)

Similar to Endler *et al.* (2018) we can calculate the standard deviation of the boundary strength in a colour pattern in terms of *Dmax* contrast using the following formula. The number of colour pattern elements present in a colour pattern corresponds to *k* (The length of the diagonal of the transition matrix). $t_{i,j}$ corresponds to the relative proportion of the non-zero transition frequency between pattern element *i* and *j*. The term $\left|\frac{Dmax_i - Dmax_j}{Dmax_i + Dmax_j}\right|$ corresponds to the absolute Michelson *Dmax* contrast of that type of boundary ($t_{i,j}$). The number of different types of non-zero entries in the off diagonal of the transition matrix (types of present types of boundaries) corresponds to *n*.

$$Bs_{Dmax} = \sqrt{\frac{n \sum_{i=1}^{k} \left[\sum_{j=i+1}^{k} t_{i,j} \left(\left|\frac{Dmax_i - Dmax_j}{Dmax_i + Dmax_j}\right| - BM_{Dmax}\right)^2\right]}{(n-1) \sum_{i=1}^{k} \left[\sum_{j=i+1}^{k} t_{i,j}\right]}} \quad (32)$$

Weighted coefficient of variation of *Dmax* boundary strength ($BCV_{Dmax}$)

As per Endler *et al.* (2018) we can express the variation of the boundary intensities in a colour pattern relative to the mean by calculating the corresponding coefficient of variance.

$$BCV_{Dmax} = \frac{Bs_{Dmax}}{BM_{Dmax}} \quad (33)$$

Weighted mean of $\Delta S_{sat}$ boundary strength ($BM_{\Delta S_{Sat}}$)

Similar to Endler *et al.* 2018 (But notably different) we can calculate the mean boundary strength in a colour pattern in terms of RNL saturation contrast using the following formula. The number of colour pattern elements present in a colour pattern corresponds to *k* (The length of the diagonal of the transition matrix). $t_{i,j}$ corresponds to the relative proportion of the non-zero transition frequencies (the number of transitions of a boundary type divided by the sum of all transitions). The term

$\left|\frac{\Delta S_{Sat,i}-\Delta S_{Sat,j}}{\Delta S_{Sat,i}+\Delta S_{Sat,j}}\right|$ corresponds to the absolute Michelson RNL Saturation contrast of that type of boundary ($t_{i,j}$).

$$BM_{\Delta S_{Sat}} = \frac{\sum_{i=1}^{k}\left[\sum_{j=i+1}^{k} t_{i,j}\left|\frac{\Delta S_{Sat,i}-\Delta S_{Sat,j}}{\Delta S_{Sat,i}+\Delta S_{Sat,j}}\right|\right]}{\sum_{i=1}^{k}\left[\sum_{j=i+1}^{k} t_{i,j}\right]} \tag{34}$$

<u>Weighted standard deviation of ΔS<sub>sat</sub> boundary strength</u> ($Bs_{\Delta S_{Sat}}$)

Similar to Endler et al (2018) we can calculate the standard deviation of the boundary strength in a colour pattern in terms of RNL saturation contrast using the following formula. The number of colour pattern elements present in a colour pattern corresponds to *k* (The length of the diagonal of the transition matrix). $t_{i,j}$ corresponds to the relative proportion of the non-zero transition frequency between pattern element *i* and *j*. The term $\left|\frac{\Delta S_{Sat,i}-\Delta S_{Sat,j}}{\Delta S_{Sat,i}+\Delta S_{Sat,j}}\right|$ corresponds to the absolute Michelson RNL Saturation contrast of that type of boundary ($t_{i,j}$). The number of different types of non-zero entries in the off diagonal of the transition matrix (types of present types of boundaries) corresponds to *n*.

$$Bs_{\Delta S_{Sat}} = \sqrt{\frac{n\sum_{i=1}^{k}\left[\sum_{j=i+1}^{k} t_{i,j}\left(\left|\frac{\Delta S_{Sat,i}-\Delta S_{Sat,j}}{\Delta S_{Sat,i}+\Delta S_{Sat,j}}\right| - BM_{\Delta S_{Sat}}\right)^{2}\right]}{(n-1)\sum_{i=1}^{k}\left[\sum_{j=i+1}^{k} t_{i,j}\right]}} \tag{35}$$

Weighted coefficient of variation of $\Delta S_{sat}$ boundary strength ($BCV_{\Delta S_{Sat}}$)

We can express the variation of the boundary intensities in a colour pattern relative to the mean by calculating the corresponding coefficient of variance.

$$BCV_{\Delta S_{Sat}} = \frac{Bs_{\Delta S_{Sat}}}{BM_{\Delta S_{Sat}}} \tag{36}$$

Weighted mean of $\Delta S_L$ boundary strength ($BM_{\Delta S_L}$)

As per Endler et al 2018 we can calculate the mean boundary strength in a colour pattern in terms of RNL luminance contrast using the following formula. The number of colour pattern elements present in a colour pattern corresponds to *k* (The length of the diagonal of the transition matrix). $t_{i,j}$ corresponds to the relative proportion of the non-zero transition frequencies (the number of transitions of a boundary type divided by the sum of all transitions). $\Delta S_{L,i,j}$ corresponds to the RNL Luminance contrast ($\Delta S_L$) of that a given type of boundary ($t_{i,j}$).

$$BM_{\Delta S_L} = \frac{\sum_{i=1}^{k}\left[\sum_{j=i+1}^{k} t_{i,j} S_{L,i,j}\right]}{\sum_{i=1}^{k}\left[\sum_{j=i+1}^{k} t_{i,j}\right]} \tag{37}$$

Weighted standard deviation $\Delta S_L$ boundary strength ($Bs_{\Delta S_L}$)

As per Endler et al (2018) we can calculate the standard deviation of the boundary strength in a colour pattern in terms of RNL luminance contrast using the following formula. The number of colour pattern elements present in a colour pattern corresponds to *k* (The length of the diagonal of the transition matrix). $t_{i,j}$ corresponds to the relative proportion of the non-zero transition frequency between pattern element *i* and *j*. $S_{L,i,j}$ corresponds to the RNL luminance contrast ($\Delta S_L$) of that a given boundary. The number of different types of non-zero entries in the off diagonal of the transition matrix (types of present types of boundaries) corresponds to *n*.

$$Bs_{\Delta S_L} = \sqrt{\frac{n \sum_{i=1}^{k} \left[\sum_{j=i+1}^{k} t_{i,j} \left(\Delta S_{L_{i,j}} - BM_{\Delta S_L}\right)^2\right]}{(n-1) \sum_{i=1}^{k} \left[\sum_{j=i+1}^{k} t_{i,j}\right]}} \tag{38}$$

<u>Weighted coefficient of variation of $\Delta S_L$ boundary strength</u> ($BCV_{\Delta S_L}$)

We can express the variation of the boundary intensities in a colour pattern relative to the mean by calculating the corresponding coefficient of variance.

$$BCV_{\Delta S_L} = \frac{Bs_{\Delta S_L}}{BM_{\Delta S_L}} \tag{39}$$

<u>Weighted mean of $\Delta S$ boundary strength</u> ($BM_{\Delta S}$)

As per Endler et al 2018 we can calculate the mean boundary strength in a colour pattern in terms of RNL chromaticity contrast using the following formula. The number of colour pattern elements present in a colour pattern corresponds to *k* (the length of the diagonal of the transition matrix). $t_{i,i}$ corresponds to the relative proportion of the non-zero transition frequencies between pattern element *i* and *j*. $\Delta S_{i,i}$ corresponds to the RNL chromaticity contrast ($\Delta S$) of that a given boundary (between pattern element *i* and pattern element *j*).

$$BM_{\Delta S} = \sum_{i=1}^{k} \left[ \sum_{j=i+1}^{k} t_{i,j} \Delta S_{i,j} \right] \quad (40)$$

Weighted standard deviation $\Delta S$ boundary strength ($Bs_{\Delta S}$)

As per Endler et al 2018 we can calculate the standard deviation of the boundary strength in a colour pattern in terms of RNL chromaticity contrast using the following formula. The number of colour pattern elements present in a colour pattern corresponds to *k* (The length of the diagonal of the transition matrix). $t_{i,j}$ corresponds to the relative proportion of the transition frequencies between pattern element *i* and *j*. $\Delta S_{i,j}$ corresponds to the RNL chromaticity contrast ($\Delta S$) between pattern elements *i* and *j*. The number of different types of non-zero entries in the off diagonal of the transition matrix (types of present types of boundaries) corresponds to *n*.

$$Bs_{\Delta S} = \sqrt{\frac{n \sum_{i=1}^{k} \left[ \sum_{j=i+1}^{k} t_{i,j} \left( \Delta S_{i,j} - BM_{\Delta S} \right)^2 \right]}{(n-1) \sum_{i=1}^{k} \left[ \sum_{j=i+1}^{k} t_{i,j} \right]}} \quad (41)$$

Weighted coefficient of variation of $\Delta S$ boundary strength ($BCV_{\Delta S}$)

We can express the variation of the chromatic boundary intensities in a colour pattern relative to the mean by calculating the corresponding coefficient of variance.

$$BCV_{\Delta S} = \frac{Bs_{\Delta S}}{BM_{\Delta S}} \quad (42)$$

#### Parameter Abbreviations for QCPA Results Output

To make the QCPA output file easier to navigate we have used a coded contraction of the parameters. We have divided the parameters into three families (Table 1). CAA for colour adjacency analysis, VCA for visual contrast analysis and BSA for border strength analysis.

| Variable Name | Abbreviation |
|---|---|
| Simpson colour diversity - $S_c$ (eq. 1) | CAA:Sc |
| Relative Simpson colour diversity - $J_c$ (eq. 2) | CAA:Jc |
| Simpson transition diversity - $S_t$ (eq. 3) | CAA:St |
| Relative Simpson transition diversity - $J_t$ (eq. 4) | CAA:Jt |
| Shannon colour diversity - $H_c$ (eq. 5) | CAA:Hc |
| Relative Shannon colour diversity - *Qc* (eq. 6) | CAA:Qc |
| Shannon transition diversity - $H_t$ (eq. 7) | CAA:Ht |
| Relative Shannon transition diversity - $Q_t$ (eq.8) | CAA:Qt |
| Simpson colour pattern complexity $S_{cpl}$ (eq. 9) | CAA:Scpl |
| Shannon colour pattern complexity $Q_{cpl}$ (eq. 10) | CAA:Qcpl |
| Pattern Complexity - *C* (eq. 11) | CAA:C |
| Average patch size – *PT* (no equation) | CAA:PT |
| Average horizontal patch size - $PT_{Hrz}$ (no equation) | CAA:PT Hrz |
| Average vertical patch size - $PT_{Vrt}$ (no equation) | CAA:PT Vrt |
| Aspect ratio – *A* (no equation) | CAA:Asp |

| | |
|---|---|
| Weighted mean of pattern **luminance** contrast - $M_L$ (eq. 13) | VCA:ML |
| Weighted standard deviation of pattern **luminance** contrast - $s_L$ (eq. 14) | VCA:sL |
| Weighted CoV of pattern **luminance** contrast - $CV_L$ (eq. 15) | VCA:CVL |
| Weighted mean of pattern ***Dmax*** contrast - $M_{Dmax}$ (eq. 16) | VCA:MDmax |
| Weighted standard deviation of pattern ***Dmax*** contrast - $s_{Dmax}$ (eq. 17) | VCA:sDmax |
| Weighted CoV of pattern ***Dmax*** contrast - $CV_{Dmax}$ (eq. 18) | VCA:CVDmax |
| Weighted mean of pattern **RNL saturation** contrast - $\Delta S_{Sat}$ (eq. 19) | VCA:MSsat |
| Weighted standard deviation of pattern **RNL saturation** contrast - $s_{\Delta S_{Sat}}$ (eq. 20) | VCA:sSsat |
| Weighted CoV of pattern **RNL saturation** - $CV_{\Delta S_{Sat}}$ (eq. 21) | VCA:CVSsat |
| Weighted mean of **RNL luminance** pattern contrast - $M_{\Delta S_L}$ (eq. 22) | VCA:MSL |
| Weighted standard deviation of **RNL luminance** pattern contrast - $s_{\Delta S_L}$ (eq. 23) | VCA:sSL |
| Weighted CoV of **RNL luminance** pattern contrast - $CV_{\Delta S_L}$ (eq. 24) | VCA:CVSL |
| Weighted mean of pattern **RNL chromaticity** contrast - $M_{\Delta S}$ (eq. 25) | VCA>MS |
| Weighted standard deviation of pattern **RNL chromaticity** contrast - $s_{\Delta S}$ (eq. 26) | VCA:sS |
| Weighted CoV of pattern **RNL chromaticity** contrast - $CV_{\Delta S}$ (eq. 27) | VCA:CVS |
| Weighted mean of **luminance** boundary strength - $BM_L$ (eq. 28) | BSA:BML |
| Weighted standard deviation of **luminance** boundary strength - $Bs_L$ (eq. 29) | BSA:BsL |
| Weighted CoV of **luminance** boundary strength - $BCV_L$ (eq. 30) | BSA:BCVL |
| Weighted mean of ***Dmax*** boundary strength - $BM_{Dmax}$ (eq. 31) | BSA:BMDmax |
| Weighted standard deviation of ***Dmax*** boundary strength - $Bs_{Dmax}$ (eq. 32) | BSA:BsDmax |

| | |
|---|---|
| Weighted CoV of ***Dmax*** boundary strength - $BCV_{Dmax}$ (eq. 33) | BSA:BCVDmax |
| Weighted mean of **RNL saturation** boundary strength - $BM_{\Delta S_{Sat}}$ (eq. 34) | BSA:BMSsat |
| Weighted standard deviation of **RNL saturation** boundary strength - $Bs_{\Delta S_{Sat}}$ (eq. 35) | BSA:BsSsat |
| Weighted CoV of **RNL saturation** boundary strength - $BCV_{\Delta S_{Sat}}$ (eq. 36) | BSA:BCVSsat |
| Weighted mean of **RNL luminance** boundary strength - $BM_{\Delta S_L}$ (eq. 37) | BSA:BMSL |
| Weighted standard deviation of **RNL luminance** boundary strength - $Bs_{\Delta S_L}$ (eq. 38) | BSA:BsSL |
| Weighted CoV of **RNL luminance** boundary strength - $BCV_{\Delta S_L}$ (eq. 39) | BSA:BCVSL |
| Weighted mean of **RNL chromaticity** boundary strength - $BM_{\Delta S}$ (eq. 40) | BSA:BMS |
| Weighted standard deviation of **RNL chromaticity** boundary strength - $Bs_{\Delta S}$ (eq. 41) | BSA:BsS |
| Weighted CoV of **RNL chromaticity** boundary strength - $BCV_{\Delta S}$ (eq. 42) | BSA:BCVS |

**Table 1:** Summary of the parameter abbreviations in the QCPA output file

#### AcuityView 2.0 & Gaussian Convolution Filter

Using acuity parameters of an animal (morphological or behavioural), it is possible to simulate the loss-of-contrast in an image of known angular width by removing high-spatial frequency information, often performed using Fast Fourier Transform (FFT) techniques. The methodology behind this approach is described in detail in Caves *et al.* (2016) and has been thoroughly reviewed in Stoddard & Osorio (2019). Caves and Johnsen (2017) have compiled this approach into an R package called AcuityView which we have re-written to be used in ImageJ. However, among other modifications attempting to increase user friendliness, for QCPA the input is no longer required to be of square format (Fig. 2S). However, due to the computational nature of FFT the image still needs to be rectangular. Therefore, we introduce the ability to blur irregular regions of interest (ROI) using a Gaussian filter kernel convolution (Marr, 2010). Standard Gaussian filters (e.g. those used by MATLAB or ImageJ) use separable convolutions, which require rectangular images or the use of edge padding to mitigate for non-independence of selection surroundings. This padding can introduce new (potentially misleading) pattern details. Our technique uses a custom-written non-separable convolution which ignores out-of-kernel pixels and adjusts the convolution denominator appropriately. This is more computationally intensive than applying separable convolutions in the spatial or frequency domains but can process irregularly shaped image sections completely independently of their backgrounds. At this stage the image can also be scaled to a specified number of pixels per minimum resolvable angle (MRA) to eliminate unnecessary spatial detail and increase the efficiency of subsequent processing steps. We recommend using 5 pixels per MRA to ensure no loss of spatial information.

Prior to applying acuity-control image blurring it makes sense to reduce the resolution of the image, thereby making subsequent processing steps faster without any loss of spatial information. We have determined the lowest pixel per the viewing anima's minimal resolvable angle (MRA) ratio to be approximately 5 pixels/MRA. If an image is resized to a lower resolution it will result in a loss of spatial information following acuity control, greater than 2%.

AcuityView (Caves & Johnsen, 2017) uses FFT-based processing, which requires square or rectangular images. We therefore wrote our own acuity control method which uses a Gaussian convolution, and unlike FFT or standard Gaussian blur filters which use a separable convolution (vertical and horizontal pixels convolved separately, which also requires square/rectangular images). Our Gaussian convolution is therefore more computationally intensive but is capable of acuity-control smoothing in a ROI of any shape without being affected by the ROI's surrounds or using potentially inappropriate surround manipulation (as used with separable Gaussian filters in programs such as MATLAB). The sigma of the Gaussian kernel is used to specify the desired level of blurring, and we therefore needed to determine which sigma values to use in order to reduce spatial information to a given MRA. We wrote a script which searched for the sigma level required to reduce a sine-wave image's amplitude to 2% of the original amplitude at the specified MRA given the number of pixels per MRA in the image. There was a near-perfect linear relationship between these values, and we used the model (shown in Fig. 3S) to determine the sigma level required given the user's specified pixels/MRA value.

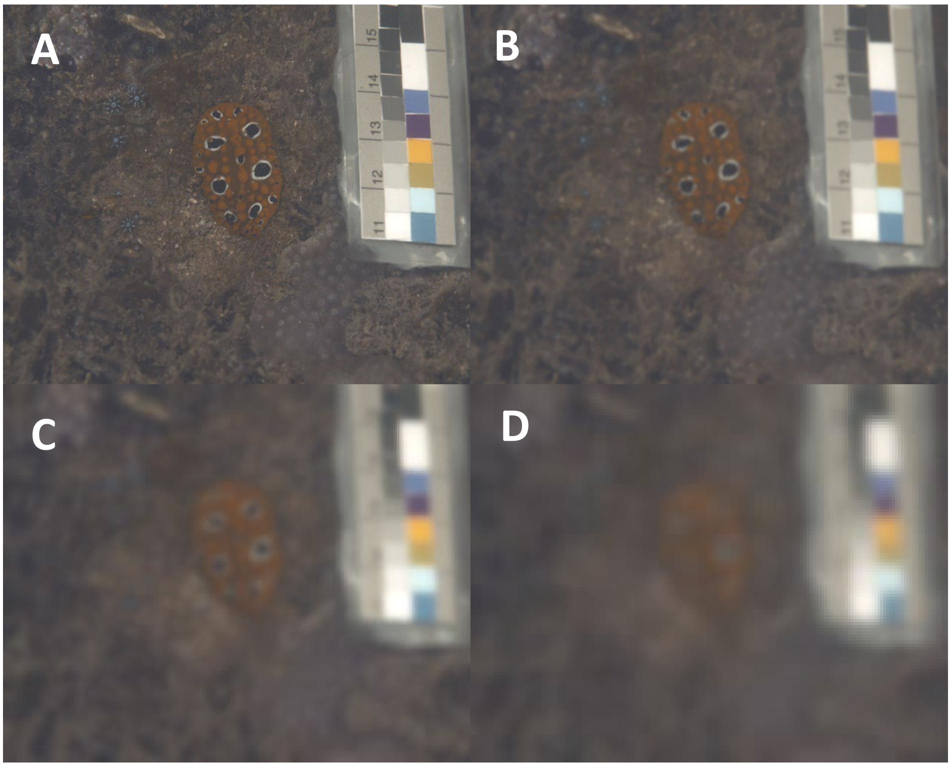

**Figure 2S:** Examples of a nudibranch, modelled as seen by a triggerfish (*R. aculeatus*) in 5m depth at various viewing distances modelled using AcuityView. A: No acuity modelling B: 10cm C: 30cm D: 50cm

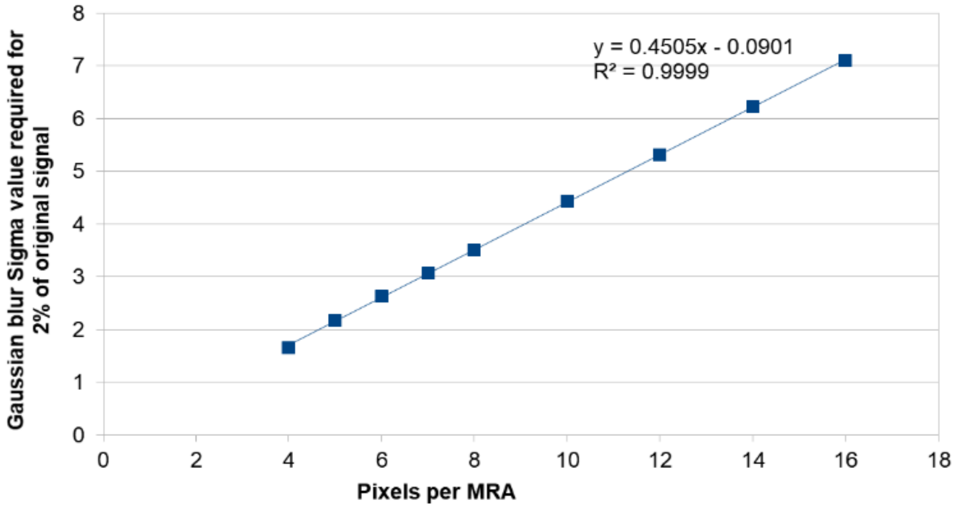

**Figure 3S:** Modelling the sigma value required to reduce a sine-wave amplitude to 2% of its original across a range of pixels/MRA values.

#### Receptor Noise Limited (RNL) Ranked Filter

The filter first compares each focal pixel's colour and luminance contrast to each of its neighbouring pixels within a given pixel scale radius selected by the user. The chromatic ($\Delta S_C$) and achromatic ($\Delta S_L$) discrimination threshold is used to rank the neighbouring pixels from those with the most to the least similar colour and luminance. Next, the focal pixel's equivalent cone-catch value is averaged with its neighbours using weighting which follows an exponential decay curve across the rankings, meaning the focal pixel is blended most with neighbours which share similar colours and brightness, and least with dissimilar neighbours. The fall-off of that smoothing curve can be specified by the user. The result is a colour and luminance discrimination threshold-based filter that reduces noise while preserving (when applied on a non-blurred image) or recovering (when applied on a blurred image) chromatic and achromatic edges.

By setting the radius of the 'RNL Ranked Filter' to equal or just above the pixel/MRA ratio the filter spans the scale of the 'blurriness' and uses appropriate Weber fractions for each receptor to remove artificial intermediate areas in the image. This restores sharp boundaries between blurred parts of the image without altering its spatial information content (Fig S4, S5 & S6). While this tool is not intended to directly mimic any particular stage of biological image processing, the resulting recovered sharp edges support the phenomenon of “hyperacuity”, whereby visual systems are capable of interpolating the location of edges at a scale which exceeds the theoretical resolution offered by the retinal arrangement of photoreceptors (Hering, 1861).

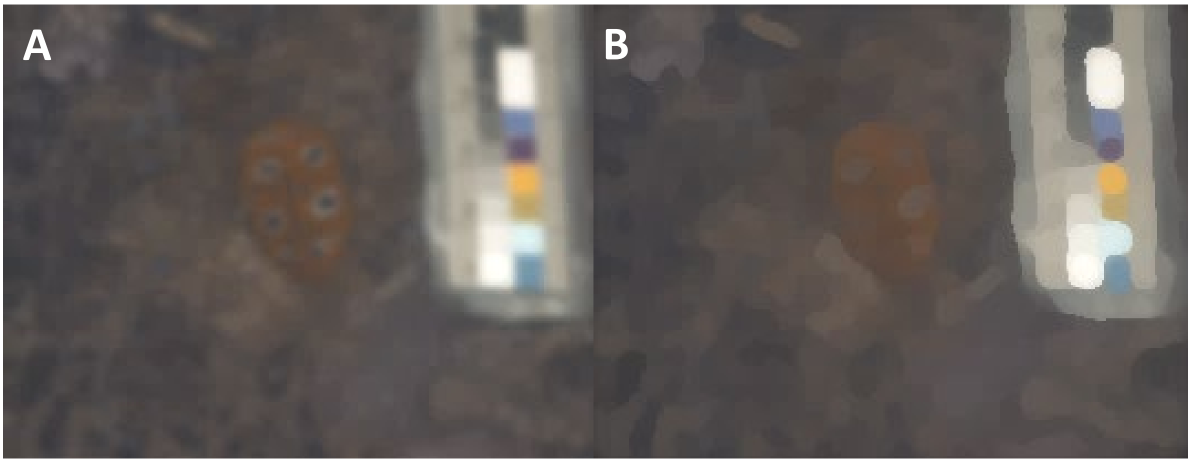

**Figure S4:** Example of an image modelled as viewed by a triggerfish from 30cm distance (A) and application of the ‘RNL Filter’ to restore sharp edges.

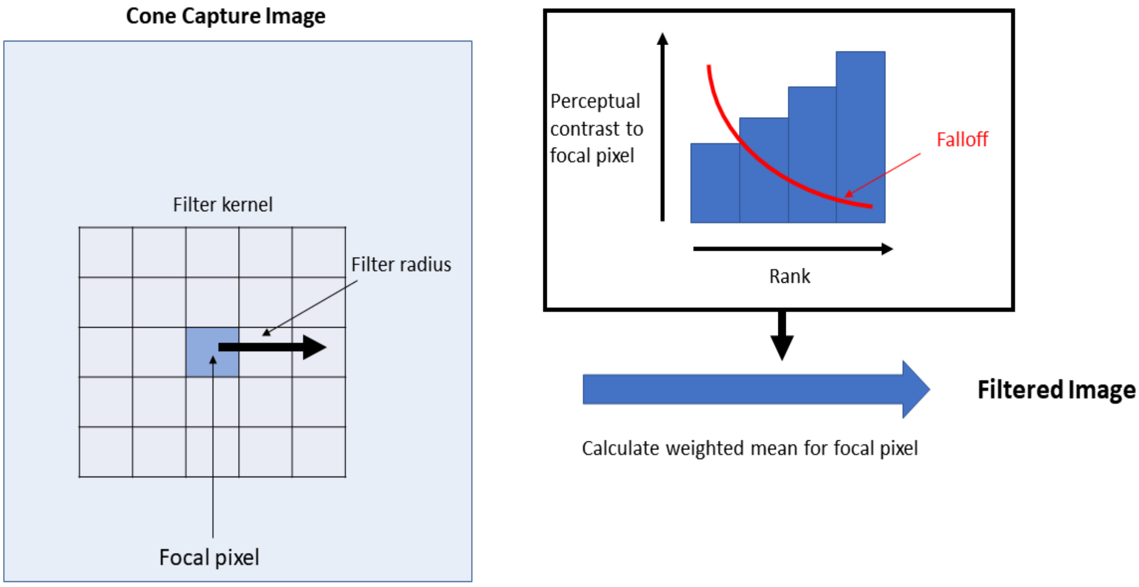

**Figure S5** Schematic representation of the RNL ranked filter. This specific example shows a 3x3 pixel filter kernel (Filter radius = 3) and a falloff close to 5. The bars in the right panel go in the opposite direction of the falloff because pixels with more similarity to the focal pixel get a rank closer to 1 which in return results in a higher weighting from the falloff curve.

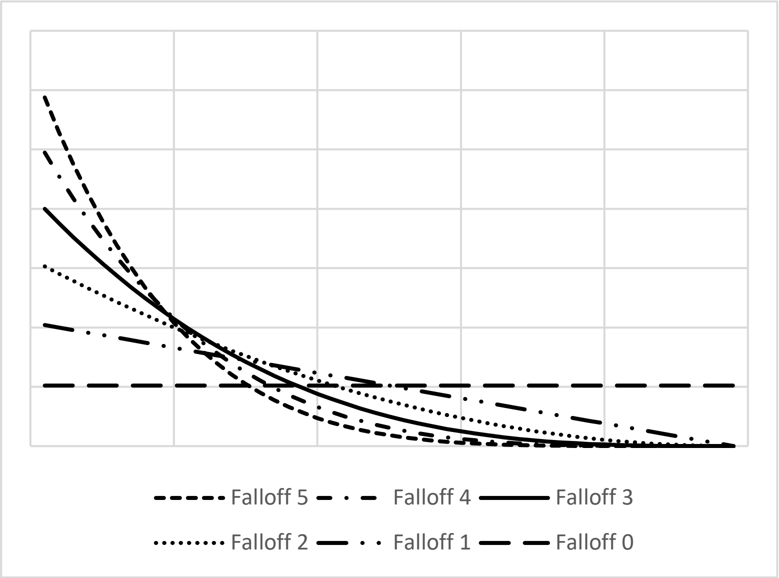

**Figure S6** Visualisation of different falloff intensities. 0 corresponds to equal weighting across the ranks within the filter kernel. 5 leads to a much more distinct weighting of the highest ranks. The area underneath all the curves = 1.

#### Receptor Noise Limited Clustering

Segmentation of cone-catch images is performed using an agglomerative hierarchical clustering approach, which is formally presented here for the first time. Initially, each pixel in the cone-catch image stack is assigned as a unique cluster with corresponding cone-catch values. The algorithm then calculates the colour and luminance distance between neighbouring clusters, pairing the most similarly coloured clusters (Fig S7). This distance is based on both colour and luminance contrast values (ΔS), creating a single distance weighted contrast measure ($\Delta S_T$) weighted by the user defined thresholds as shown in equation S1 (below) where $\Delta S_C$ is the colour contrast between two clusters, $\Delta S_L$ the luminance contrast, $S_C$ the colour discrimination threshold and $S_L$ the luminance discrimination threshold. In this example chromatic and achromatic information is weighted based on our experience of optimal clustering output. However, users may find alternative weighting to be more suitable. This is subject to ongoing research and likely highly context dependant (e.g. Kelber *et al.*, 2003).

$$\Delta S_T = \sqrt{\left(\frac{\Delta S_C}{S_C}\right)^2 + \left(\frac{\Delta S_L}{S_L}\right)^2} \qquad \text{Equation S1}$$

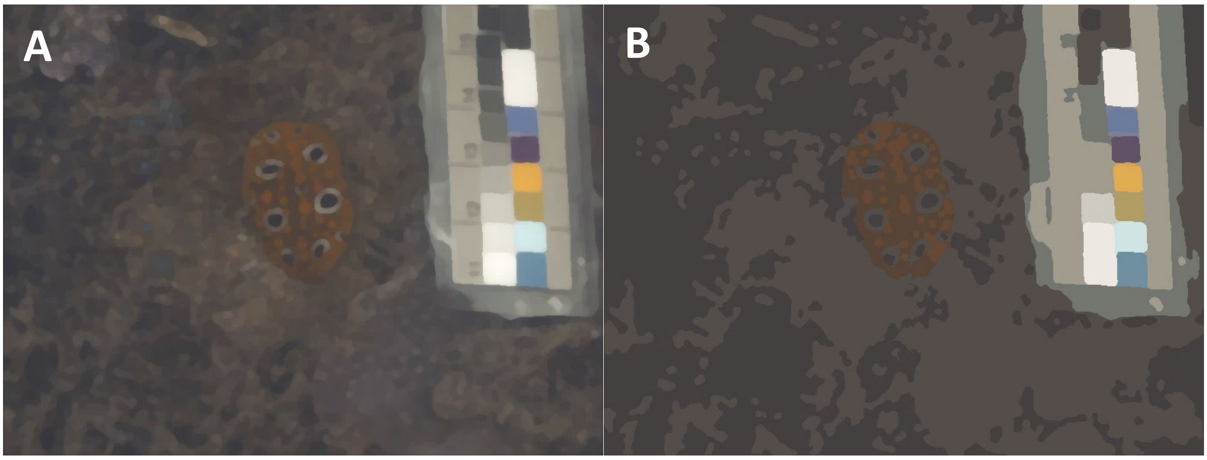

**Figure S7**: A: Example of an image as viewed by a triggerfish from 10cm at 5m depth, treated with RNL Filtering (but no clustering). B: Image A after being clustered with conservative threshold assumptions of the colour ΔS=2 and the luminance ΔS =5

If a cluster and its nearest neighbour are similar enough in colour and luminance (result in $\Delta S_T \leq k$, where k is a threshold such as 1 ΔS), then the pixels are combined into the same cluster. Combined clusters have their mean cone-catch values recalculated, weighted by the number of pixels in each cluster. The clustering algorithm repeats this process over several sequential passes. Within each pass, each cluster can be combined with one other cluster if they meet the threshold criteria. So, for example, if cluster 'A' is closest in colour to cluster 'B', and cluster 'B' is closest to cluster 'C', all three are combined into a single cluster in that pass. This method is therefore a 'single-linkage', or 'nearest neighbour' approach, which lends itself to the pairwise RNL colour comparison techniques.

The radius (or 'receptive field') over which clusters are compared is initially small (e.g. 2 pixels), however over successive passes this radius can be set to increase so that as the number of clusters decreases with each pass, the radius, and therefore number of cluster comparisons increases. The user can specify the rate at which the radius of this receptive field increases with each pass. This keeps the processing load to manageable levels (e.g. it would be computationally impractical to compare every cluster with every other cluster in a large image). This is not the same radius as the one used for the RNL ranked filter and these processes run separately and sequentially. After approximately 5 to 6 clustering passes the total number of clusters in the image have typically reduced to the thousands or hundreds, at which point it becomes more computationally efficient to compare every cluster to every other cluster. The user can specify the point at which this switch occurs, and in larger images it may be more efficient to switch to whole-image comparisons after a larger number of initial passes.

The concept of the RNL clustering resembles a graph-cut image segmentation (Greig *et al.*, 1989) and mimics basic principles of a multi-layer neural network whose inputs have small receptive fields, and increasing receptive field size in subsequent layers, so that after a number of layers inputs from the entire image can be combined. The resulting clustering mechanism allows for segmentation of an image using only colour discrimination thresholds ($\Delta S_C \leq k$), only luminance discrimination thresholds ($\Delta S_L \leq k$) or both ($\Delta S_T \leq k$) in combination, depending on the purpose of the segmentation and the relative importance of colour and luminance to a species (See suppl. Material). To achieve the best clustering results for colour and luminance discrimination we recommend *k* between 1 and 3 which also fits most critical Δ*S* values used in the literature (e.g. Osorio *et al.*, 2004; Martin Schaefer *et al.*, 2007; Stevens *et al.*, 2014, 2015; Kemp *et al.*, 2015). Using thresholds higher than 1 ΔS reflects a conservative assumption of *k* for an animal for which the RNL model has not been tested or underlying parameters such as noise levels and photoreceptor abundances are not precisely known (Vorobyev & Osorio, 1998; Vorobyev *et al.*, 2001); this applies to almost all species to date. However, we emphasise that discrimination and detection thresholds are known to be highly context dependant (e.g. Furchner *et al.*, 1977; Heinemann & Chase, 1995; Smith *et al.*, 2000; Purves *et al.*, 2002) and may extend well beyond the assumed conservative threshold of *k*=3 and may furthermore be subject to non-linearities across colour space (e.g. Mullen & Kulikowski, 1990; Sankeralli *et al.*, 2002; Cheney *et al.*, 2019). Therefore, considering perceptual and cognitive constraints that may influence the choice of suitable discrimination and detection thresholds is crucial. These thresholds should ideally be determined using behavioural experiments (e.g. Olsson *et al.*, 2017).

#### Local Edge Intensity Analysis: LEIA

The BSA uses the coefficient of variation (CoV) to describe the heterogeneity and intensity of edges in a scene. However, when a typical natural scene is processed into any kind of edge intensity image, the distribution of these edges is bounded at zero, with an intense skew (somewhat like a Poisson or gamma distribution) due to the low frequency of high-contrast edges. This means the CoV – which assumes a normal distribution – does not adequately capture the level of variation in the scene. We therefore introduce the option for log or square root transforming the ΔS values to create normal distributions, and additionally we include parameters which can capture this higher order deviation in ΔS distribution: skewness and kurtosis. These describe the shape of this non-normal edge intensity distribution using standardised moment measures on top of the standard deviation and CoV. An edge intensity distribution with a large right-hand tail would have a higher skewness value, while a distribution with a long tail of extreme outliers would have a higher kurtosis value. A pattern with high edge contrast skewness would therefore be more complex (including both high and low-contrast edges), while a high kurtosis might indicate the pattern is more salient (characterised by predominantly low-contrast edges, but with a large number of extreme-contrast edges). However, we provide the user with the option to log and sqrt transform the edge intensity values as well as ignore sub-threshold values prior to analysis which has profound impact on these output parameters and their meaning in a given context. We currently do not provide numerical examples (i.e. example distributions) but will certainly update these to the website and/or a later stage of the manuscript.

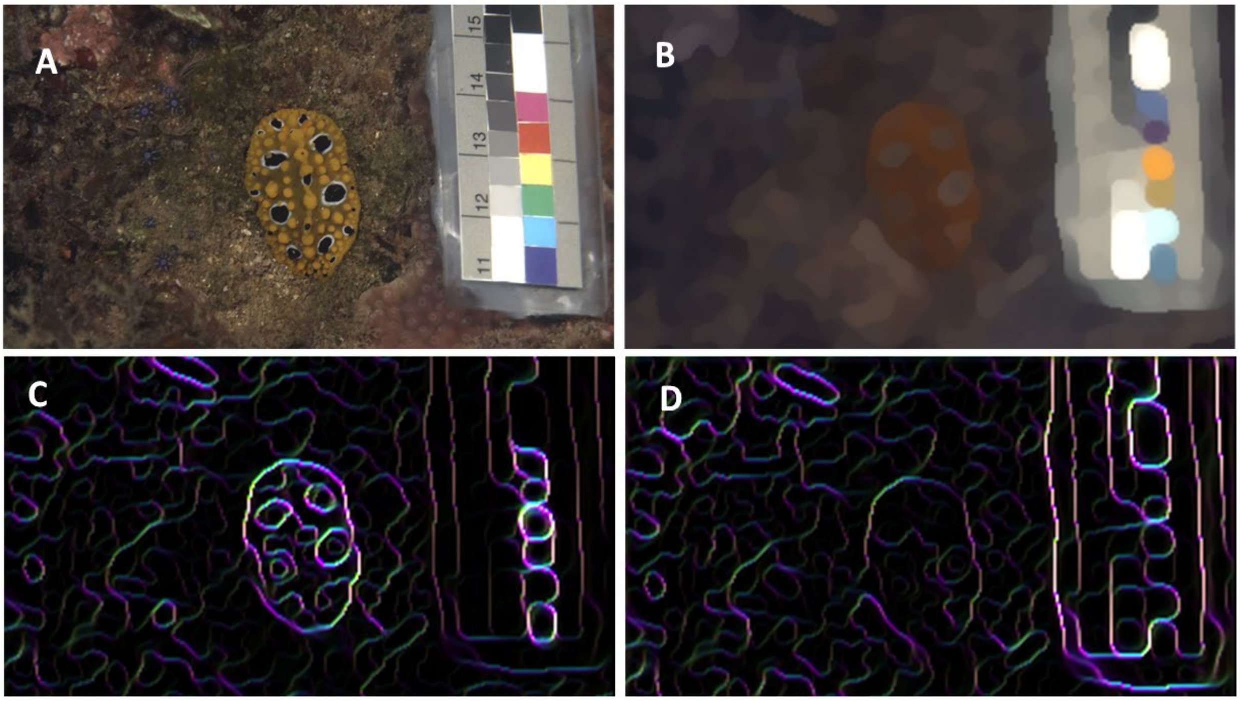

**Figure S8:** Example of an unclustered image (A), treated with RNL Filtering and subsequent extraction of the chromatic ΔS Edge Maps. B shows a RGB reconstruction of the image after modelling of triggerfish (*R. aculeatus*) colour and spatial vison at 30cm distance in a greenish natural light environment. C shows a composite image of all 4 directions of the chromatic edge contrast, each indicated with a different colour. Blue=horizontal, Yellow=vertical, Green=diagonal top left to bottom right, Magenta= diagonal top right to bottom left. The brightness corresponds to the edge intensity (See suppl. Material for details). D is the equivalent of C but using the luminance contrast in ΔS.

#### XYZ coordinates and Chromaticity Heat Maps

XYZ coordinates can be calculated for each pixel using the RNL equations provided by Renoult et al. (2017). The saturation image shows $\Delta S_{Sat,i}$ values (i.e. the distance of each pixel's colour to the achromatic point, equivalent to saturation).

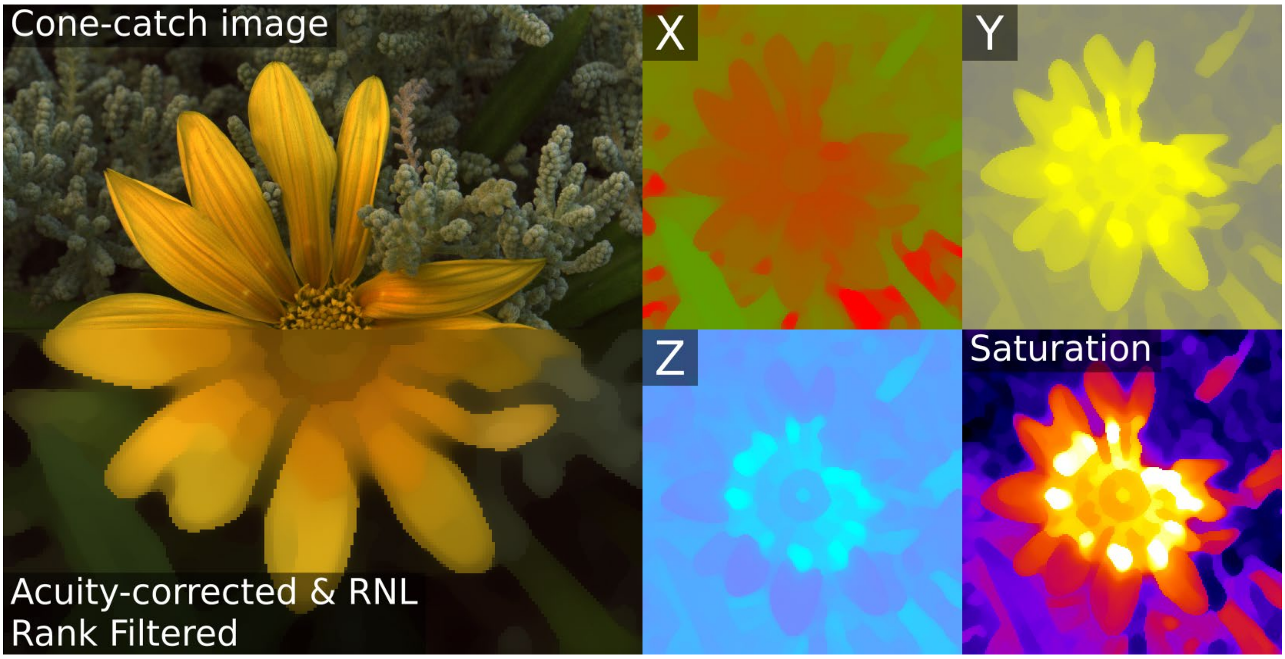

**Figure S9** An example of the red-green (lw:mw) opponent channel (X), blue-yellow ((lw+mw):sw) channel (Y) and the UV channel (Z) where the colour indicates the position of a pixel along that axis. The saturation image shows the distance of each pixel to the achromatic point.

#### Naïve Bayes Clustering

If a pattern consists of very distinctly coloured elements or vision parameters are unknown, QCPA provides an alternative image segmentation technique, Naïve Bayes Clustering. The user can define a set of clusters by selecting corresponding pixels from a given pattern element (e.g. the yellow petal of a flower). Using a Naïve Bayes classifier (Domingos & Pazzani, 1997) the rest of the pixels in a pattern can be attributed to each of the user defined categories based on the probability of belonging to each of the categories. This approach is similar to the frequently used k-means clustering (Hartigan & Wong, 1979). However, Naïve Bayes Clustering allows the user to pre-define the clusters by making active selections in a pattern whereas k-mean clustering segments an image simply based on the similarity of each pixel in an image to each other and a pre-defined number of clusters. Thus, naïve Bayes clustering is a lot more interactive and allows a more tailored segmentation than k-means.

#### Impact of RNL Ranked Filter: Falloff

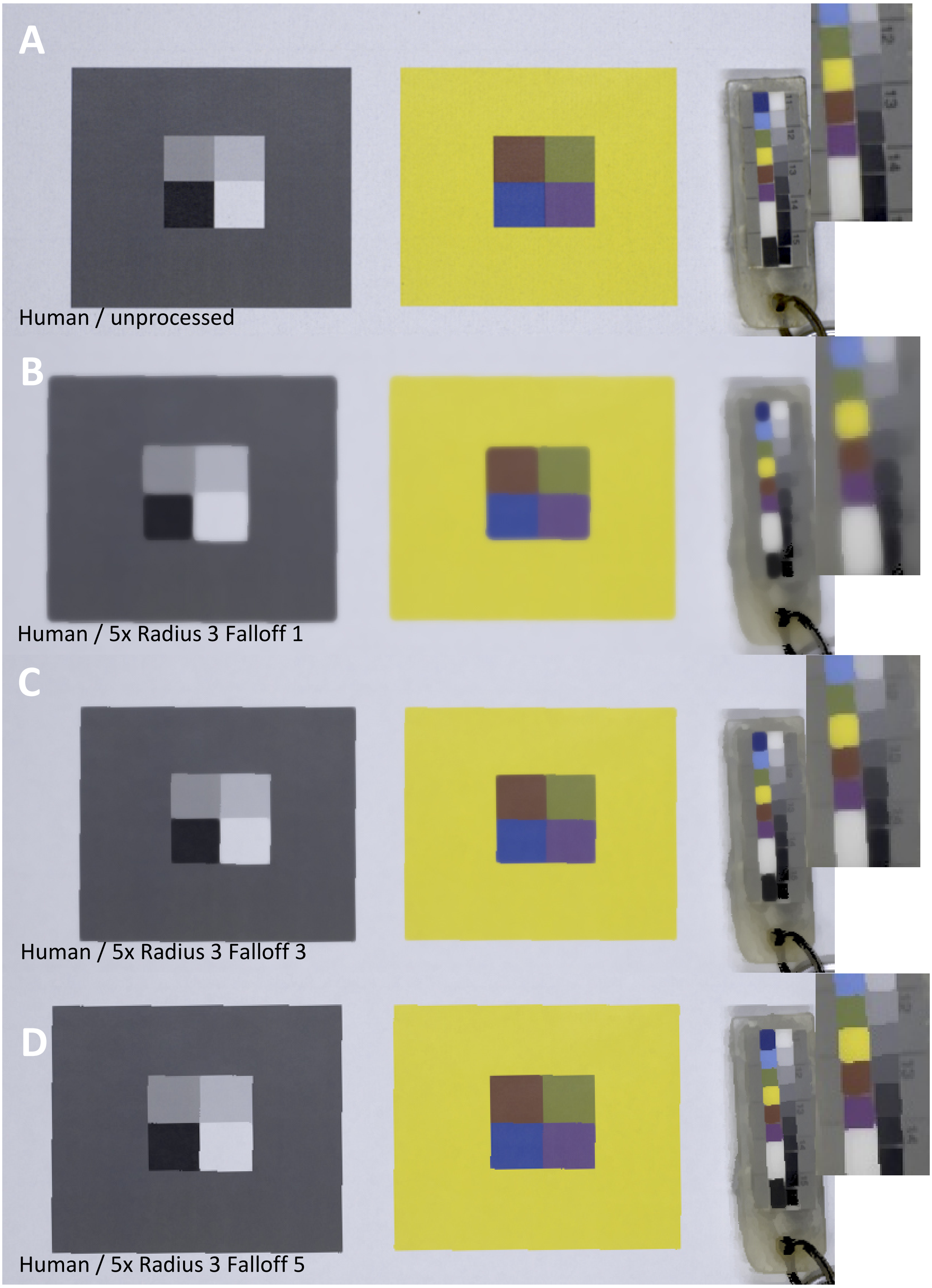

**Figure S10** Examples showing the impact of varying the falloff intensity

#### Impact of RNL Ranked Filter: Radius

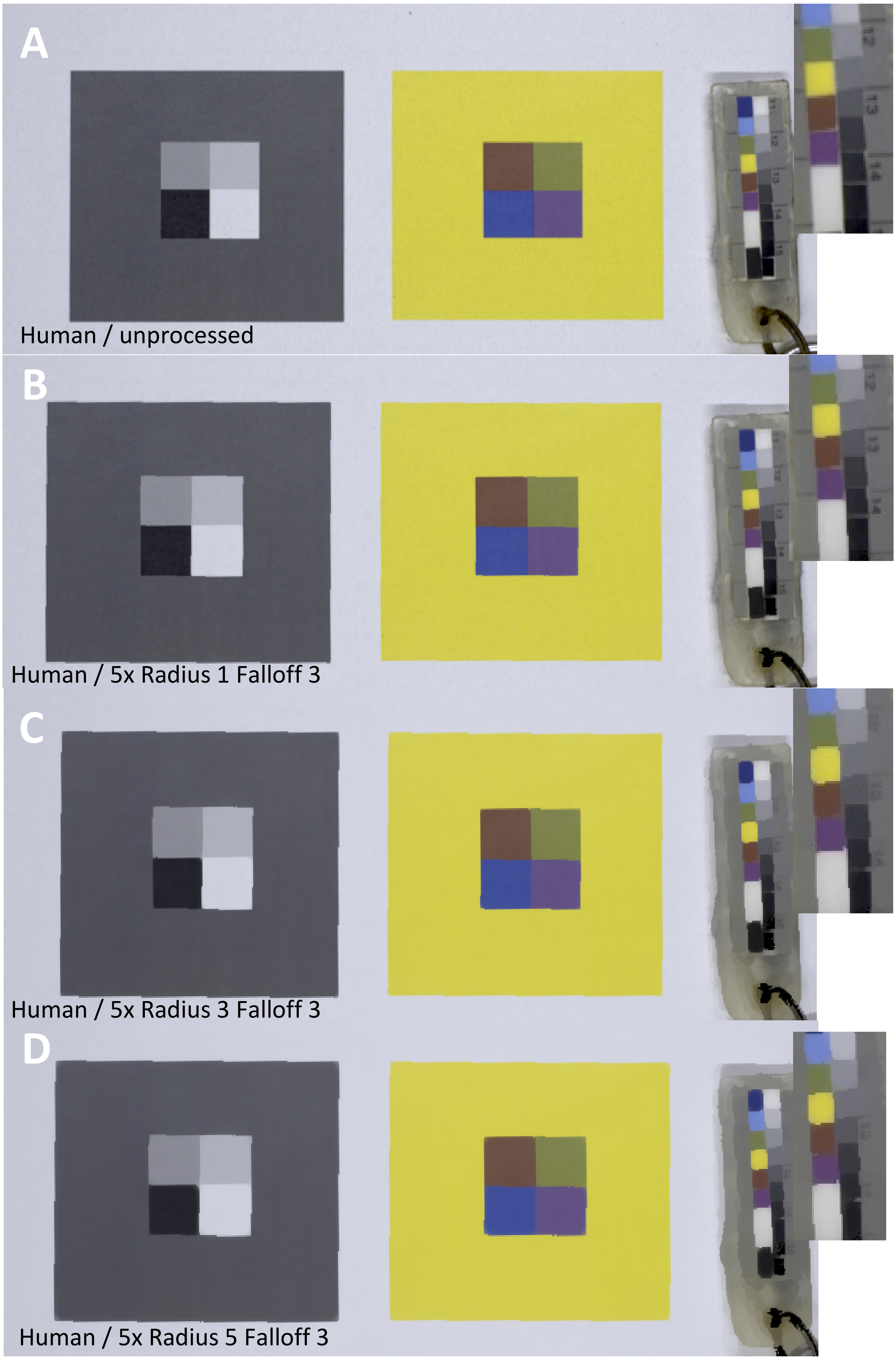

**Figure S11** Examples showing the impact of varying the filter radius.

#### RNL Clustering and RNL Ranked Filter in Combination

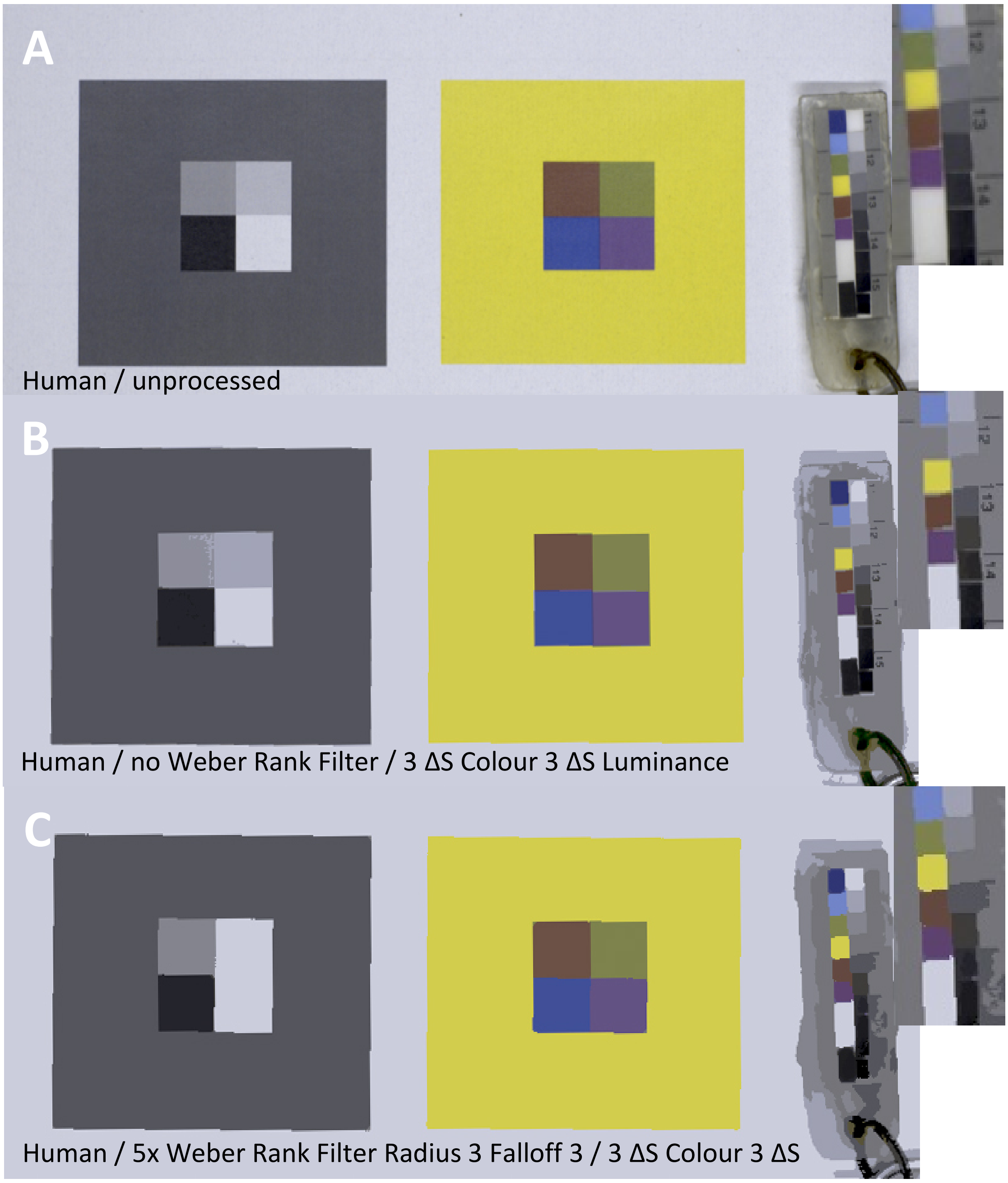

**Figure S12** Example showing the impact of the RNL Ranked Filter on the RNL Clustering. Note the loss of small spatial detail between B & C but the increase in 'smoothness'. However, this can be modulated by choosing different filter settings to suit the best outcome.

#### RNL Clustering: Colour vs. Luminance Clustering

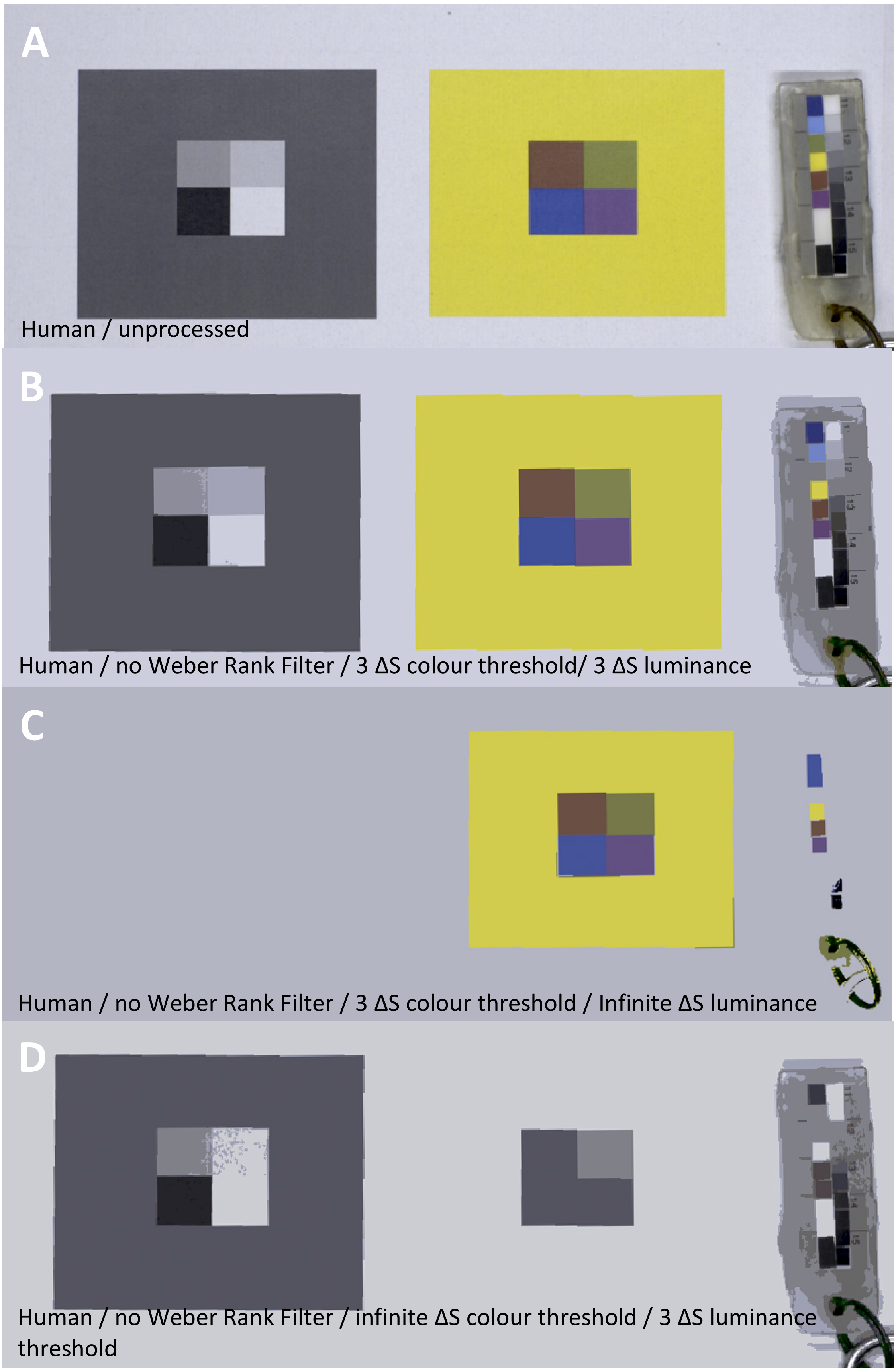

**Figure S13** Example showing the impact of clustering with both an achromatic and chromatic threshold (B), only a chromatic threshold (C) and only an achromatic threshold (D).

#### AcuityView, RNL Ranked Filter and RNL Clustering in Combination

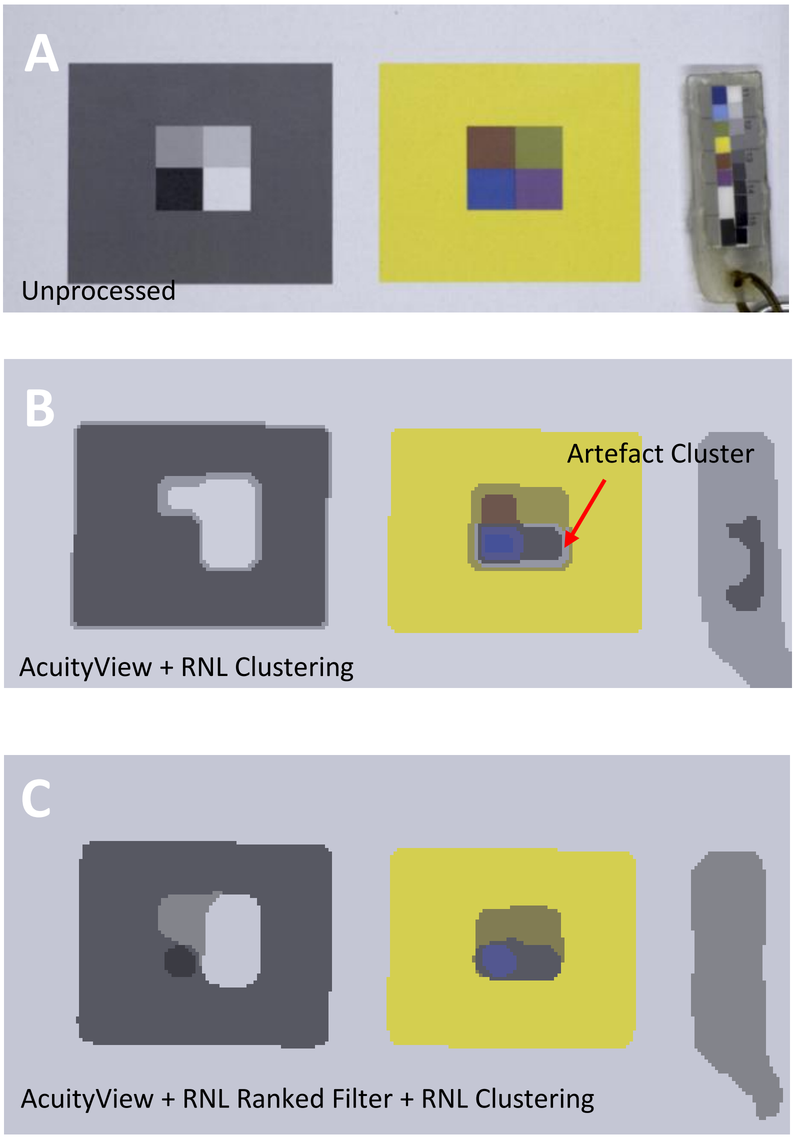

Human / unprocessed

Human / 3 cycles/degree from 0.5m distance/ 3 ΔS Colour / 3 ΔS Luminance threshold / no Weber Filter

Human / 3 cycles/degree from 0.5m distance/ 3 ΔS Colour / 3 ΔS Luminance threshold / 5x radius=MRA (5) Falloff 3 Weber Filter

**Figure S14.** Example showing the difference between using (C) and not using (B) the RNL Ranked Filter after modelling spatial acuity. Note the resulting artefact clusters in B.

#### Worked examples

Example 1: Visual defences in Nudibranchs through the eyes of a triggerfish

Nudibranch molluscs are a diverse family of shell-less marine gastropods. Many of them are thought to be highly aposematic using vivid colour patterns together with chemical and mechanical defences to deter predators (Tullrot & Sundberg, 1991; Haber *et al.*, 2010; Carbone *et al.*, 2013). They have become an increasingly popular model organism for the study of the design, function and evolution of aposematic colouration (Cortesi *et al.*, 2009; Cheney *et al.*, 2014; Winters *et al.*, 2017). In this example we show the use of QCPA and its components to compare the colour pattern of two individuals of nudibranchs through the eyes of a potential predator, the lagoon triggerfish (*Rhinecanthus aculeatus*). The first one is an individual of the species *Phyllidia ocellata*, an assumedly aposematic species with high levels of chemical defence (Cheney *et al.* unpublished data). The second animal belongs to the species *Dendrodoris krusensternii*, an assumedly camouflaged species with no known chemical defences.

We first show the spectral sensitivities of the camera that was used, the visual system of the triggerfish as well as the light spectra under which the images had been taken and finally the light spectra for which we used the MICA toolbox to calculate the triggerfish's photoreceptor stimulation. We show the original image captured by the camera and the reconstructed RGB image based on the photoreceptor stimulation of a triggerfish at 5m depth in clear water.

We then estimate the spatial information available to our triggerfish viewer as per a given viewing distance and known visual acuity (Champ *et al.*, 2014) using Fast Fourier Transform (FFT) based acuity modelling. We then recreate distinct boundaries in the image (As the fish is unlikely to perceive a blurred image). This is done by using the RNL Ranked Filter. We set the Filter so that its radius equals the Minimum Resolvable Angle to which the image has been rescaled (In this case, 5 pixels per MRA). This prevents the creation of artificial clusters when we then continue to segment the image into its colour pattern elements using the RNL Clustering. We can use both the clustered and the un-clustered image to derive secondary image statistics and data visualisations. These steps can be done at multiple simulated viewing distances or light environments, which we do not do here in this example.

Similarly, we can analyse the second individual from the species *Dendrodoris krusensternii*. We can then compare some selected image statistics to quantify the degree of background matching in either one of the animals. We can also quantify the salience of each animal colour pattern.

**Spectra and Sensitivities**

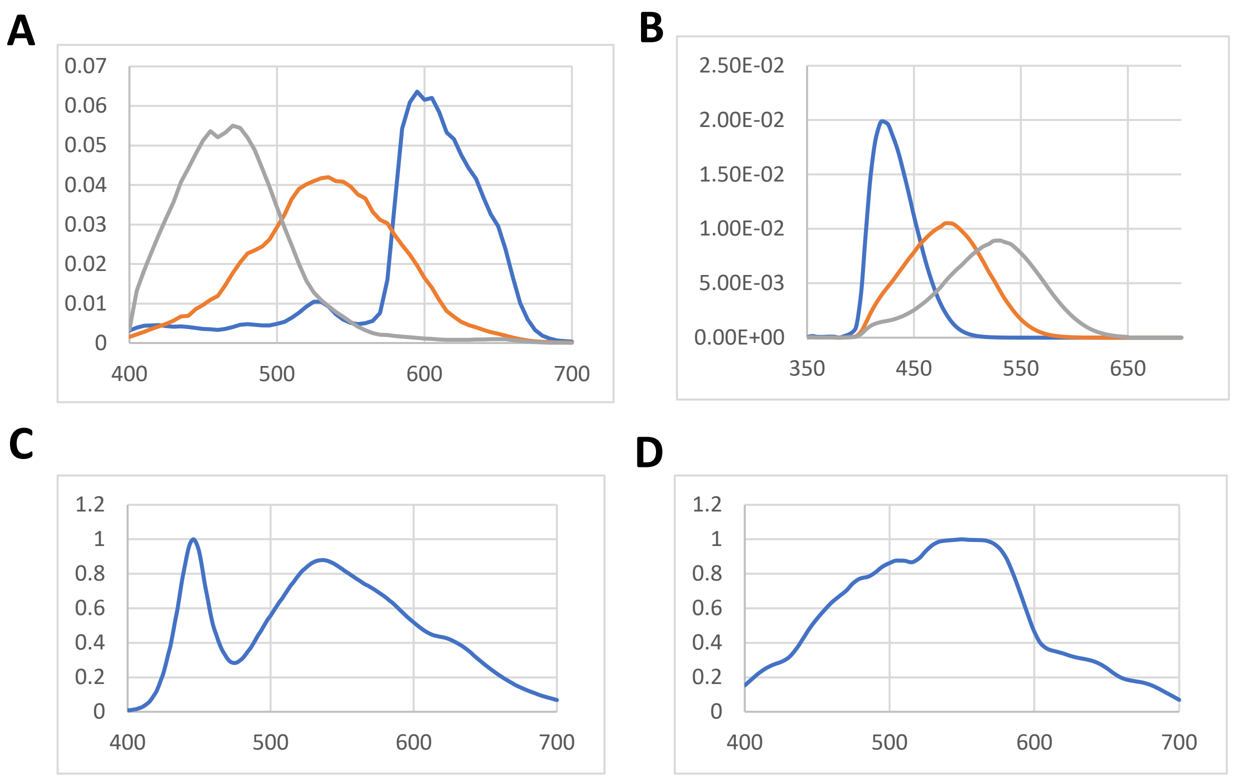

**Figure S15:** A: Spectral sensitivity of the camera used in this example (Olympus PEN-EPL5). The spectral sensitivity is available in the micaToolbox. B: Spectral sensitivity of the lagoon triggerfish (*R. aculeatus*) C: Light spectrum of the white LED video lights used for photography D: Light spectrum at 5m depth with a moderate amount of green algae in the water column.

***Phyllidia ocellata* (Aposematic nudibranch)**

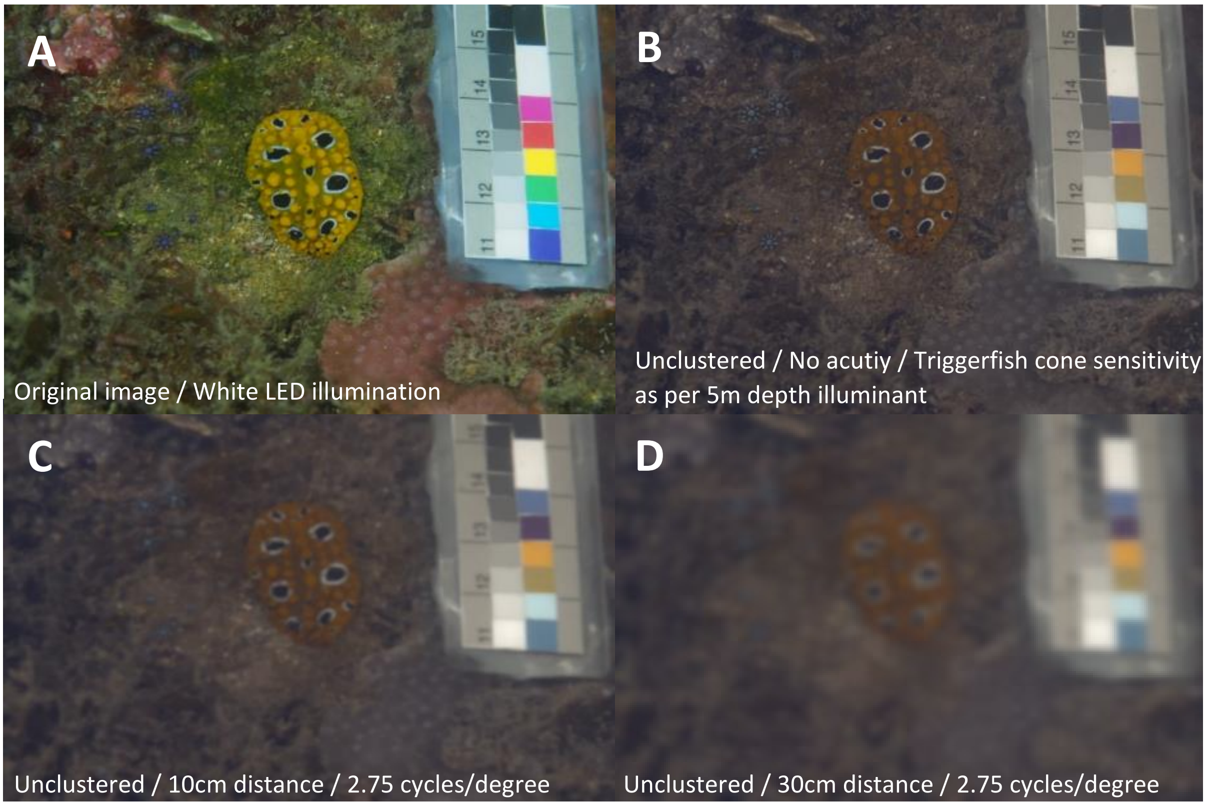

**Figure S16:** Modelling cone capture quanta (B) and spatial acuity (C&D).

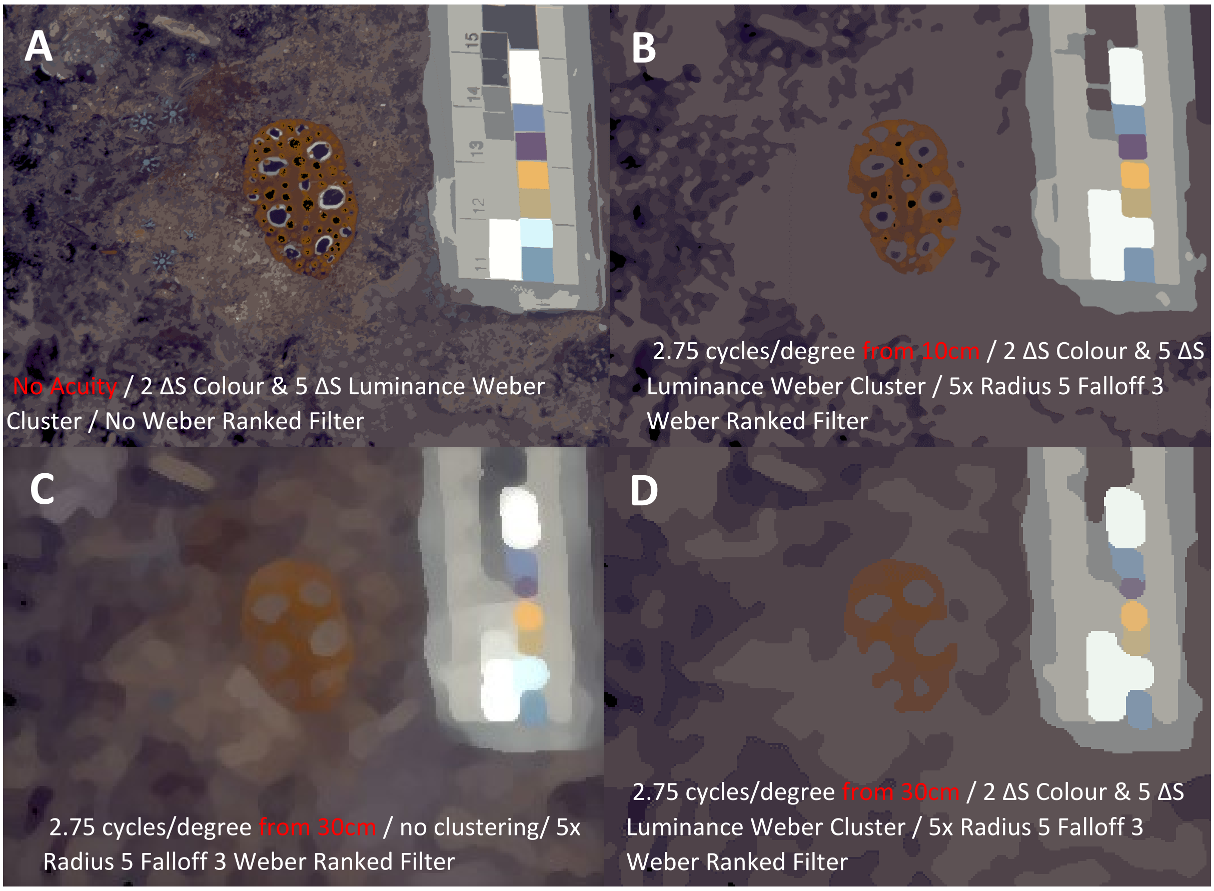

**Figure S17:** Clustering the image and recreating distinct pattern boundaries

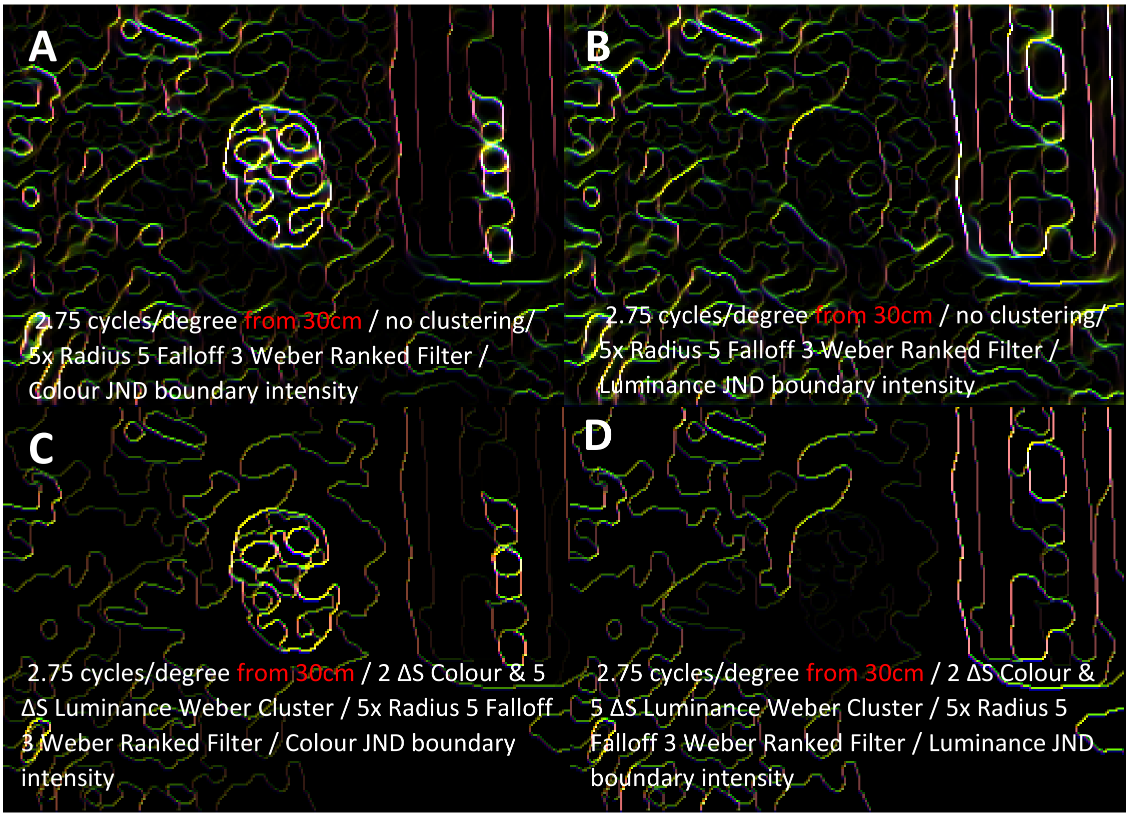

**Figure S18:** Using the un-clustered image (Fig S17c) to visualise local edge contrast with LEIA. These images can be quantified using the LEIA output parameters.

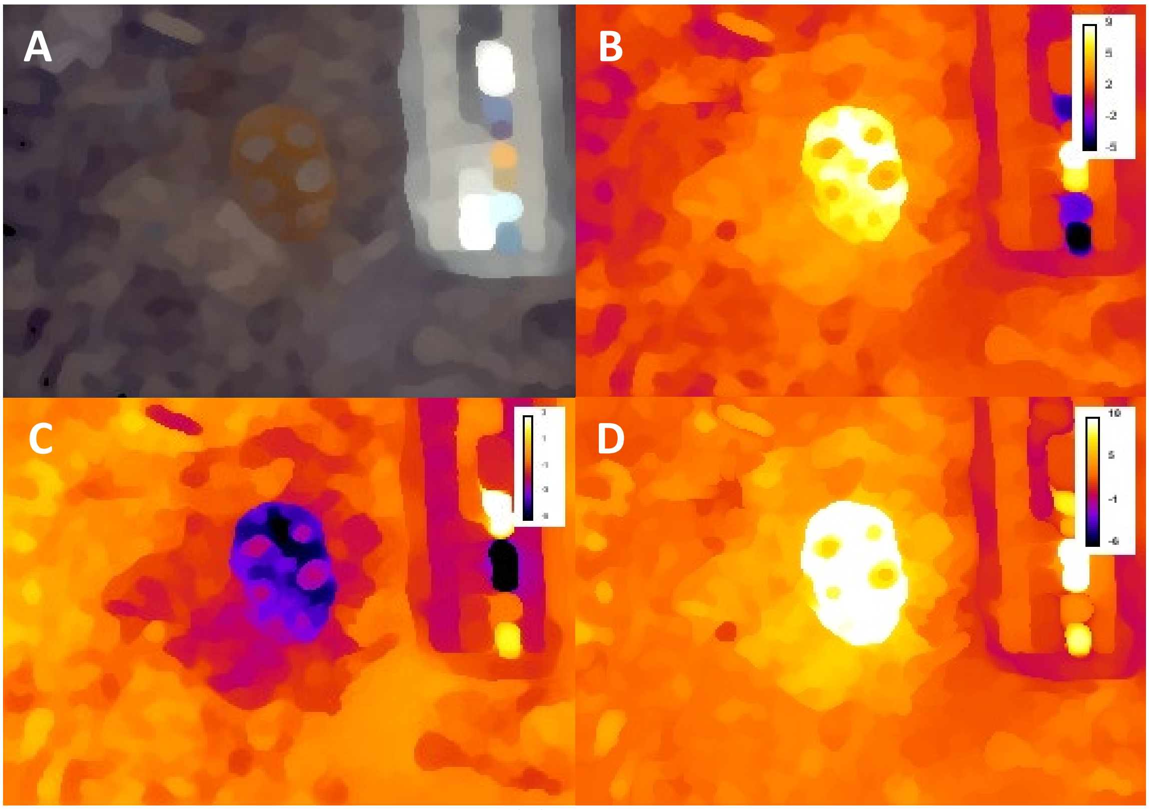

**Figure S19:** Using the un-clustered image (A) to visualise opponent channel stimulation in 'XYZ Chromaticity Images'. B: The X-axis of colour space. C: The Y-axis of colour space. Note that there is no Y-axis as we are using a tri-chromatic visual system. D: Saturation Image. The user may want to quantify these images using pattern analysis or image analysis tools provided in ImageJ and MICA.

*Dendrodoris krusensternii* **(Cryptic Nudibranch)**

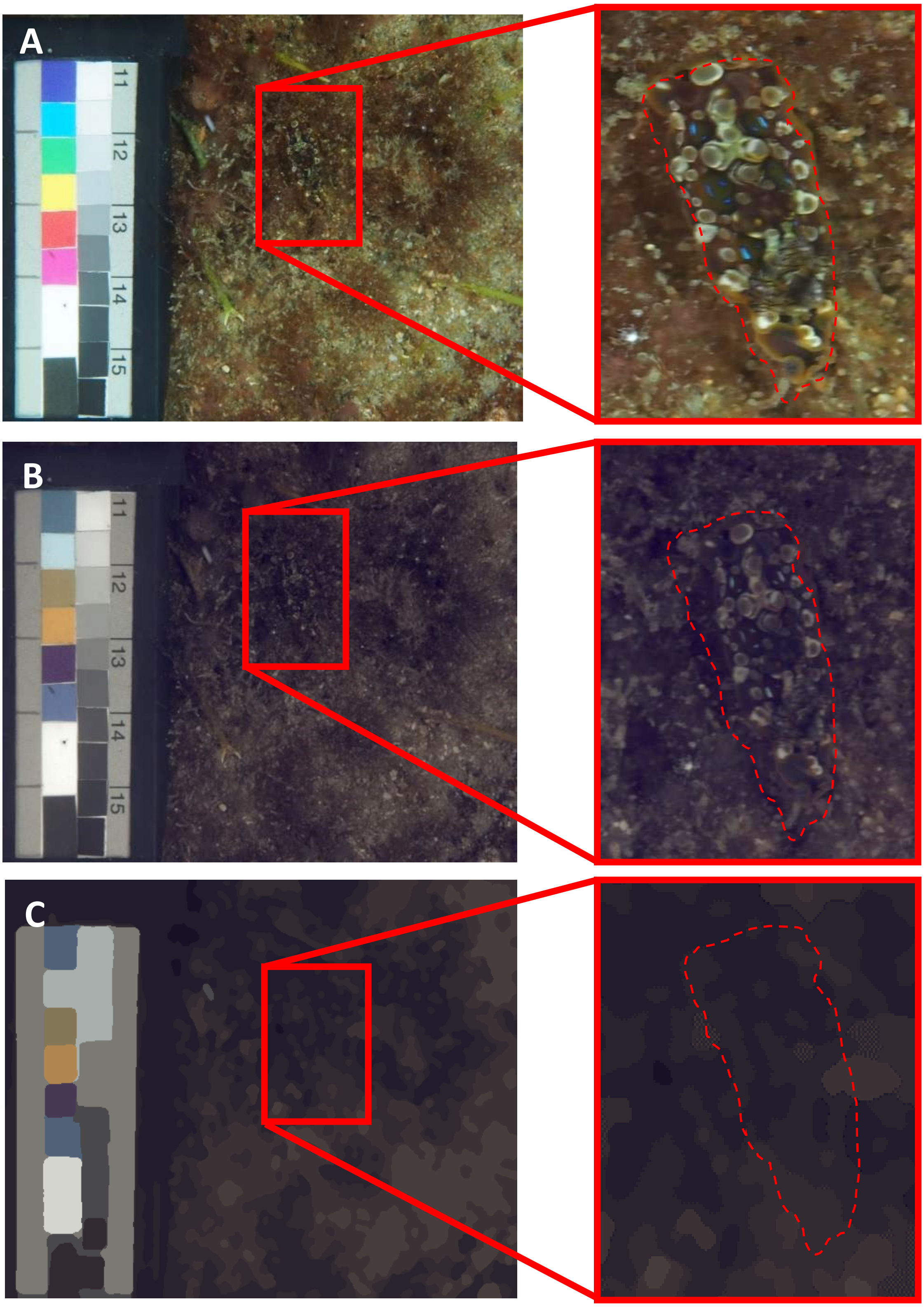

**Figure S20:** A) RAW image of a cryptic nudibranch (*Dendrodoris krusensternii*) against its natural background. The animal is highlighted with a dashed red line. B) The same image transformed into triggerfish vision (*R. aculeatus*) without visual acuity modelling. C) With RNL ranked filter and RNL clustering but no visual acuity modelling.

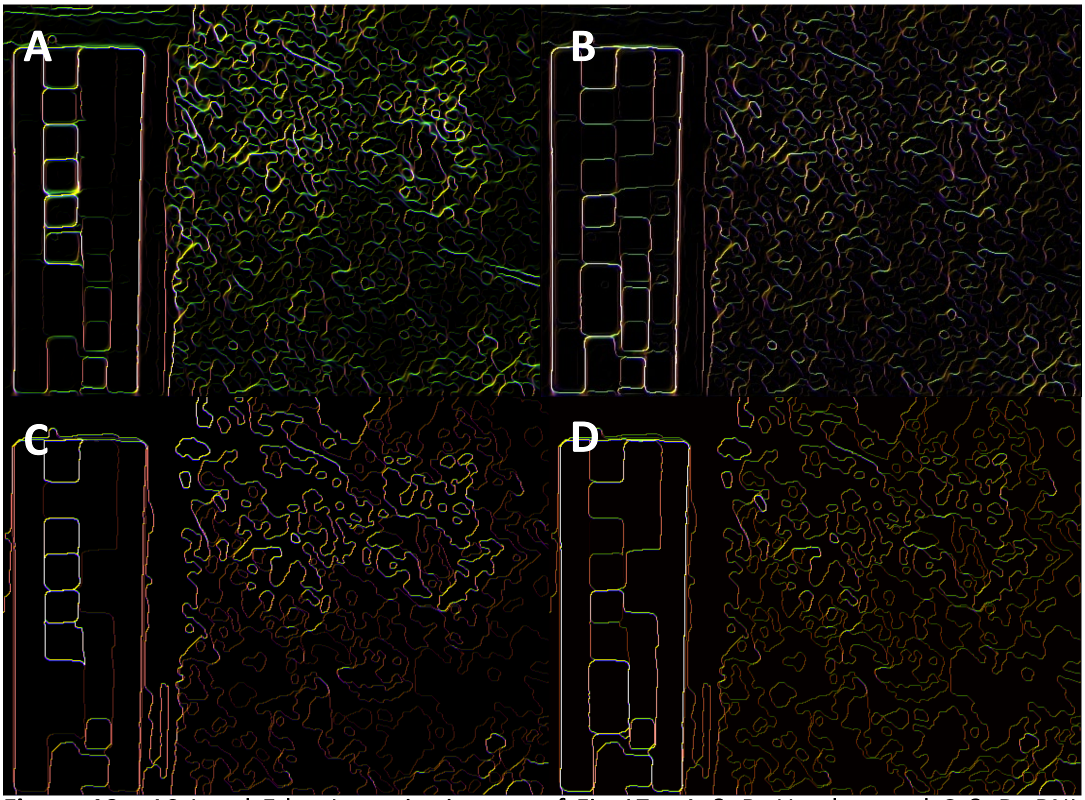

**Figure 19:** ΔS Local Edge Intensity images of Fig 17a. A & B: Un-clustered C & D: RNL Clustered. A & C show chromatic edge intensities B & D achromatic edge intensities. Note how the clustering essentially just removes the low intensity edges. The animal is not visible in either one of the images. Intensity images can be quantified using the LEIA parameters.

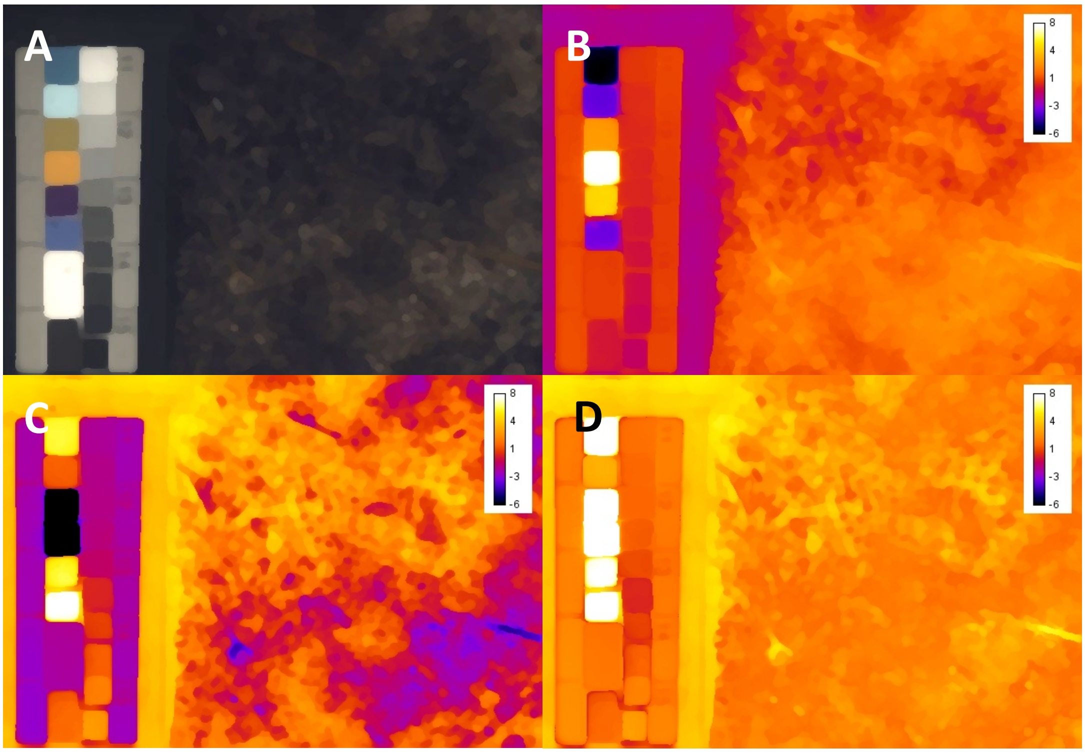

**Figure 20.** A: Un-clustered image at 10cm viewing distance using triggerfish visual acuity and spectral sensitivity. B: X-axis chromaticity image. C: Y-axis chromaticity image. D: Saturation image. The user may want to apply pattern analysis and imageJ image analysis tools to quantify these images.

**Deriving Animal + Background Parameters**

To interpret the colouration of our two animals (separate as well as in contrast of their visual backgrounds) we need to analyse them separately as regions of interest (ROIs) in each picture.

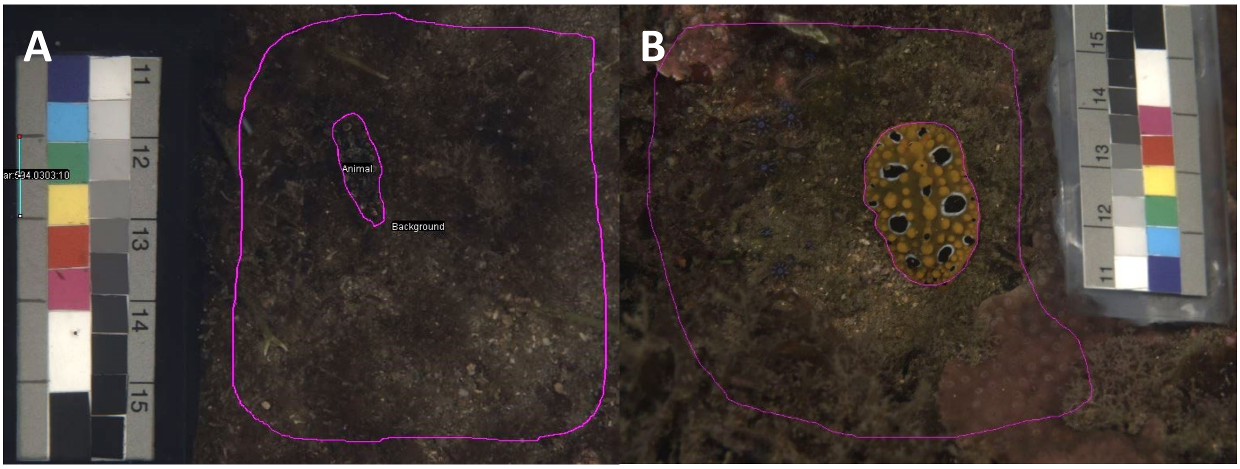

**Figure S11:** The Region of Interest (ROI) selection for the animal vs. background comparison. Please see the online user guide for details on ROI selection. Note: The area of the background can also be standardised, e.g. using a fix animal to background area ratio if that is desirable.

The selection of the ROIs can be made in an RGB reconstructed colour image which is often easier than using the grey scale intensity image layers of a multispectral image. The ROI manager in ImageJ remembers the ROI selection so you can select the multispectral image and click on the ROI which will then show up. Thus, the ROI selection can be made at any of the two following stages:

1. Right after creating or opening a multispectral image
2. After translating the multispectral image into cone catches

The prerequisite is that the size of the image is the same between these changes so the ROI remains the same. If you change the size of the image (i.e. after rescaling in the process of modelling visual acuity) and select the ROI it will not match anymore.

In this case we are analysing the animal separate from its background and thus (After modelling cone captures) we want to continue processing the ROIs separately from each other.

Because the ROIs are irregularly shaped we can’t use AcuityView 2.0 but instead must use the Gaussian filter to model spatial acuity of our triggerfish (See ‘AcuityView 2.0 & Gaussian Convolution Filter’ in this document). We choose a viewing distance of 10 cm and 30cm for both nudibranchs. Note that any kind of acuity modelling quickly turns the cryptic pattern into a uniform brown (Fig. S22). The MRA (Minimum resolvable angle) should be set at 5 pixels/MRA or above to avoid losing information (See ‘AcuityView 2.0 & Gaussian Convolution Filter’). Also note that we need to select a size standard prior to acuity modelling.

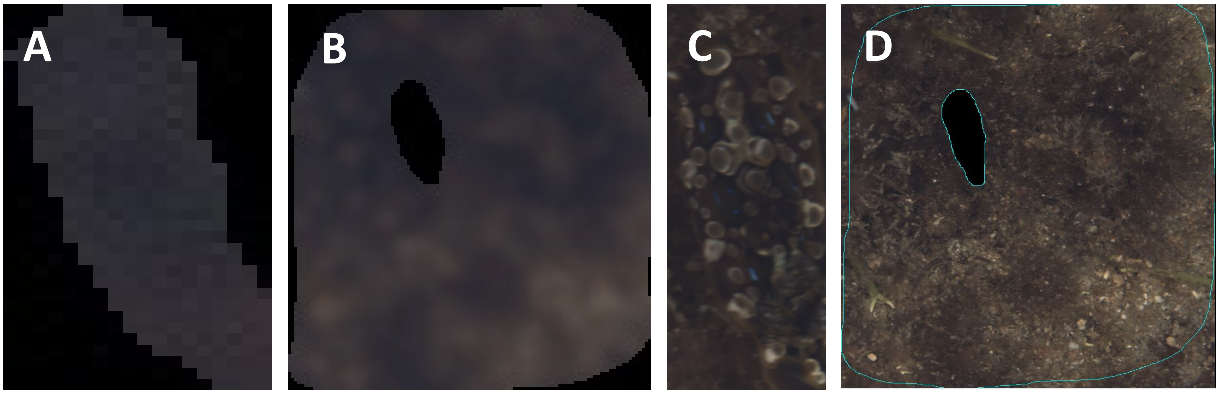

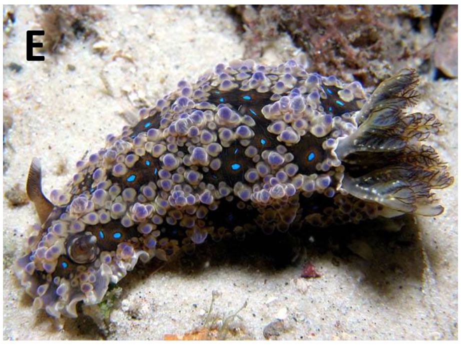

**Figure S22:** The ‘Animal’ ROI (A) and the ‘Background’ ROI (B) treated with the Gaussian acuity filter using a viewing distance of 30cm and a spatial acuity of 2.75 cycles/degree. The resolution is set to 5 pixels/MRA. As indicated in Fig. 20, the ‘cryptic’ nudibranch is likely not detectable at this distance by a triggerfish. If wee look at the unclustered ROIs (C & D) we can see that the animal actually sports iridescent blue spots, a common feature of many marine animals. E) picture of the nudibranch, *Dendrodoris krusensternii* (by Dave Harasti) in close up on a more contrasting background.

The ‘blurred’ ROIs then need to be treated with the RNL ranked filter to remove most of the blur. For this we choose the following settings: 5 repeats with a radius of 5 pixels (the size of our MRA) and a falloff of 3. We set the noise ratios at 0.05 (mw), 0.05 (lw), 0.07 (sw) and 0.05 (dbl) (Fig. S23 A & C).

As Fig. S23 suggests, even at 10cm it is likely that our cryptic nudibranch does not bear any discriminable patterning when considering our triggerfish observer. The background most likely consists of a few brownish hues into which the cryptic nudibranch blends nicely. This notion of background matching is supported when looking at the similarity of the clustered nudibranch colour and its surrounding background. The nudibranch has a saturation contrast (Distance from the achromatic point in the log-transformed RNL colour space) of less than 0.1 ΔS, a chromatic contrast (distance between the animal and its background in the log- transformed RNL colour space) of 0.58 ΔS and a luminance Michelson contrast of 0.02%. We can also quantify the chromatic background matching of this assumedly cryptic nudibranch using a RNL colour map (Fig. S24). This shows us that our animal is an almost perfect chromatic match to its background. The overlap measure provided by the colour map feature tells us that the animal ROI overlaps with 27.5% of the background ROI.

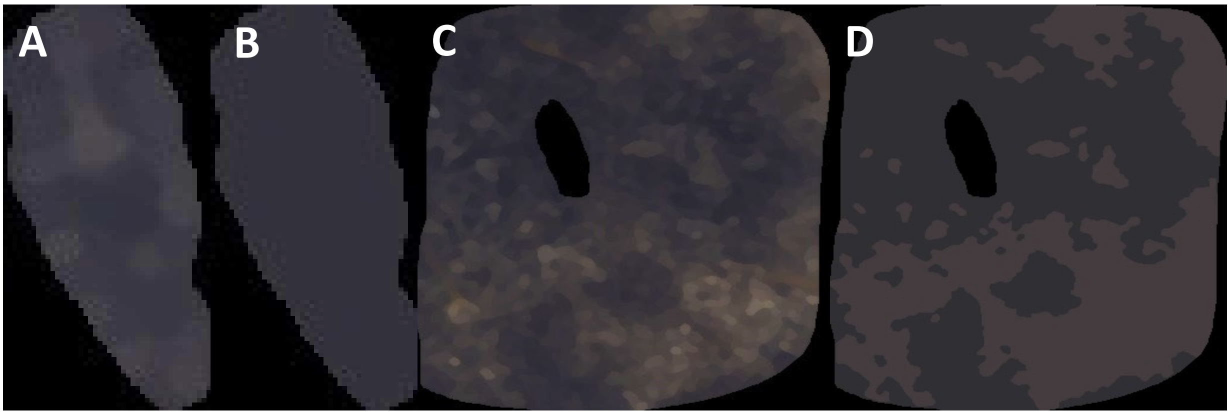

**Figure S23:** A) The RNL treated animal ROI from 10cm. B) The clustered animal ROI using a colour discrimination threshold of 3 ΔS and an achromatic discrimination threshold of 4 ΔS. Note that while patterning is visible in A the contrast is not sufficient to be picked up by the clustering, resulting in a single uniform cluster for the animal. C) The RNL treated background ROI from 10cm D) the clustered background using a colour discrimination threshold of 3 ΔS and an achromatic discrimination threshold of 4 ΔS. Note the difference to image S20c where no acuity modelling was applied.

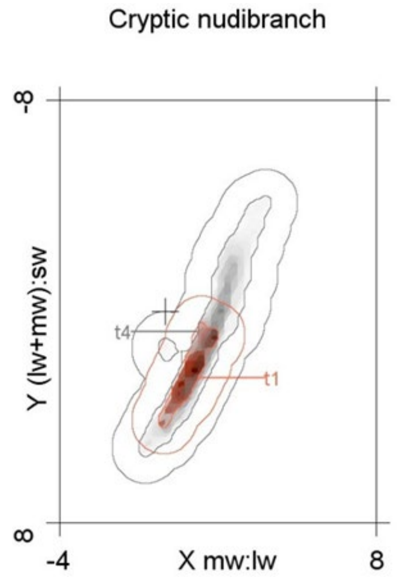

**Figure S24:** A colour map of our cryptic nudibranch (t1) and its visual background (t4). The darkness indicates the frequency of pixels in that area of the log-transformed RNL colour space. The border line around each cloud corresponds to 1 ΔS. The cross indicates the location of the achromatic point. We can easily see that the animal is an almost perfect subsample of its visual background.

Based on this outcome for the cryptic nudibranch we can see that there is no information on pattern to analyse at very close distances. It would therefore lead to the conclusion that the selective pressure that lead to the evolution of the blue spots in *Dendrodoris krusensternii* is probably due to an animal with better spatial resolution than what we assume to be the case for our triggerfish (2.75 cycles/degree). It could also be that the blue spots contribute to the additive blurring that leads to the observed background matching. While we could go and analyse the ROIs using an even smaller viewing distance or no acuity modelling at all (i.e. assuming human-like spatial acuity), we do not do this here.

We can now look at our assumedly 'aposematic' nudibranch, *Phyllidia ocellata*. We do know (Cheney et al. unpublished data) that this species is highly distasteful to potential predators, but we now want to investigate if it also shows signs of being 'conspicuous' i.e. displaying vivid visual contrast that helps predators to remember and detect the animal.

First, we model spatial acuity with a 10cm viewing distance for each the animal and its visual background, using the same settings as in the previous example (Fig. S25).

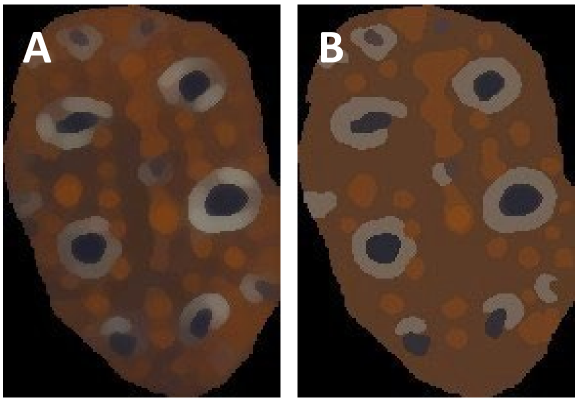

**Figure S25:** A) The RNL treated animal ROI from 10cm. B) The clustered animal ROI using a colour discrimination threshold of 3 ΔS and an achromatic discrimination threshold of 4 ΔS. C) The RNL treated background ROI from 10cm D) the clustered background using a colour discrimination threshold of 3 ΔS and an achromatic discrimination threshold of 4 ΔS. Note the similarity of the background to the cryptic nudibranch example.

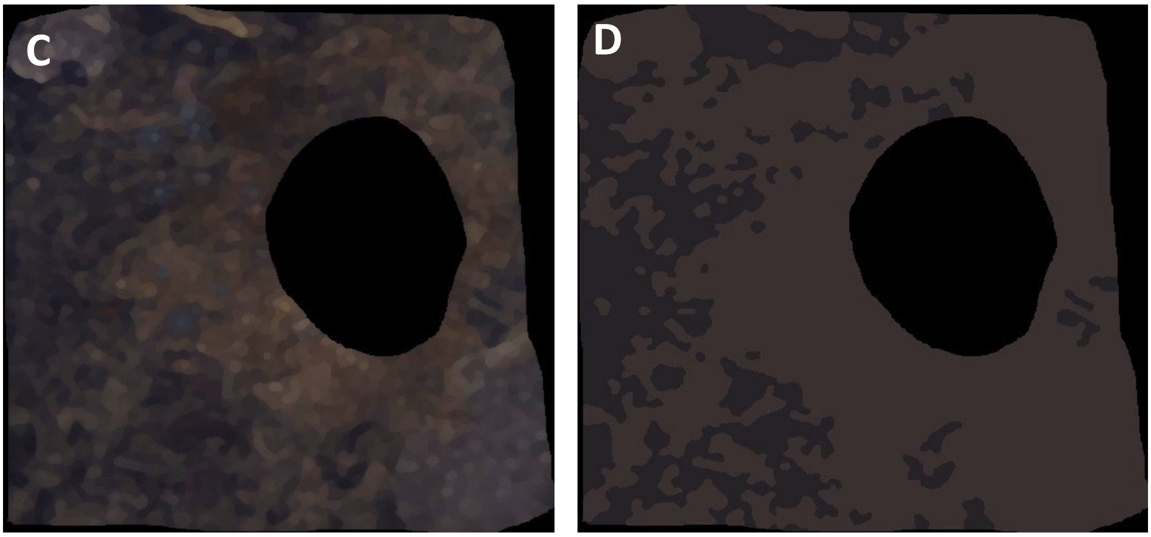

We can now use the clustered ROIs to run the pattern analyses on. This gives us all the pattern statistics of the adjacency analysis, the visual contrast analysis and the boundary strength analysis as well as descriptions of each colour pattern element in the ROIs. We can also use the RNL ranked filter treated images to estimate background matching using a colour map overlap analysis.

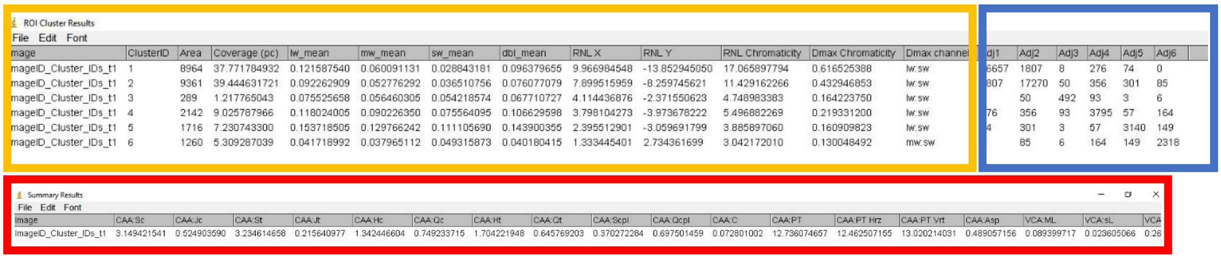

ROI Cluster Results

File Edit Font

| mage | ClusterID | Area | Coverage (pc) | lw_mean | mw_mean | sw_mean | dbl_mean | RNL X | RNL Y | RNL Chromaticity | Dmax Chromaticity | Dmax channel | dj1 | Adj2 | Adj3 | Adj4 | Adj5 | Adj6 |
|---|---|---|---|---|---|---|---|---|---|---|---|---|---|---|---|---|---|---|
| mageID_Cluster_IDs_t1 | 1 | 8964 | 37.771784932 | 0.121587540 | 0.060091131 | 0.028843181 | 0.096379655 | 9.966984548 | -13.852945050 | 17.065897794 | 0.616525388 | lw:sw | 6657 | 1807 | 8 | 276 | 74 | 0 |
| mageID_Cluster_IDs_t1 | 2 | 9361 | 39.444631721 | 0.092262909 | 0.052776292 | 0.036510756 | 0.076077079 | 7.899515959 | -8.259745621 | 11.429162266 | 0.432946853 | lw:sw | 807 | 17270 | 50 | 356 | 301 | 85 |
| mageID_Cluster_IDs_t1 | 3 | 289 | 1.217765043 | 0.075525658 | 0.056460305 | 0.054218574 | 0.067710727 | 4.114436876 | -2.371550623 | 4.748983383 | 0.164223750 | lw:sw | | 50 | 492 | 93 | 3 | 6 |
| mageID_Cluster_IDs_t1 | 4 | 2142 | 9.025787966 | 0.118024005 | 0.090226350 | 0.075564095 | 0.106629598 | 3.798104273 | -3.973678222 | 5.496882269 | 0.219331200 | lw:sw | 76 | 356 | 93 | 3795 | 57 | 164 |
| mageID_Cluster_IDs_t1 | 5 | 1716 | 7.230743300 | 0.153718505 | 0.129766242 | 0.111105690 | 0.143900355 | 2.395512901 | -3.059691799 | 3.885897060 | 0.160909823 | lw:sw | 4 | 301 | 3 | 57 | 3140 | 149 |
| mageID_Cluster_IDs_t1 | 6 | 1260 | 5.309287039 | 0.041718992 | 0.037965112 | 0.049315873 | 0.040180415 | 1.333445401 | 2.734361699 | 3.042172010 | 0.130048492 | mw:sw | | 85 | 6 | 164 | 149 | 2318 |

Summary Results

File Edit Font

| Image | CAA:Sc | CAA:Jc | CAA:St | CAA:Jt | CAA:Hc | CAA:Qc | CAA:Ht | CAA:Qt | CAA:Scpl | CAA:Qcpl | CAA:C | CAA:PT | CAA:PT Hrz | CAA:PT Vrt | CAA:Asp | VCA:ML | VCA:sL | VCA |
|---|---|---|---|---|---|---|---|---|---|---|---|---|---|---|---|---|---|---|
| ImageID_Cluster_IDs_t1 | 3.149421541 | 0.524903590 | 3.234614658 | 0.215640977 | 1.342446604 | 0.749233715 | 1.704221948 | 0.645769203 | 0.370272284 | 0.697501459 | 0.072801002 | 12.736074657 | 12.462507155 | 13.020214031 | 0.489057156 | 0.089399717 | 0.023605066 | 0.26 |

**Figure S26:** The output of the QCPA pattern analysis. The orange box contains information on each colour pattern element in the ROI. The blue box shows the adacency analysis (horizontal and vertical output is possible which triples the parameter output). The red box shows all pattern parameters.

We will not dive any deeper into the analysis of the pattern parameters at this stage. However, we would like to point out that various pattern parameters will indicate animal-background contrast (Fig S26) on top of several other measures indicating significant chromatic, achromatic and pattern contrast of our second animal against its background (e.g. Fig. S27). We will continue to elaborate worked examples and applications of specific tools in QCPA on the website (www.empiricalimaging.com) so please continue to check there on updates on worked examples, tutorial videos, user guides as well as to get in touch with the wider user community in the forum.

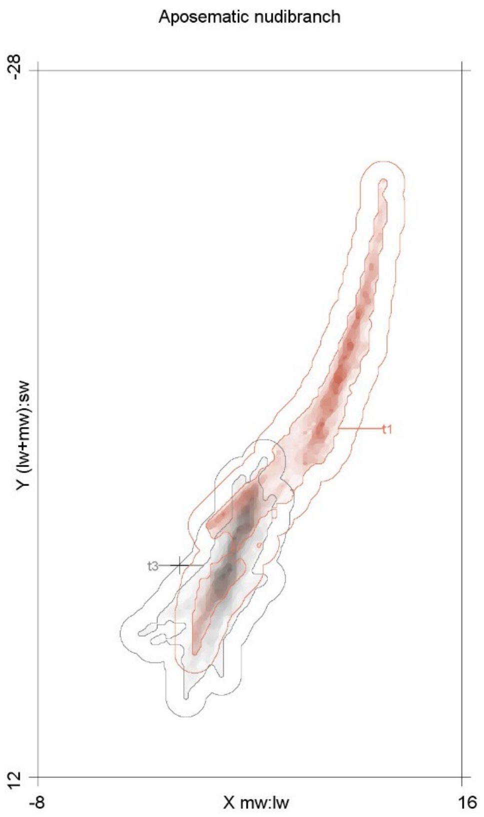

**Figure S27:** A colour map of our cryptic nudibranch (t1) and its visual background (t3). The darkness indicates the frequency of pixels in that area of the log-transformed RNL colour space. The border line around each cloud corresponds to 1 ΔS. The cross indicates the location of the achromatic point. In this case the animal shares a mere 8% overlap with its visual background.
